# Distinct cell wall molecular architecture of dimorphic *Talaromyces marneffei* cells revealed by solid-state NMR spectroscopy

**DOI:** 10.1101/2025.09.26.678783

**Authors:** Qinghui Cheng, Xinyue Xu, Siming Liao, Yaxin Chen, Hao Liang, Jian Wang, Fang Wang, Sanqi An

**Author notes:** These authors contributed equally: Qinghui Cheng and Xinyue Xu. Corresponding authors: Qinghui Cheng, Jian Wang, Fang Wang, and Sanqi An.

## Abstract

*Talaromyces marneffei*, causing systemic infections in immunocompromised patients ranging from HIV/AIDS individuals to cancer and transplant recipients, is an increasingly urgent global pathogen. However, the fungus remains underrecognized despite the systemic infection disease talaromycosis caused by this pathogen is associated with high mortality rates. Its pathogenicity depends on a temperature-triggered shift from saprophytic mold (25 °C) to pathogenic yeast (37 °C), and the two growth forms display distinct sensitivity to antifungal drugs, which processes involve extensive cell wall structure and components remodeling. To dissect these processes, we use solid-state nuclear magnetic resonance (ssNMR) and other techniques to show that *T. marneffei* yeast and hyphal cells have distinct cell wall thickness and hydrophobicity, and different assembly of mobile and rigid polymers within the *T. marneffei* cell wall. The yeast wall was 2.3 times thicker and more hydrated. ssNMR revealed a rigid core of β-1,3-glucans, chitin and chitosan, with β-1,3-glucan rising from 57% in mold to 72% in yeast. Both forms showed tight polysaccharide packing, but only mold exhibited lysine-containing protein interactions with chitin and chitosan. These insights not only map the structural basis of host temperature adaptation and also inform targeted antifungal design in future.

## INTRODUCTION

*Talaromyces marneffei* (formerly *Penicillium marneffei*) is a dimorphic fungus endemic to Southeast Asia, Northeastern India, and Southern China.(*1, 2*) Uniquely among *Penicillium* species, it causes systemic mycosis in humans.(*1, 3*) In contrast, other *Penicillium* species (e.g., *P. chrysogenum*, *P. roqueforti*, *P. cluniae*, *P. digitatum*, *P. notatum*, *P. stipitatus*) rarely cause human infection, typically only under specific occupational or environmental exposures, *T. marneffei* poses a severe threat, particularly to immunocompromised individuals.(*4–9*) *T. marneffei* presents a significant clinical burden, particularly in immunocompromised individuals. Historically associated with HIV/AIDS, disseminated talaromycosis predominantly affects patients with severe CD4⁺ T-cell depletion, causing systemic infection characterized by fever, weight loss, anemia, and distinctive skin lesions following its transition from environmental mold to pathogenic yeast.(*10–12*) The mortality rate remains alarmingly high, ranging from 24% to 33% even with antifungal therapy.(*13*) By mid-2022, over 288,000 cases had been reported across 34 countries, representing a pooled prevalence of 3.6% among people living with HIV.(*14–16*) Notably, advancements in cancer therapy and organ transplantation are elevating the incidence of *T. marneffei* infections in HIV-uninfected individuals, putting those individuals with no previously recognized immune disorders at risk.(*17*) Despite being ranked the world’s second most feared fungal disease with profound clinical impact, talaromycosis remains globally overlooked regarding diagnostic and therapeutic strategies.(*18, 19*)

Standard antifungal therapy relies on four main classes: polyenes, azoles, echinocandins, and the pyrimidine analog 5-flucytosine.(*20*) However, treatment failure persists as a major clinical challenge due to complex fungal adaptive responses.(*21, 22*) Contributing factors include host immune compromise, pharmacological limitations such as pharmacokinetic variability, suboptimal drug-target engagement, and adverse drug interactions, alongside intrinsic fungal survival mechanisms including morphological plasticity, drug tolerance, and resistance pathways.(*23*) A likely key contributor to these challenges is the unique composition and dynamic remodeling of the *T. marneffei* cell wall. Unlike many fungal pathogens, *T. marneffei* undergoes an essential dimorphic transition between mold and yeast forms for virulence(*24, 25*), necessitating extensive, dynamic reorganization of its β-glucan- and chitin-rich wall. This continual remodeling likely compromises standard therapies (e.g., echinocandins, azoles) by altering drug-accessible targets.(*26*),(*27, 28*) Despite this challenge, the fungal-specific presence of these polysaccharides (absent in human cells) makes them ideal therapeutic targets. Therefore, a deeper understanding of *T. marneffei* cell wall composition and adaptive remodeling during infection is crucial for identifying novel therapeutic targets.

Dimorphic transition involves significant cell wall reorganization characterized by compositional shifts that enhance stress resistance and immune evasion.(*29*) However, the molecular mechanisms driving these changes, such as temperature-dependent enzyme activity, polysaccharide cross-linking, and protein-carbohydrate interactions, remain poorly characterized. For instance, β-1,3-glucans, the primary target of echinocandins, are often shielded by outer glucan or galactomannan layers in fungi, reducing drug efficacy;(*30, 31*) preliminary evidence suggests β-1,3-glucans are enriched in both *T. marneffei* morphotypes, yet their spatial distribution, interaction networks, and hydration states are not fully mapped.(*32*) Furthermore, the activity of multiple chitin synthases (CHS) and chitin deacetylases (CDAs) generates heterogeneous chitin and chitosan polymorphs, contributing to complex wall architecture.(*33, 34*) Elucidating the molecular-level structure of these components is therefore critical for understanding how *T. marneffei* cell wall adaptations support pathogenicity and antifungal resistance.

In this study, uniformly ¹³C/¹⁵N-labeled fungal samples were cultured at 25 °C (mold form) and 37 °C (yeast form) to facilitate a comprehensive analysis of polysaccharide composition, dynamics, and intermolecular interactions within the cell walls of *T. marneffei*.(*35*) Complementary TEM imaging illustrated temperature-dependent alterations in cell wall thickness, whereas ssNMR results demonstrated detailed, site-specific insights into the rigid core structures and more mobile outer cell wall layers.(*36–38*) This work indicate that β-1,3-glucans, chitin, and chitosan constitute the rigid core of *T. marneffei* cell wall, while galactomannan and additional polysaccharides primarily form the mobile matrix. Notably, the yeast demonstrated enhanced intermolecular polysaccharide interactions and increased hydration levels, while still preserving conserved spatial contacts between β-1,3-glucan/chitosan and proteins. A 2.3-fold thickening cell wall and an elevated β-1,3-glucan content (rising from 57% in mold to 72% in yeast) are detected from TEM analysis in the yeast cell wall compared to the mold form. High-resolution ssNMR of intact cells identified β-1,3-glucans, chitin, and chitosan as predominant components of the rigid core. Both morphological forms exhibited intermolecular polysaccharide packing, indicative of compact cell wall assembly; however, differences in polysaccharide-protein interactions were observed. Specifically, lysine interactions with chitin/chitosan is exclusive to the mold form, while tryptophan interaction with β-1,3-glucan and chitosan are detected in both phases. Therefore, these differences in dimorphic *T. marneffei* cells induced by growth temperatures have provided heretofore unavailable molecular insights into the pathogenicity and drug resistance of the yeast form. These findings not only shed light on the structural basis of *T. marneffei*’s dimorphism but also identify potential vulnerabilities for targeted antifungal development. By bridging the gap between molecular-level analysis and phenotypic adaptation of fungus in response to temperatures, this work highlights the indispensability of ssNMR in advancing fungal cell wall research and provides a structural basis for combating drug-resistant pathogens. Our findings provide critical insights into the molecular underpinnings that govern cell wall remodeling in *T. marneffei*, potentially revealing novel regulatory pathways that could be targeted for therapeutic intervention.

## MATERIALS AND METHODS

### Preparation of Uniformly ^13^C/^15^N-labeled *T. marneffei*

The fungal strain *T. marneffei* (ATCC18224) was cultured at 25 °C and 37 °C in growth media enriched with ¹³C/¹⁵N isotopes using the following protocol (**Supplementary figure 1**). First, fungal conidia were grown on a pre-sterilized solid medium prepared with potato dextrose agar (Sigma-Aldrich, USA) for 10 days at 27 °C. Subsequently, the collected conidia were inoculated into a sterilized 250 mL flask containing 100 mL of pre-sterilized ¹³C/¹⁵N-enriched growth medium (U-^13^C_6_-labeled D-glucose, CAS#110187-42-3, Cambridge Isotope Laboratories, Inc, USA; ^15^N_2_-ammonium sulfate, CAS#43086-58-4, Cambridge Isotope Laboratories, Inc, USA; Yeast nitrogen base w/o amino acids, catalog#DF0919-15-3). The flasks (Prod#CFT021500, Shanghai Jinpan Biotech Co., Ltd) were placed in shakers set to 200 revolutions per minute (RPM) and incubated at 25 °C and 37 °C, respectively. After 14 days of cultivation (**Supplementary figure 2**), the fungal pellets were harvested by centrifugation (Thermo Scientific Sorvall ST 40R, Cat# 75004524, Thermo fisher Scientific Inc., Germany) at 10,000 RPM for 30 minutes at 4 °C. The pellets were then washed three times with phosphate-buffered saline (PBS, CAS#P1020-500, Beijing Solarbio Science & Technology Co., Ltd) to remove water-soluble metabolites and residual growth medium components. Finally, the collected fungal pellets were stored at −80 °C for further usage.

### Measurement of Cell wall thickness and morphological imaging

Both the fungal samples were initially fixed using a solution containing 2.5% glutaradehyde (CAS#NC1536477, TED PELLA INC., CA, United States) at 4 °C for 24 hours, followed by post-fxiation in 1% osmium tetroxide (Prod#18456, TED PELLA INC., CA, United States) with PBS (CAS#P1020-500, Beijing Solarbio Science & Technology Co., Ltd) rinses between steps. The samples were dehydrated for 10 min using ethanol (CAS# 64-17-5, Sinopharm Chemical Reagent Co., Ltd) at incremental concentrations (30%, 50%), followed by staining with uranyl acetate (CAS#541-09-3, Electron Microscopy Sciences, PA, United States) in 70% ethanol for 3 hours. Subsequent steps included sequential immersion in 80% ethanol (10 min), 95% ethanol (15 min), and two rounds of 100% ethanol (50 min each), after which the samples were transitioned via propylene oxide (CAS#15448-47-2, Meryer (Shanghai) Biochemical Technology Co., Ltd). Next, the specimens were infiltrated with a 1:1 mixture of propylene oxide and Eponate 12 resin (Cat#NC1261266, TED PELLA INC, CA, United States), embedded in pure resin, and polymerized at 45 °C for 12 hours followed by 72 °C for 24 hours. Ultrathin sections were achieved by a LEICA UC-7 Ultramicrotome, mounted on a copper grid, and then stained with lead citrate solutioin (Prod#. 19312, TED PELLA INC, CA, United States). Imaging was carried out on a Hitachi HT7800 TEM equipped with a Morada G3 camera, at magnifications of 5,000x-50,000x to observe the perpendicular cross section of both fungal samples and the representative images were displayed in **Supplementary figure 2**. Cell wall thickness for both samples were obtained using ImageJ software 1.54m(*39*) and the statistical analysis for the cell wall thickness was done by t-test for both samples (**Supplementary table 1**).

### Solid-state NMR experiments

All the solid-state NMR experiments were conducted on a Bruker Avance 700 MHz (16.4 T) spectrometer using a 3.2 mm probe. All the experiments were acquired under 15 kHz MAS at 277 K. ^13^C chemical shifts were calibrated externally using the methylene carbon of the adamantane, set at 38.48 ppm relative to the tetramethylsilane (TMS) scale. Unless otherwise specified, typical radiofrequency field strengths were employed: 77.4 kHz for proton decoupling, 63.1 kHz for proton cross-polarization (CP) and hard pulses, 63.1 kHz for ^13^C. All the parameters were listed in **Supplementary table 2**.

1D ^13^C spectra were conducted using three different polarization methods to selectively detect the signals from molecules with distinct dynamics. Specifically, a 1D dipolar-mediated ^1^H-^13^C cross-polarization (CP) experiment was employed to preferentially detect the signals of the rigid components using a contact time of 0.5 ms between ^1^H-^13^C. Additionally, a 1D ^13^C direct polarization (DP) technique with a short recycle delay of 2 s was used to selectively detect mobile molecules exhibiting fast ^13^C-T_1_ relaxation. Finally, a 1D ^13^C direct polarization (DP) experiment with a long recycle delay of 35 s was performed to observe all molecules, ensuring a comprehensive detection of all ^13^C-containing cellular components by allowing sufficient time for full relaxation.

To facilitate the resonance assignment of the specific chemical moieties and carbon sites within individual type of molecules which corresponds to each NMR signal, three types of solid-state NMR experiments were measured: (1) 2D refocused ^13^C J-INADEQUATE spectra with DP and short recycle delays of 2 s for selective detection of the molecules located in the mobile domain, i.e., in the outer layer of the cell walls;(*40*) (2) ^13^C CP J-INADEQUATE for the identification of the molecules within rigid domain, located inside the cell wall; (3) 2D ^13^C-^13^C spectra with 53-ms mixing CORD.(*41*) The first and second were utilized to track the carbon connections within each individual molecule while the third method is for the intramolecular cross peaks detection of rigid molecules by using 0.5 ms CP contact time between ^1^H and ^13^C.

The relative abundance of carbohydrates was estimated based on the peak volumes of well resolved peaks identified from the 2D ^13^C-^13^C correlation spectra with 53-ms CORD mixing and 2D ^13^C DP J-INADEQAUTE spectra. To be specific, the peak volumes of each type of carbohydrates were integrated on topspin software and then used for the estimation of the molar %age of each type of carbohydrates by normalizing the sum of peak volumes with their corresponding counts.(*42*) The equations for the calculations of relative abundance (*RA*) and error are listed as follows: 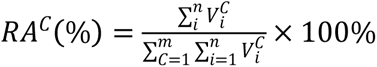 and 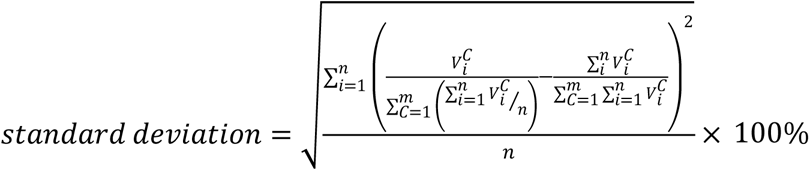, where RA^C^ (%) denotes the relative abundance (RA) of component C. 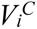 is the volume of the ith peak for component C. n represents the total number of peaks considered for component C, and m is the number of identified components included in the calculation.

To elucidate the intermolecular interactions among carbohydrate polymers, as well as between carbohydrates and proteins, 2D ^13^C-^13^C proton-driven spin diffusion (PDSD) experiments were carried out on fungal samples cultivated at 25 °C (molds) and 37 °C (yeasts), using a very long mixing time of 1.0 s.(*43, 44*) 236 intermolecular interactions were identified for the molds (25 °C), while 261 intermolecular interactions were detected for the yeasts (37 °C), and details were listed in **Supplementary table 13**.

To evaluate the water accessibility of carbohydrates, we initially conducted ^1^D ^1^H experiments to establish a T_2_ filter (1.0 s x 2), which reduced the ^1^H signals to 90% for molds and 88% for the yeasts.(*45, 46*) This T_2_ filter was then applied in 1D water-edited ^13^C-detection experiments, effectively suppressing 97% of the polysaccharide signals. Following this, the ^1^H magnetization from water was transferred to polysaccharides during a 4-ms ^1^H mixing period, and the magnetization was subsequently transferred from ^1^H to ^13^C through ^1^H-^13^C cross-polarization (CP) for high-resolution ^13^C detection. A 50-ms ^13^C-^13^C DARR mixing period was incorporated into the water-edited experiment to selectively detect well-hydrated molecules, while the control version of 2D ^13^C-^13^C DARR spectrum displayed full signal intensities. The relative intensity ratios (S/S_0_) between the water-edited spectrum (S) and the control spectrum (S_0_) were calculated for all identifiable carbon sites, enabling a comparison of the relative hydration levels among different polysaccharides and between the two samples.

The dynamics of the rigid carbohydrate polymers were examined using the 1D ^13^C Torchia-CP experiment, with the z-filter duration varying from 0.1 ms to 9 s.(*47*) The intensity decay for each resolved peak was measured as the z-filter time increased, and the pre-normalized data were fitted to a single exponential equation to calculate the ^13^C-T_1_ relaxation time constants.

## RESULTS

### b-1,3-glucans, chitins, and chitosan form the rigid core of *T. marneffei* cell wall

To characterize the initial morphological difference of yeast and hyphal cells of *T. marneffei*, we firstly applied transmission electron microscopy (TEM) to compare the cell wall structure. Cross-sectional imaging of cells cultured at 25 °C and 37 °C demonstrated a marked increase of cell wall thickness at yeast cell (**Fig. 1a**). Quantitative assessment using ImageJ software 1.54d,(*39*) revealed a 2.3-fold expansion in thickness from 67 ± 9 nm (mold form) to 152 ± 21 nm (yeast form) (**Fig. 1b, Supplementary table 1**). This trend aligns with observations in *Rhizopus delemar*, where cell wall thickness doubled following treatment with the chitin synthase inhibitor nikkomycin illustrating the plasticity of the fungal cell wall under stress.(*33*) In another similar fashion, *Aspergillus sydowii* adapts to hypersaline conditions by thickening its cell wall.(*48*) Together, these examples demonstrate how fungi actively remodel their cell walls in response to challenging environments, revealing a common adaptive strategy across different species.

**Fig. 1.**
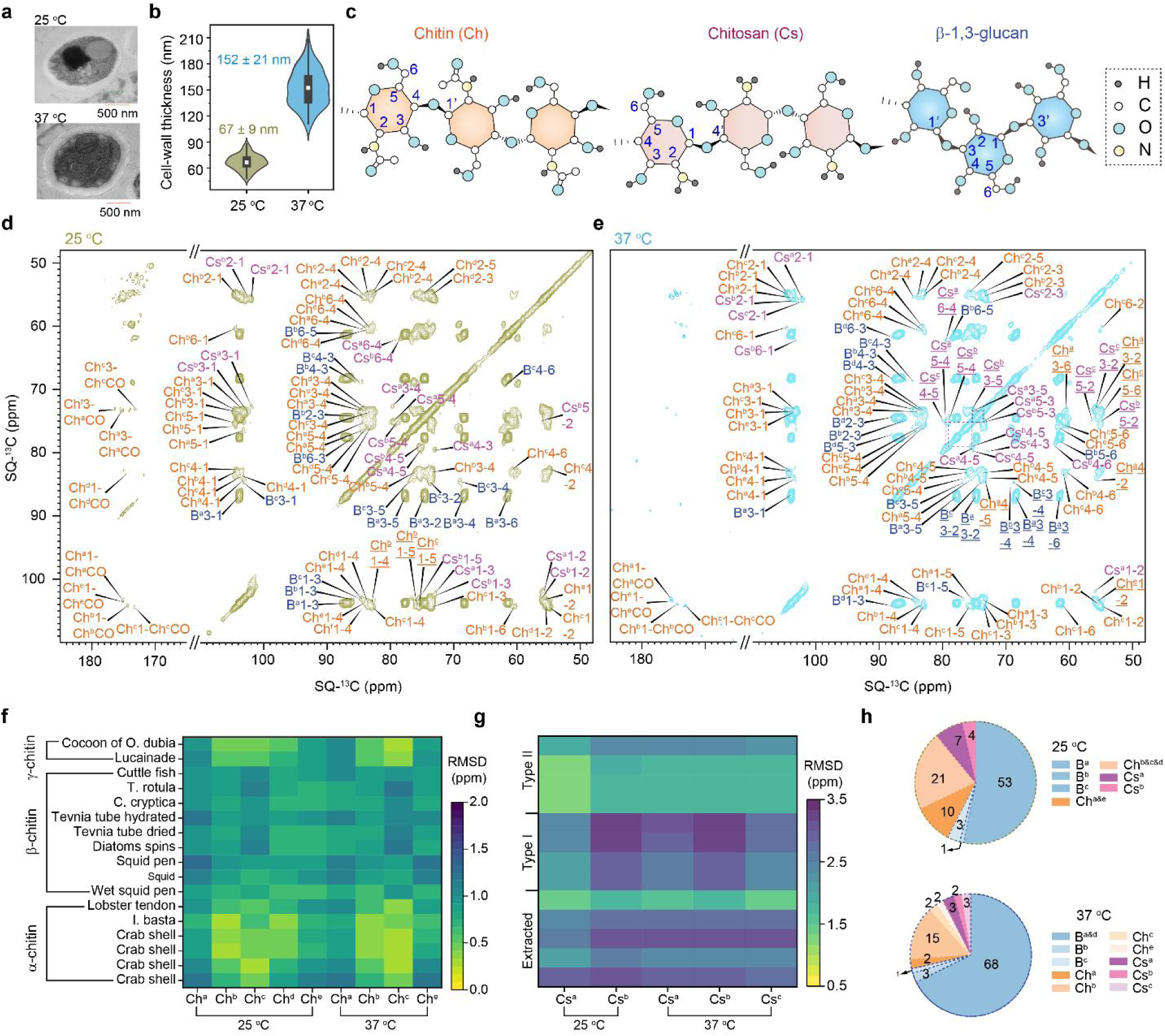
b-1,3-glucan together with chitin and chitosan constitute the rigid core. (**a**) Representative TEM images of both samples (top: 25 °C; bottom: 37 °C). (**b**) Violin plots of the cell wall thickness of *T. marneffei* cultivated at 25 and 37 °C. Six cells were analyzed for each sample by TEM imaging and data analysis was from 90 measurements. (**c**) Molecular structures of identified polysaccharides in fungal cell wall, as shown in different colors. (**d-e**) Rigid carbohydrate of hyphal (d) and yeast (e) cells detected by 2D ^13^C-^13^C correlation spectra using 53-ms CORD, where the polysaccharides identified are abbreviated and color-coded: Ch for chitin (orange), Cs for chitosan (pink), and B for b-1,3-glucan (blue). (**f-g**) ^13^C chemical shifts-based RMSD heatmaps for the identified chitins (f) and chitosan (g) with the literature data. (**h**) Estimation of relative abundances of the identified polysaccharides in hyphal (top panel) and yeast (bottom panel) cells. Data was calculated by the integrals of well-resolved cross peaks identified for the individual ones.

To further identify the molecular basis of *T. marneffei*’s temperature-dependent cell wall remodeling, we firstly focus on the rigid polysaccharides, which is critical for mechanical stability of the cell wall. Prior to this analysis, we emphasize that chitin, chitosan, and b-1,3-glucan are well-documented structural components of fungal cell walls, forming their rigid scaffold (**Fig. 1c**). To confirm the presence of these carbohydrates, we employed two-dimensional ^13^C-^13^C correlation spectroscopy with 53-ms CORD mixing. This technique selectively detects molecules exhibiting strong ^13^C-^1^H dipolar couplings, a hallmark of rigid polysaccharides. Spectra acquired from mold-form cells (25°C) and yeast-form cells (37°C) are shown in **Fig. 1d** and **1e**, respectively. Major signals are from b-1,3-glucan, chitin and its deacetylated forms, chitosan. The unique downfield ^13^C chemical shift of 85 ppm at the linkage site of carbon 3 (C3) indicates the signals from 1,3-glucans, and the C1 chemical shifts were found to 103.6 ppm typical for the existence of the b-1,3-glucans. Peak multiplicities of b-1,3-glucans were observed for both samples, 5 types for the molds, and 4 types for the yeasts. For the assignment of chitin, the intramolecular correlations between carbonyls (176-170 ppm) and C1 (102-104 ppm) and between methyl groups (∼20-25 ppm) and carbonyls suggest these types of polysaccharides are acetylated. Then we also observed C1 peaks are correlated with C2 peaks around 55 ppm, together with the C4 resonance peaks present around 82-85 ppm which further confirmed that these polysaccharides belong to chitin. Identification of the chitosan is relatively simple. The correlation peaks between C1 (97-103 ppm) and C2 (∼55 ppm) were observed while none of correlation peaks between these carbon sites and carbon sites resonating either 170-176 ppm or 20-25 ppm and combined the C4 peaks resonating at 77-80 ppm, typically suggesting the chitosan co-existed with the above polysaccharides to construct the rigid core. Subtypes of chitin and chitosan were also identified, 5 types of chitins and 4 types of chitosan were found in hyphal cell walls, but one type of chitin disappeared when cultivating at 37 °C (yeast). On one hand, the diversity of resonance patterns observed in the rigid carbohydrates is likely due to structural heterogeneity. On the other hand, the presence of multiple carbohydrate subtypes may also be attributed to carbohydrates with chemical similarities being located in different microenvironments within the rigid core, resulting in significant magnetic inequivalence.(*49*) We also employed the through-bond 2D ^13^C refocused CP-INADEQUATE experiments (**Supplementary figure 7**) to unambiguously track the carbon connectivity within each carbohydrate unit, which complementarily confirmed the analytical results by 2D ^13^C-^13^C correlational spectra with 53-ms CORD mixing. All the ^13^C chemical shifts of rigid carbohydrates were listed in **Supplementary table 9**.

Futhermore, analytical results revealed the presence of multiple chitin and chitosan types in the cell walls of both molds and yeast, with their structural polymorphism directly linked to diverse biological functions. Characterizing the conformations of these polymers in native cellular environments is inherently challenging. To address this, researchers frequently use ¹³C NMR chemical shifts to infer structural conformations by comparing deviations from reference biomolecules such as cellulose and amyloid fibrils.(*34*) Following this methodology, we analyzed the chitin and chitosan in *T. marneffei* cell walls by calculating the root mean square deviations (RMSD) between their assigned ¹³C chemical shifts and the published datasets summarized in a paper by Fernando et al.(*34*)

Our results (**Fig. 1f**) indicate that chitin structures in *T. marneffei* predominantly adopt comforamtions resembling a-chitin, with polymers stabilized by intra- and inter-chain hydrogen bonds to form a rigid structural core. However, yeast-phase cells cultured at 37 °C exhibited type-c chitin showing closer structural alignment to g-chitin, a hybrid form combining a- and b-chitin conformations, as observed in *Orychophora dubia* cocoons and *Lucaina* species.(*34*) For chitosan (**Fig. 1g**), comparative analysis with 13 literature datasets revealed structural similarities between mold-phase chitosan and Type II chitosan, indicative of a dynamic conformational state.(*34*) In contrast, yeast-form chitosan exhibited smaller RMSD values relative to specific extracted chitosan types, suggesting higher crystallinity within yeast cell walls.(*50, 51*) These variations align with transcriptomic findings that identified distinct gene expression patterns during phase transitions between mold and yeast forms of *T. marneffei*(*52*). These transcriptional shifts likely drive condition-specific protein expression profiles, including altered activity of enzymes such as chitin synthases and chitin deacetylases (CDAs). Such enzymatic changes may induce conformational variations in both chitin and its deacetylated derivative, chitosan, particularly in yeast. The observed structural modifications in chitin and chitosan across phases suggest a mechanism by which *T. marneffei* enhances environmental adaptability during host infection or stress responses. Notably, temperature-dependent variations in chitosan structure highlight the fungus’s ability to modulate chitosan synthase pathways in response to environmental stressors. These findings underscore the critical role of biopolymer structural plasticity in fungal adaptation and survival strategies.

For the compositional analysis of the rigid polysaccharides, we tentatively estimated the peak volumes of the well-defined peaks corresponding to each carbohydrate, as identified by high-resolution 2D ^13^C-^13^C correlation spectra obtained using 53-ms CORD mixing (**Fig. 1h**). The analysis demonstrated that β-1,3-glucan serves as the main component of the rigid core, accounting for 57 ± 5% of the rigid domain in the hyphal cell wall and increasing to 72 ± 8% in the rigid domain of the yeast cell wall. In contrast, the relative proportion of chitin decreased from 31 ± 2% in the rigid cell wall of the mycelia to 21 ± 4% in the rigid cell wall of the yeast (**Supplementary table 4**). A comparable reduction was observed for its deacetylated form, chitosan, which declined from 11 ± 3% in the rigid domain of the hyphal cell wall to 8 ± 2% in the rigid domain of the yeast cell wall. Notably, the cultivation temperature had no impact on the degree of deacetylation (DD) during the conversion of chitin to chitosan, which remained approximately 40% (calculated as the molar %age of chitosan divided by the sum of the molar %ages of chitosan and chitin). Chitin deacetylation is initiated by chitin deacetylases (CDAs), which remove acetyl groups from chitin molecules, producing chitosan through a consistent and high-quality deacetylation process.(*53*) Fungal CDAs are typically closely associated with chitin synthase, enabling the rapid deacetylation of newly synthesized chitin before it matures and crystallizes. Previous studies have indicated that CDAs exhibit high thermal stability, with optimal activity temperatures ranging from 30 to 60 °C.(*54*) This characteristic of CDAs likely results in a consistent degree of deacetylation (DD) in response to cultivation temperature. However, further evidence is needed to fully clarify this phenomenon.

### Diverse polysaccharides within the mobile domain

In addition to the rigid polysaccharides of the cell wall, certain carbohydrates are in the outer layers, where they exhibit relatively higher mobility. To investigate the biomolecules within the mobile domain, 1D ^13^C direct polarization (DP) with a 2-s recycle delay was used, targeting molecules with fast ^13^C **-**T_1_ relaxation. The NMR signals primarily arise from the polysaccharides, with contributions from proteins and lipids as well. The results are shown in **Supplementary figure 4**, where each molecule is highlighted in its respective region of the spectra. The mobile carbohydrates were estimated to comprise 51% of the whole cell wall of mycelia and 44% in that of yeast, respectively, based on the integrals of the corresponding chemical shift regions (**Supplementary table 3**). However, severe peak overlap in the 1D ^13^C spectra prevents accurate identification of individual carbohydrates, as shown in **Supplementary figure 5**. To resolve this, we combined short-recycle-delay ^13^C DP with 2D ¹³C DP J-refocused INADEQUATE spectra (**Fig. 2a** and **Supplementary figure 5**), enabling detailed assignments of major polysaccharides.

**Fig. 2.**
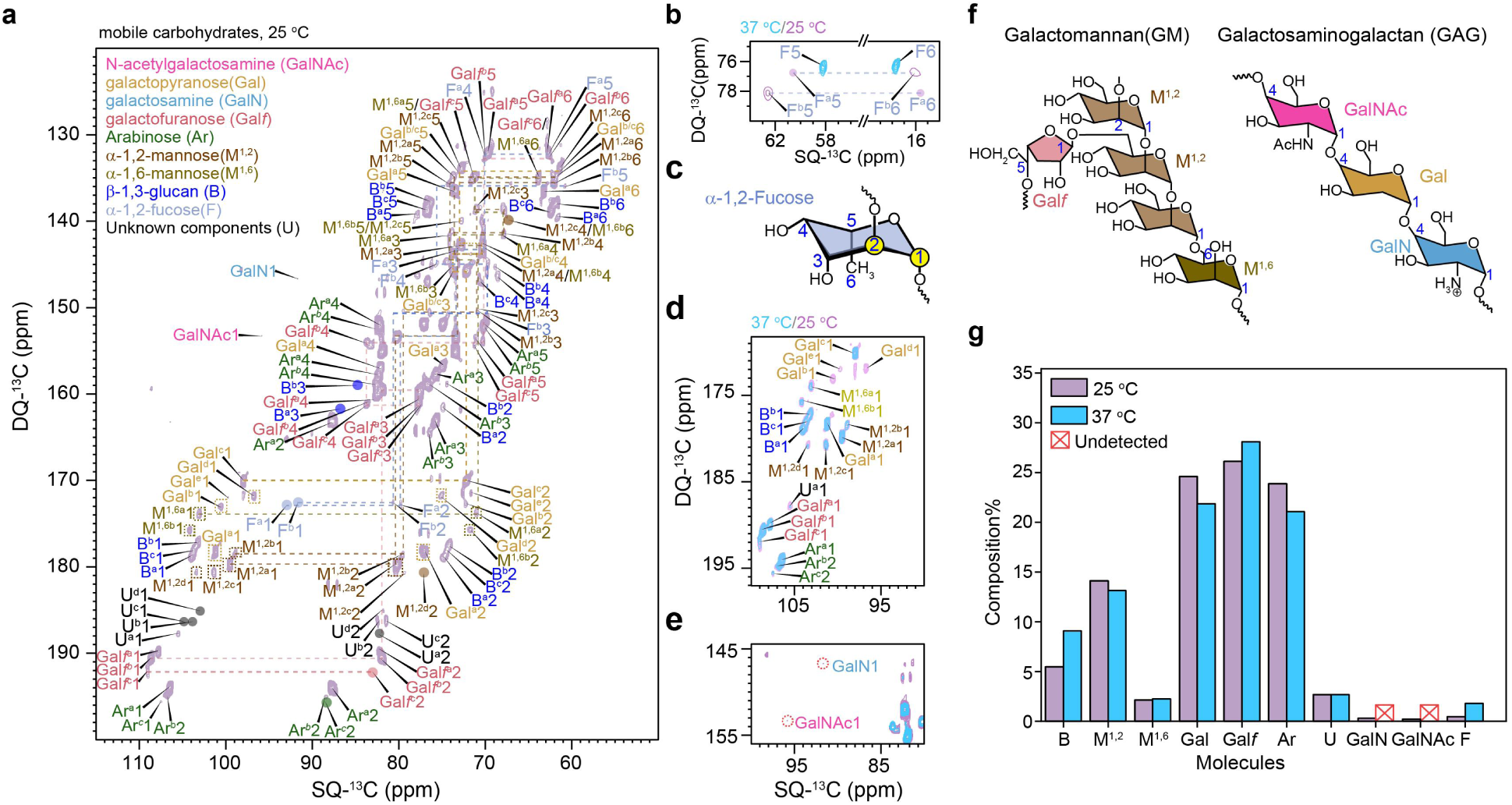
Mobile carbohydrates identified within cell walls. (**a**) Mobile carbohydrates were identified through 2D ^13^C DP J-refocused INADEQUATE spectra for hyphal cell wall (25 °C). Key monosaccharides are listed with full names and standardized NMR abbreviations; mobile monosaccharides are color-coded for clarity. (**b**) Evidence for confirming the existence of fucoses within both molds (purple) and yeast (cyan blue) cell walls. (**c**)Proposed glycosidic linkages of fucose. (**d**) The C1 region of the hyphal cell wall (25 °C in purple) overlaps with the corresponding region in yeast cell walls (37 °C in cyan blue). (**e**) Confirming the absence or negligible presence of galactosamine (GalN) and *N*-acetylgalactosamine (GalNAc) in yeast cell walls. (**f**) Illustrative structures of GM and GAG within mobile domain, highlighting key sugar units with labels: galactofuranose (Gal*f*, red), a-1,2-mannose (Mn^1,2^, pale brown), a-1,6-mannose (Mn^1,6^, dark bronze), galactopyranose (Gal, yellow), galactosamine, and N-acetylgalactosamine (GalNAc, magenta). (**g**) Plots of the relative abundance of all assigned mobile carbohydrates. The relative abundance was determined by integrating the one or two well-resolved peaks corresponding to each monosaccharide, which can be found in **Supplementary table 5**.

Three subtypes of galactofuranose (Gal*f*) residues were identified through characteristic correlation peaks: the first subtype showed C1/C2 signals at 108.9/83.2 ppm and a C4 signal at 83.2 ppm, while two additional subtypes were resolved using similar spectral fingerprints. These Galf residues are known components of galactomannan (GM) sidechains. We further confirmed α-1,2- and α-1,6-linked mannose residues forming the GM backbone (**Fig. 2d**). Distinct fucose signatures were observed in molds, with two variants (F^a^: 62.7/15.4 ppm; F^b^: 60.7/16.1 ppm) identified via C5/C6 correlations, whereas yeast exhibited only one fucose type (58.5/17.8 ppm) (**Fig. 2b**). Notably, all fucose residues displayed downfield C2 shifts (79.7/80.6 ppm in molds; 77.8 ppm in yeast), indicative of 1,2-glycosidic linkages (**Fig. 2c**).

Comparative analysis of 2D spectra between mold (25°C) and yeast (37°C) cell walls of *T. marneffei* revealed the absence of specific galactose subtypes (b, d, e) in yeast form (**Fig. 2d**). Cross-referencing these spectra with published data for *R. delemar* (*33*)and *A. fumigatus*(*42*) enabled preliminary identification of amino sugars in molds, including *N*-acetylgalactosamine (GalNAc; C1: 95.7 ppm, C2: 57.6 ppm) and galactosamine (GalN; C1: 91.6 ppm, C2: 54.7 ppm), which were undetectable in yeast (**Fig. 2e** and **Supplementary figure 5**). In the mobile domain, type-a galactose exhibited a downfield C4 chemical shift (82.1 ppm) with C1 resonating at 101.1 ppm, confirming a 1,4-glycosidic linkage. However, the anomeric conformation (α or β) of galactose within fungal cell walls remains unresolved. While the anomeric carbon of β-monosaccharides typically resonates at higher ^13^C chemical shifts (downfields) than α-monosaccharides due to the interaction of electron charge clouds between C1 and the ring oxygen,(*55*) this distinction could not be conclusively determined here.

The detected amino sugars and the 1,4-linked galactose subtype suggest a likelihood of galactosaminogalactan (GAG) contributing to formation of hyphal cell walls of *T. marneffei* (**Fig. 2f**). GAG, a linear heteropolymer of α-1,4-linked galactose and *N*-acetylgalactosamine, is a well-characterized virulence factor in *A. fumigatus*, where it mediates biofilm formation and immune evasion.(*56*) However, beyond its functional role as validated in *Aspergillus* species,(*42*) the presence of GAG in *T. marneffei* remains poorly documented, and additional research is required to confirm whether GAG is present or absent in this fungal species. Consequently, further investigation is needed to resolve this uncertainty.

Mobile carbohydrate profiles were largely conserved between mold and yeast forms, yet galactosamine, GalNAc, and galactose subtypes b/d/e were disappeared during mold-to-yeast transition. This parallels observations in *A. fumigatus* GAG-deficient mutants, where GalN and GalNAc levels plummet post-mutation.(*42*) Arabinose, a five-membered ring sugar reported in *Aspergillus*, was also detected (C1 106–109 ppm, C2 87–88 ppm).(*49*) Intriguingly, β-1,3-glucans, typically rigid components, appeared in the mobile domain, evidenced by C1 (103–104 ppm), C2 (74–75 ppm), and C3 (84–87 ppm) shifts (**Supplementary table 9**). Unlike *R. delemar*, no mobile chitin or chitosan was observed, underscoring their inherent rigidity.(*33*)

The dual presence of β-1,3-glucan in rigid and mobile domains suggests two structural possibilities: either a mobile subdomain exists within the rigid layer, or β-1,3-glucan occupies the interface between layers. This glucan may act as a molecular bridge, covalently linking rigid chitin to mobile galactomannan (GM) to form a chitin–β-1,3-glucan–GM complex, like other fungal species.(*57*) Such an architecture could balance structural integrity with dynamic adaptability, reflecting fungal strategies to modulate cell wall composition in response to environmental cues.

Following the assignment of mobile carbohydrate peaks, we performed a compositional analysis based on well-resolved spectral signals (**Supplementary table 5**), with results presented in **Fig. 2g**. The analysis quantified key polysaccharides in both hyphal and yeast cell wall forms. b-1,3-glucan represented 5.5% of hyphal cell walls and increased to 9.1% in yeast. Galactofuranose (Gal*f*) constituted 26.1% in hyphae and 28.1% in yeast, while mannose residues (both a-1,2 and a-1,6-mannose) accounted for 16.2% in hyphae and 15.3% in yeast. Arabinose levels were 23.9% in hyphae and 21.0% in yeast. Galactose comprised 24.6% of hyphal cell walls but decreased to 21.8% in yeast. Fucose showed a marked increase from 0.5% in hyphae to 2.7% in yeast. Notably, amino sugars were detected at 0.5% in hyphae but fell below detection limits or became absent in yeast, consistent with their phase-dependent absence observed in prior analyses. These quantitative shifts highlight dynamic reorganization of carbohydrate composition during morphological transition.

### Dynamics and water hydration of wall carbohydrates

The dynamic properties of rigid cell wall polysaccharides, crucial for maintaining cell wall stiffness, were investigated using 1D ^13^C Torchia CP at the nanosecond (ns) timescale (**Fig. 3a** and **Supplementary table 6**).(*58*) Molecules exhibiting fast ^13^C-T_1_ relaxation are highly dynamic, a characteristic likely attributed to rapid local reorientation motions. Likewise, molecules with slow ^13^C-T_1_ relaxation are relatively rigid, contributing to a stiffer structure because of constraints imposed by their local environments. For both forms of *T. marneffei*, the chitin polymers are the slowest measured with average ^13^C-T_1_ 7.95 ± 2.32 s for the hyphal form and average ^13^C-T_1_ 6.97 ± 2.20 s for the yeast form. The slowest dynamics of chitin polymers means that chitin plays a significant role in maintaining the structural rigidities of the stiff domains. The large standard deviation suggests that the chitins in both forms likely exhibit varying degrees of heterogeneity, influenced by their local interactions with neighboring polymers.

**Fig. 3.**
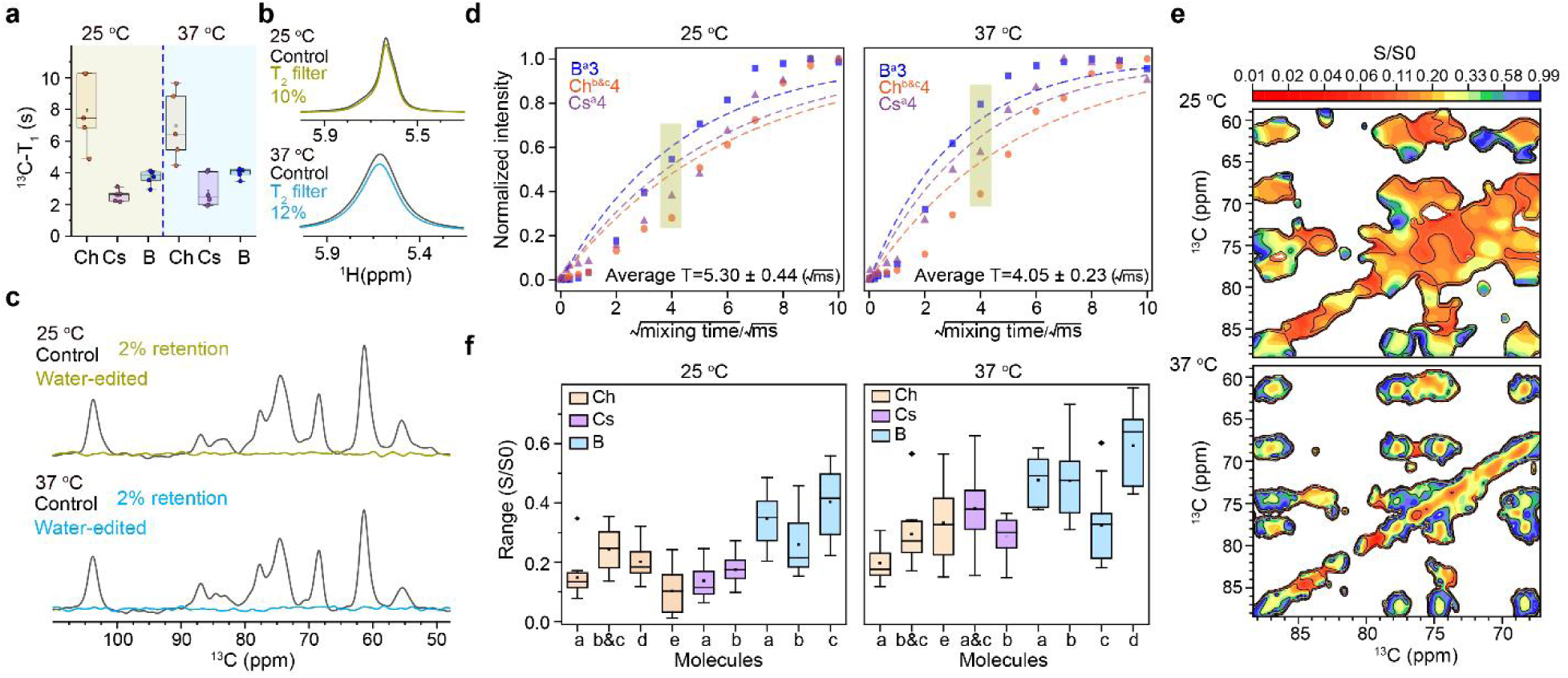
Dynamics and water hydration. (**a**) ^13^C–T_1_ relaxation times of carbohydrates plotted in boxes (color-coded for all three molecules) for both molds and yeast, which represent nanosecond-timescale motions. (**b**) ^1^H-T_2_ filtered (gold yellow-25 °C and blue-37 °C, black-control without ^1^H-T_2_ filter) ^1^H NMR spectra, with 10% (25 °C) or 12% (37 °C) of water signal reduction after using ^1^H T_2_ filter. (**c**) ^1^H-T_2_ filtered (gold yellow-25 °C and blue-37 °C) and control (black) ^13^C spectra are shown for both samples cultured under different temperatures, where the ^13^C signal intensities were reduced to 2% for both samples after ^1^H T_2_ usage. (**d**) Representative ^13^C signal accumulation curves for carbohydrates are shown, with carbon atoms in each compound color-coded for clarity: β-1,3-glucan, chitin, and chitosan are highlighted in light blue, light yellow, and purple, respectively. Among these, β-1,3-glucan exhibited the fastest ^13^C signal saturation, while overall accumulation occurred more rapidly at 37 °C compared to 25 °C. These trends were validated by the fitted rate constant T, derived from the exponential equation y = 1 − A × exp(−T/x), which describes the signal buildup dynamics. (**e**) The hydration profiles of carbohydrates were mapped for the rigid polysaccharides in both the hyphal and yeast forms using 2D ^13^C-^13^C water-edited correlation spectroscopy. (**f**) Box-and-whisker plotting the relative intensities (S/S0) of b-1,3-glucan (blue-25 °C, n=44, blue-37 °C, n=48), chitin (orange-25 °C, n=53, blue-37 °C, n=35), and chitosan (purple-25 °C, n=33, purple −37 °C, n=24) in both hyphal and yeast forms of *T. marneffei*.

Interestingly, both chitosan and β-1,3-glucans demonstrated relatively rapid relaxation dynamics, as indicated by their average ¹³C-T_1_ relaxation times. In the hyphal form of *T. marneffei*, chitosan exhibited a relaxation time of 2.57 ± 0.38 s, slightly shorter than its yeast form value of 2.84 ± 1.02 s. Similarly, β-1,3-glucan showed relaxation times of 3.74 ± 0.45 s in hyphae and 4.01 ± 0.29 s in yeast, with these trends consistent across other fungal species. These results suggest that β-1,3-glucans undergo moderate relaxation, while chitosan displays more dynamic behavior. The enhanced mobility of chitosan could originate from its positioning within outer lamellar regions near rigid chitin layers, like the disordered N-terminal fragments of protein assembly gain structural flexibility through unrestricted isotropic rotational motion.(*59*) An alternative possibility is that chitosan’s shorter polymer chains and weaker topological constraints decrease intermolecular interactions relative to longer, organized assemblies.(*60*) Despite these differences, both hyphal and yeast forms of *T. marneffei* maintained similar dynamic properties, implying that the rigid carbohydrate domains retain a stable local environment during morphological transitions, potentially due to the preservation of individual carbohydrate assemblies.

The water association of fungal cell wall polysaccharides, which arises from the collective perturbation of confined interfacial water molecules surrounding these macromolecules, is closely linked to their biological functions.(*61–63*) Following the application of a ^1^H T_2_ filter (2 × 1 ms) to reduce ^1^H signal intensities to 90% (hyphal form) and 88% (yeast form) (**Fig. 3b**) and lowering the ^13^C signal intensities of polysaccharides to 3%-5% without ^1^H spin diffusion time (**Fig. 3c**), we ran a series of 1D ^13^C spectra using ^1^H-^1^H mixing times spanning 0.1 μs to 100 ms (**Supplementary figure 9**). For both samples, β-1,3-glucan exhibited the fastest signal buildup rate compared to chitin or chitosan, while the overall buildup rates of hyphal carbohydrates were slower than those of yeast polysaccharides. Buildup rate constants, derived by fitting normalized data to the exponential equation y=1-A×exp(-x/T), yielded average values of 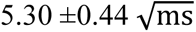 for hyphal carbohydrates and 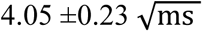 for yeast polysaccharides (**Fig. 3d**) and details can be found in **Supplementary table 7**. Water accessibility for both fungal forms was examined using 2D ^13^C-^13^C correlation water-edited experiments. These experiments utilized a 4 ms ^1^H-^1^H mixing time to facilitate water magnetization transfer to the polysaccharides. The relative intensities (S/S0) of a hydration heatmap were plotted to provide a clear visualization of differences in water associations of major rigid polysaccharides in response to varying cultivation temperatures (**Fig. 3e**). The heatmap results clearly demonstrated that the cell wall polysaccharides in the hyphal form exhibit greater hydrophobicity compared to their counterparts in the yeast form.

Furthermore, we extracted 1D slices for each individual polysaccharide based on specific carbon sites, calculated the relative ratios of S/S0, and presented the statistical results in **Fig. 3f** and were summarized in **Supplementary table 8**. For the hyphal form (25°C), the average S/S0 values were 0.15 ± 0.07 for type-a chitin, 0.24 ± 0.07 for type-b&c chitin, 0.20 ± 0.06 for type-d chitin, and 0.10 ± 0.08 for type-e chitin. In contrast, for the yeast form (37°C), the average S/S0 values were 0.20 ± 0.06 for type-a chitin, 0.29 ± 0.11 for type-b&c chitin, and 0.33 ± 0.13 for type-e chitin. Type-a and type-e chitins in the hyphal cell wall exhibited the highest hydrophobicity, with the lowest S/S0 values. While the hydration level of type-a chitin in the yeast cell wall showed a slight increase, that of type-e chitin increased significantly. For β-1,3-glucans, the most abundant carbohydrates in the hyphal cell wall, the S/S0 ratios were 0.35 ± 0.08 for type-a, 0.26 ± 0.10 for type-b, and 0.40 ± 0.11 for type-c. In the yeast cell wall, the S/S0 ratios for β-1,3-glucans were 0.48 ± 0.07 for type-a, 0.47 ± 0.12 for type-b, 0.32 ± 0.13 for type-c, and 0.59 ± 0.14 for type-d. The variations observed among the different types of chitins and β-1,3-glucans indicate the formation of subdomains with distinct hydration environments, contributing to the complexity of the molecular architecture.

Chitosan, the least abundant rigid carbohydrate in the hyphal cell wall, had average S/S0 values of 0.14 ± 0.06 for type-a and 0.17 ± 0.05 for type-b. In the yeast form, the S/S0 ratios for chitosan were 0.38 ± 0.12 for type-a&c and 0.29 ± 0.07 for type-b. The identified chitosan isoforms displayed similar hydration levels, suggesting they occupy comparable local water environments within the rigid core.

Overall, β-1,3-glucans (average S/S0 of 0.34 ± 0.11 in the hyphal form and 0.47 ± 0.15 in the yeast form) were identified as the most hydrophilic components. In comparison, chitins (average S/S0 of 0.18 ± 0.09 in the hyphal form and 0.27 ± 0.11 in the yeast form) and chitosans (average S/S0 of 0.16 ± 0.06 in the hyphal form and 0.33 ± 0.10 in the yeast form) showed similar hydration patterns. These trends were also consistent across other fungal species. Notably, all rigid components exhibited increased hydrophilicity during the transition from the hyphal to the yeast form.

### Conformational variations of proteins between molds and yeast of *T. marneffei*

Fungal proteins play a critical role in infection and survival, contributing to pathogenicity by participating in immune evasion, host cell adhesion, and intracellular persistence.(*64–66*) However, unlike traditional biochemical methods which often disrupt their structural integrity or interactions, characterizing these proteins in their native state is challenging. In this study, we utilized solid-state NMR spectroscopy to analyze site-specific ¹³C chemical shifts in protein backbone and sidechain carbons, preserving their native conformational states while mapping the functional residues of amino acids.

We first focused on rigid amino acids, using 2D ¹³C-¹³C correlation spectra with a 53-ms CORD mixing period to identify intramolecular correlation peaks. These results are shown in **Fig. 4a** and **Fig. 4b**, and comprehensive chemical shift assignments and spectral interpretations were provided in **Supplementary figures 10a and 10b** and **Supplementary table 11**. The analysis began by correlating ¹³C chemical shifts of backbone carbons (Cα and CO) with side-chain carbons (Cβ, Cγ). For example, phenylalanine residues were identified by tracking Cα and Cβ shifts alongside downfield-shifted aromatic Cγ correlations. Alanine residues were distinguished by their characteristic Cα (<55 ppm) and Cβ (<25 ppm) shifts, contrasting with valine residues, which exhibited a downfield Cα (>55 ppm) and distinct Cβ (∼30 ppm) and Cγ (∼18 ppm) shifts.(*67*) This method allowed us to resolve 36 rigid amino acid residues in molds and 34 in yeast. We performed secondary structure analysis by comparing the observed Cα chemical shifts to random-coil values, as protein secondary structure is highly associated with φ and ψ torsion angles.(*68, 69*) Preliminary data suggested that rigid proteins in both molds and yeast predominantly adopts α-helical conformations, with structural variations emerging in response to temperatures.

**Fig. 4.**
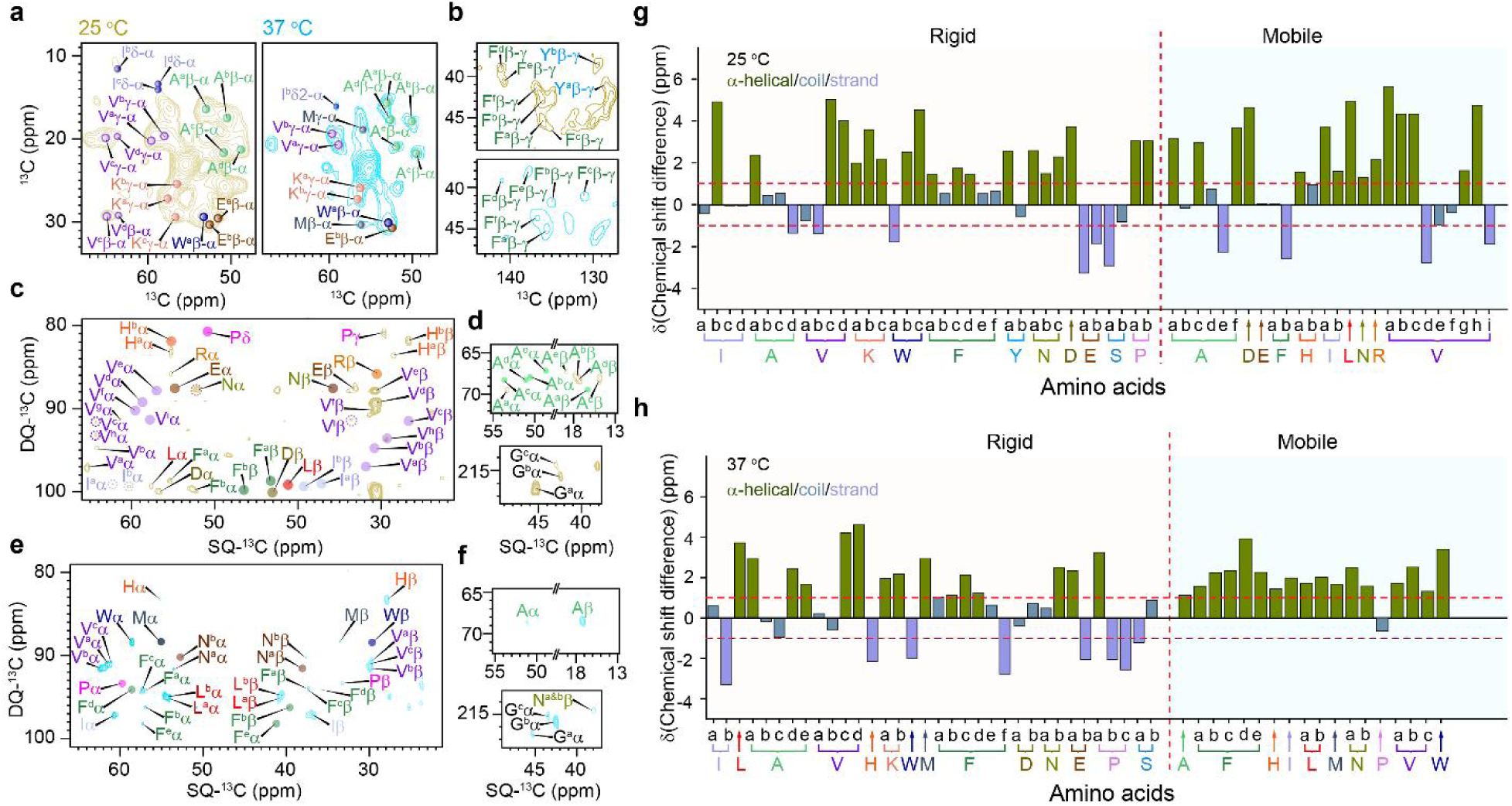
Proteins identified within *T. marneffei*. (**a**) and (**b**) Rigid proteins were identified using 2D ^13^C-^13^C correlation spectra with 53-ms CORD mixing, where amino acids identified for the mold (25 °C) and yeast (37 °C) samples were plotted in gold yellow and blue, respectively. (**c**) and (**d**) Mobile proteins of the molds (25 °C) were assigned based on their fingerprint Ca and Cb as well as their sidechain ^13^C chemical shifts. (**e**) and (**f**) Mobile proteins of the yeast (37 °C) were assigned based on their fingerprint Ca and Cb as well as their sidechain ^13^C chemical shifts. (**g**) and (**h**) Secondary structure of proteins determined by ^13^C chemical shifts of Ca, a-helical, coil, and b-strand conformations are in olive green, blue gray and periwinkle, where the horizontal red dash lines are error bars and the vertical red dash lines are for the separation of rigid and mobile AAs. All the identified amino acids were presented as one-letter codes, for example, alanine is abbreviated as A.

Further analysis revealed conformational differences between molds and yeast for specific amino acid subtypes (**Fig. 4g** and **4h**). For isoleucine, molds exhibited four subtypes, two of which were random-coiled, while yeast displayed two subtypes, one of which was random-coiled. Type-b isoleucine adopted α-helical conformations in molds but β-strand in yeast. Alanine subtypes also showed divergence: molds displayed four subtypes (type-a α-helical, type-b/c random-coiled, type-d β-strand), whereas yeast showed five subtypes (type-a α-helical, type-b/c random-coiled, type-d/e α-helical). Valine conformations remained largely consistent, except for type-b, which shifted from β-strand in molds to coiled in yeast. Phenylalanine subtypes aligned structurally, except for type-f, which transitioned from coiled in molds to β-strand in yeast.

Asparagine/aspartic acid in molds were mostly α-helical, while their yeast counterparts were predominantly random-coiled. Glutamic acid adopted β-strand conformations in molds, with one yeast subtype following suit. Proline in molds favored β-strand conformations, while yeast proline included both β-strand and α-helical types. Serine subtypes showed minimal variation, although yeast type-a exhibited smaller Cα deviations from random-coil values. Lysine residues in both phases adopted α-helical conformations, with mold type-b displaying enhanced structural order. Tryptophan residues in molds were observed exclusively in α-helical conformations, whereas yeast exhibited an additional β-strand subtype. Tyrosine residues were uniquely detected in molds, while histidine, leucine, and methionine appeared solely in yeast. The absence of specific amino acids in either phase does not confirm their nonexistence but likely reflects limitations in detection sensitivity. This may arise from their low abundance, inherent structural flexibility, or a combination of both factors, which can attenuate spectroscopic signals and hinder observation. Overall, proteins in both phases primarily retained α-helical secondary structures, despite phase-specific conformational variations.

We also characterized the mobile amino acids in both mold and yeast forms using 2D ¹³C DP J-refocused INADEQUATE spectra (**Fig. 4d, 4e** and **4f**), with detailed peak assignments provided in supplementary information (**Supplementary figures 10c**, **10d**, **10e** and **10f**) and corresponding ¹³C chemical shifts listed in **Supplementary table 12**. This analysis identified 28 mobile amino acid residues in molds and 20 in yeast, with 11 common residues shared between both phases, including alanine, phenylalanine, and glycine, while each phase exhibited unique residues, such as methionine and tryptophan in yeast.

To further investigate secondary structure differences, we analyzed Cα chemical shift deviations from random-coil reference values and found distinct conformational preferences between the two phases (**Fig. 4g** and **4h**). In molds, seven residues adopted random-coil conformations, and four exhibited β-strand characteristics, whereas yeast showed only one random-coil residue and no β-strand formation. Both phases displayed a strong tendency toward α-helical structures, with 17 residues in molds and 19 in yeast adopting this conformation. Similar to rigid proteins, mobile proteins in both molds and yeast predominantly adopted α-helical secondary structures.

These observed variations align with gene expression patterns during the dimorphic transition, where rapid upregulation of stress-responsive genes (pepA, ftrA, sidA, hmgR) facilitates early adaptation during the mold-to-yeast shift, while delayed-response genes such as hgrA, which governs hyphal cell wall integrity, activate later in the yeast-to-mold transition.(*52*) Such differential gene regulation likely influences protein synthesis and structural organization. While this approach offers residue-level structural insights without disrupting native protein states, a remaining challenge is the precise assignment of these spectral signatures to specific proteins within the complex cellular environment.

### Temperature-induced modifications in the molecular packing of the cell walls

The morphological transition from hyphal to yeast has been reported to signify a shift from the saprophytic phase to the parasitic phase, enabling the fungus to act as a causative agent in immunocompromised individuals.(*70, 71*) Following the detailed analyses of relative compositions and the identification of characteristic polysaccharides, it is essential to further investigate the supramolecular assembly of the fungal cell walls in both forms of *T. marneffei*. Molecular interactions play a vital role in maintaining the mechanical properties of the cell wall and providing a structural basis for the development of potential antifungal therapies.(*26, 72, 73*) To explore these interactions, we employed 2D ^13^C-^13^C PDSD (Proton-Driven Spin Diffusion) with a long mixing time of 1 s, compared with 53-ms 2D ^13^C-^13^C CORD spectra, allowing us to probe intermolecular interactions among polymers at a sub-nanometer scale (**Fig. 5**).(*74*) The observed intermolecular interactions arise from multiple sources and can be classified into three main categories (**Supplementary table 13**): 1) long-range interactions between subtypes of carbohydrate polymers, such as chitin-chitin, chitosan-chitosan, β-1,3-glucan-β-1,3-glucan, and others; 2) long-range interactions between different types of polysaccharides, including chitin-chitosan, chitosan-β-1,3-glucan, and chitin-β-1,3-glucan; and 3) intermolecular interactions between carbohydrates and proteins. Due to the overlap and inherent uncertainties (∼0.5 ppm of FWHM) in the chemical shifts of certain polysaccharides, intermolecular interactions are accounted for across all types. For example, the interaction labeled as Ch^a/b/c^/B^a^5-Cs^a^3 is interpreted as representing two distinct interactions: one between chitin and chitosan, and another between β-1,3-glucan and chitosan, when analyzing specific interactions between different molecular types. First, we identified 30 intermolecular interactions between chitin-chitin in the hyphal form and 52 such interactions in the yeast form. Additionally, we observed 8 intermolecular interactions between chitosan-chitosan in the hyphal form and 13 in the yeast form. Furthermore, 26 intermolecular interactions between β-1,3-glucan-β-1,3-glucan were detected in the hyphal form, compared to 23 in the yeast form. The intermolecular interactions between subtypes of the same isoforms of polysaccharides indicate a highly polymorphic supramolecular architecture. This indicates that polysaccharides of the same type, yet with different chemical environments, play a role in creating distinct subdomains within the rigid core. This phenomenon was also reported for other fungal cell walls.(*34*) The yeast form exhibits a greater number of intermolecular interactions between the same type of polysaccharides with different chemical environments compared to the hyphal form, suggesting a more tightly packed spatial arrangement in the yeast form.

**Fig. 5.**
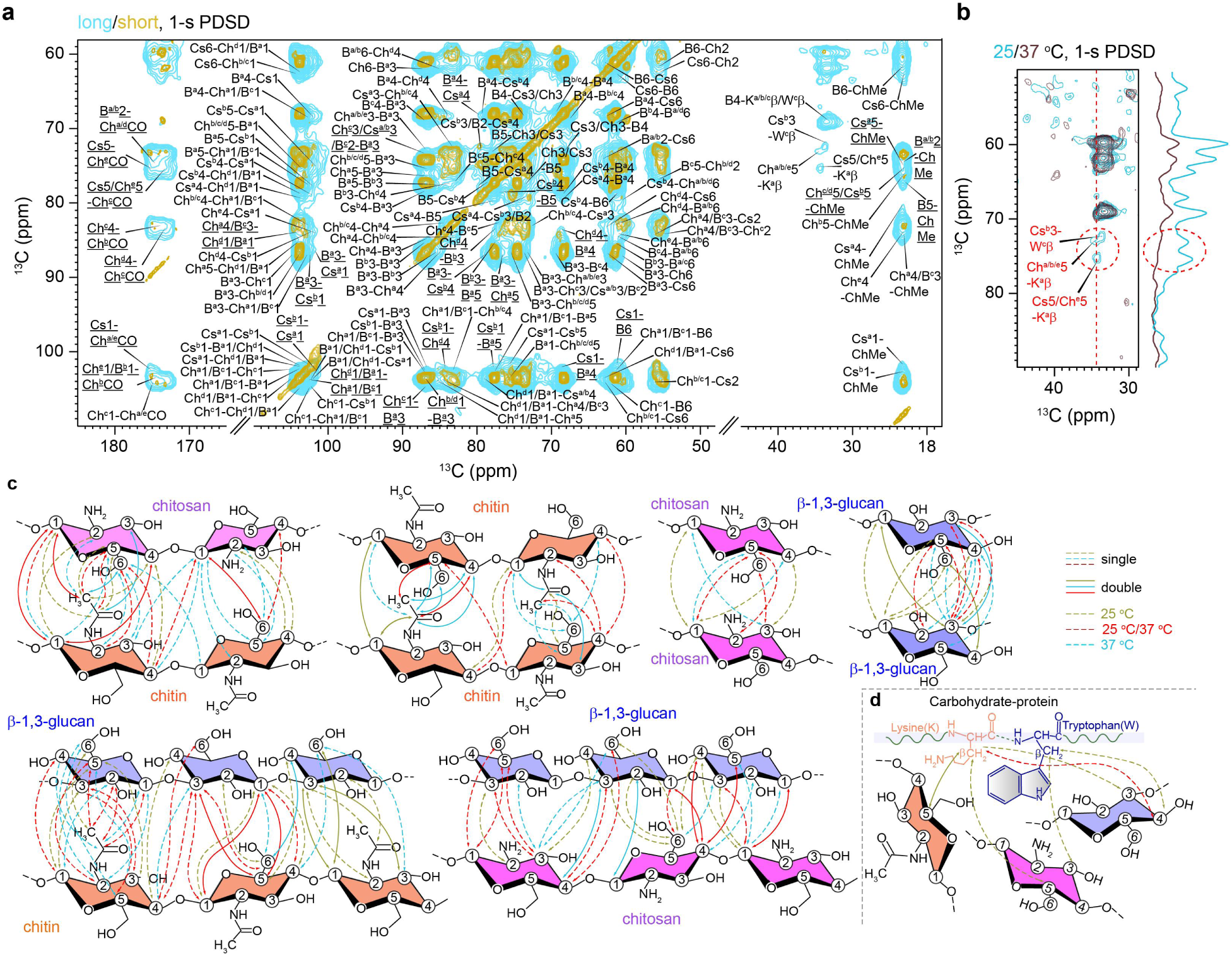
Intermolecular interaction identified in both forms of *T. marneffei* (mold, 25 °C and yeast, 37 °C). (**a**) Overlay of 2D ^13^C-^13^C correlation spectra with short 53-ms CORD mixing (gold yellow) and 2D ^13^C-^13^C PDSD using an extended mixing of 1 s (Cyan blue). (**b**) Long-range contacts between carbohydrates and proteins, where 1D ^13^C slices of F_1_ dimension from 2D ^13^C-^13^C PDSD with 1 s mixing were extracted for clarity (cyan blue-25 °C, dark purple-37 °C). (**c**) Schematic illustration of the intermolecular interactions inferred from the NMR spectra, with arrowheads indicating the directionality of polarization transfer. Olive lines represent interactions specific to the hyphal form at 25°C, red lines indicate interactions common to both yeast and hyphal forms, and cyan blue lines correspond to those in the yeast form at 37°C. Dashed lines, following the arrowheads, signify single interactions of a given type, whereas solid lines indicate double interactions of the same type. (**d**) Illustrative interactions between carbohydrates and proteins, highlighting specific amino acid residues such as lysine (K) and tryptophan (W).

Secondly, intermolecular interactions between different types of polysaccharides were also observed. Specifically, 45 interactions between chitin and chitosan were identified in the hyphal form, compared to 59 in the yeast form. Additionally, 70 interactions between chitin and β-1,3-glucan were identified in the hyphal form, compared to 61 in the yeast form. Similarly, 48 interactions between chitosan and β-1,3-glucan were observed in the hyphal form, while 51 were detected in the yeast form. In contrast to the intermolecular interactions between different isoforms of the same type of polysaccharides, the interactions among different types of rigid polysaccharides further reinforce the supramolecular assembly of the fungal cell wall architecture. In other words, these polysaccharides not only interact within their own groups but also exhibit spatial proximity with neighboring polysaccharides of different types, resulting in a more tightly packed cell wall. From the perspective of polysaccharide interactions within the cell wall, the yeast form of *T. marneffei* demonstrates more extensive polysaccharide interactions than the mold form. This could explain the morphological changes observed during the phase transition, including the significant thickening of the cell wall. Such changes likely require enhanced interactions between structural molecules to maintain cell wall integrity and protect against potential immune system recognition and attack.

After gaining insights into the intermolecular interactions among carbohydrates, we extended our analysis to investigate the interactions between carbohydrates and proteins. While most proteins, along with certain mobile polysaccharides, are typically highly mobile in fungal cell walls, making it challenging to directly measure their interactions with carbohydrates in their native state. Following this, we performed a comparative analysis of our 2D ^13^C-^13^C PDSD with a 1-s mixing time and the 2D ^13^C-^13^C correlation spectra. This analysis uncovered cross peaks between carbohydrates and proteins in the hyphal form of *T. marneffei*, with these interactions linked to specific amino acid residues, i.e., lysine and tryptophan (**Fig. 5b**). Specifically, in the hyphal form, we identified three long-range interactions between type-a β-1,3-glucan and the Cβ of type-a lysine, two interactions of type-a lysine with either type-a/b/e chitin or chitosan, and two interactions between type-c tryptophan and type-b chitosan, as drawn in **Fig. 5d**. In contrast, no protein interactions with other types of chitin or chitosan were observed in the yeast form, and only two interactions involving lysine or tryptophan with β-1,3-glucan were retained. The cell wall proteins are either covalently linked to wall polysaccharides or interact with other wall proteins through van der Waals forces, hydrogen bonding, or ionic interactions. These interactions have been observed in live cells of *R. delemar* as well as in chemically extracted cell walls of *A. fumigatus*.(*33, 42*) Variations in cultivation temperatures not only influence the morphological characteristics of the fungus but also impact the activity of molecular proteins. Moreover, these temperature differences induce a unique spatial arrangement of proteins and polysaccharides, likely facilitating the fungal adaptation of macrophage killing or attacking the host cells.

## DISCUSSION

The cell wall of *T. marneffei*, a thermally dimorphic fungus, is a key biological feature that facilitates its transition between saprophytic mold and pathogenic yeast forms, while also serving as a crucial surface structure for mediating interactions with the host immune system.(*28, 75–77*) Fungal cell wall polysaccharides, which are largely absent in humans, offer promising targets for the development of novel antifungal therapies.(*78*) However, molecular-level research on the cell wall structures of *T. marneffei*’s dimorphic forms is still limited. This study aims to fill this gap by presenting a high-resolution solid-state NMR analysis of *T. marneffei*’s cell walls, revealing the molecular architectures of both the hyphal and yeast forms. First, β-1,3-glucans, chitins, and chitosans are highly rigid and polymorphic, predominantly enriched in the inner layers of the cell wall. These components extensively interact with each other to form the structural core of the inner cell wall. Protein segments are also present in the inner layer, as evidenced by 2D ^13^C-^13^C correlation spectra with 53-ms CORD mixing (**Fig.4a** and **4b**), where certain types of proteins interact with rigid carbohydrates at the sub-nanometer scale. Second, galactomannan (GM), arabinose, mannoproteins, and β-1,3-glucans are cross-linked to form a softer matrix outside the inner layer. Previous chemical analyses of *Aspergillus* species have shown that β-1,3-glucans are covalently linked to chitin and GM, forming a carbohydrate core within their cell walls.(*79*) Given the phylogenetic similarities between the hyphal form of *Talaromyces* and *Aspergillus* species, particularly within the *Eurotiales* order, it is likely that *T. marneffei* shares common features with *Aspergillus*.(*80*) The NMR-detected GM signals (**Fig. 2a**) and the bimodal distribution of β-1,3-glucans across inner and outer layers support the hypothesis that *T. marneffei* may share a carbohydrate core structure containing chitin, β-1,3-glucan, and GM (**Fig. 6b** and **6c**). However, the existence of this proposed complex necessitates additional experimental validation.

**Fig. 6.**
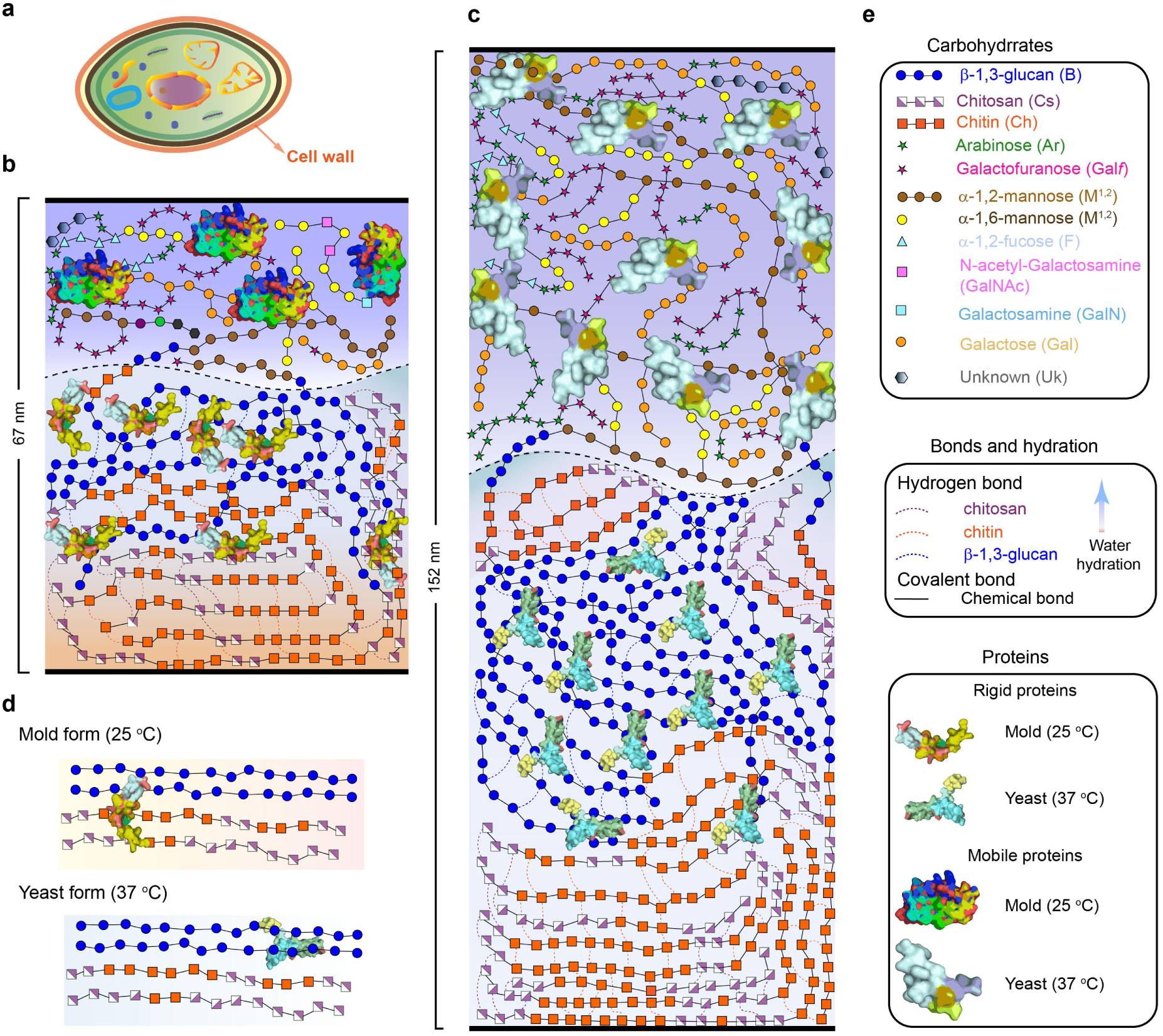
Illustration of the molecular organization in the cell walls of *T. marneffei*. (**a**) Fungal cell where cell wall is highlighted with an orange arrow. (**b-c**) Cell wall of mold (b) and (**c**) yeast. The cell wall undergoes significant thickening, increasing from 67 nm in mycelia to 152 nm in yeast. Within the rigid domain, chitin microfibrils and chitosan, the most hydrophobic components, are interconnected, with chitosan demonstrating the partial deacetylation of chitin fibrils. In contrast, b-1,3-glucan, which is relatively hydrophilic and dynamic, is positioned closer to the outer layer. Some b −1,3-glucan molecules are directly bonded to chitin, reflecting their comparable hydration levels, or are linked to chitin and GM (glycosylphosphatidylinositol-modified proteins) to form a complex chitin-b-1,3-glucan-GM structure. Mobile polysaccharides and proteins are situated at the top layer of the cell wall. (**d**) β-1,3-glucan-tryptophan interactions occur in both forms, whereas chitin/chitosan interactions with lysine-containing proteins are specific to the hyphal cell wall and absent/negligible in yeast. (**e**) Symbols for carbohydrates in **Fig. 6b, 6c**, and **6d** follow the notation defined by Glycano (http://glycano.cs.uct.ac.za). Color-coded dashed lines indicate hydrogen bonds between carbohydrates, while black solid lines denote covalent bonds. Structural illustration for rigid and mobile proteins were generated by randomly assembling identified amino acid residues into sequences; their structures were subsequently predicted using AlphaFold (https://alphafoldserver.com).

The polymorphic nature of chitin and its derivative chitosan is not surprising, as seven chitin synthase genes have been analyzed to modulate the dimorphic fungus. As reported, *CpCHS2*, *CpCHS3*, and *CpCHS6* are expressed during the saprobic phase, *CpCHS1* and *CpCHS4* are associated with the pathogenic phase, while *CpCHS5* and *CpCHS7* are active in both phases.(*81, 82*) The differential expression of these chitin synthase genes likely results in variations in chitin synthase (CHS) activity between the hyphal and yeast forms of *T. marneffei*. This may explain the absence of type-d chitin in yeast cell wall, potentially due to temperature-induced variations in the expression of chitin synthase-associated genes. Surprisingly, a specific subtype of chitosan (type-c) was uniquely detected in the yeast form of *T. marneffei*, whereas type-e chitin is found within the cell walls of mycelia but is conspicuously absent in those of the yeast. On the one hand, the transformation of chitin into chitosan is facilitated by chitin deacetylases (CDAs), enzymes that are ubiquitous across numerous fungal species.(*83, 84*) Temperature-driven morphological changes in fungi may trigger the activation or modify the expression levels of enzymes involved in polysaccharide biosynthesis, thereby modulating the production of associated cell wall components. These modifications highlight the intrinsic biological capacity of fungi to dynamically restructure their cell walls, enabling adaptation to variable environmental conditions. Similar phenomena have been documented in *Rhizopus* species, where environmental stressors resulted in the loss of a particular chitosan variant—a potential adaptive strategy to enhance survival under adverse conditions.(*33*) During the transition from mold to yeast, the relative amounts of both chitin and chitosan decline. This pattern is consistent with observations in the dimorphic fungus *Benjaminiella poitrasii*, where chitin levels are higher in the hyphal stage than in the yeast form. This could be linked to the regulatory role of CHS in the yeast-mold transition.(*85*) The degree of chitin deacetylation remains largely consistent across both fungal forms, highlighting the stability and robustness of specific chitin deacetylases (CDAs) under temperature fluctuations. This contrasts with *in vitro* studies, where CDA activity is influenced by variables such as temperature, pH, and salt concentrations. For example, the CDA from *Pestalotiopsis* sp. exhibits optimal deacetylation at pH 8.0 and 55 °C *in vitro*.(*86*) In contrast, *in vivo* CDA functionality is more complex. Anchored CDAs, tethered to the plasma membrane via glycosylphosphatidylinositol (GPI) anchors, act on nascent chitin polymers extruded by chitin synthases (CHSs) in chitosan-containing cell walls. Non-anchored CDAs, meanwhile, may target chitin oligomers released during host-pathogen interactions, such as by host chitinases.(*87, 88*) These intricate in vivo mechanisms, where CDAs operate in spatially constrained, dynamic environments rather than freely in solution, explain the observed discrepancies between in vivo and in vitro deacetylation behaviors.

*T. marneffei* exhibits β-1,3-glucan behavior similar to other fungal species, where β-1,3-glucans exhibit a bimodal distribution across both rigid and mobile domains.(*33, 48*) This observation aligns with prior studies on *Aspergillus* species, in which β-1,3-glucans are covalently bound to rigid chitin and mobile galactomannan (GM), forming a chitin-β-1,3-glucan-GM complex that serves as a structural core.(*48*) This complex likely transfers the high mobility of GM to the associated β-1,3-glucan segments (**Fig. 6b** and **6c**). Additionally, β-1,3-glucans, the most abundant polysaccharides, are highly polymorphic in the cell walls of both hyphal and yeast forms of *T. marneffei*. Earlier research on fungal cell wall b-1,3-glucan synthases (GS) has shown that fungi possess a diverse array of endo- and exo-β-1,3-glucanases, transcripted by multiple genes, which work synergistically with transglycosidases to regulate β-1,3-glucan synthesis during various stages of fungal growth.(*89–91*) The diversity of these GSs is closely linked to the biosynthesis of β-1,3-glucans and provides a plausible explanation for the polymorphic nature of β-1,3-glucans in the cell walls of *T. marneffei*, reflecting common characteristics shared among fungi.(*92, 93*) Type-d β-1,3-glucan is exclusively present in the yeast form of *T. marneffei* and absent in the mold form. β-1,3-glucan serves as a structural component in a fibrous network and acts as a key pathogen-associated molecular pattern (PAMP) recognized by host immune cells, particularly through receptors such as Dectin-1. The observed differences in β-1,3-glucans between the hyphal and yeast forms are likely associated with immune evasion strategies.(*94, 95*) Quantitative analyses of β-1,3-glucan content in dimorphic fungi such as *Blastomyces* and *Paracoccidioides* reveal elevated levels in yeast-phase cell walls compared to hyphal-phase cell walls, aligning with our observations.(*96, 97*) These variations may enable *T. marneffei* to avoid immune detection and enhance its persistence within the host during the morphological transition.

Chitin together with chitosan exhibit the highest hydrophobicity under both cultivation temperatures, while the hydration behavior of β-1,3-glucan is more complex (**Fig.6b** and **6c**). Chitin, a widely conserved structural polymer in the fungal cell wall, acts as a framework for the assembly of other wall components, playing a critical role in maintaining cell wall organization and integrity.(*84*) Both isoforms of chitosan (type-a and type-b) in hyphal cell walls exhibit a similar hydration level to type-A chitin but lower than other types of chitins detected in the hyphal cell wall. This suggests that chitosan may form a hydrophobic microdomain within the rigid core, a phenomenon also observed in molds of other fungal species.(*33, 48*) However, at physiological temperatures, the rigid chitosan undergoes remodeling, adapting to form semi-dynamic regions within the rigid core. These regions potentially act as a bridge and buffering zone among polysaccharides within the rigid core. This characteristic of chitosan results in a relatively higher hydration level compared to chitin, with hydration levels comparable to type-b β-1,3-glucan. This echoes with findings by Kollar et al., who identified a complex formed by chitin and β-1,3-glucan linked via a β-1,4 linkage in yeast cell wall of *S. cerevisiae*.(*98*) Recent NMR studies on fungal cell walls indicate that β-1,3-glucan polymers surround the inner layer, which consists of chitin, chitosan, and other polysaccharides within the rigid core, and play a role in regulating water accessibility.(*33, 48*) This aligns with our findings, which show that β-1,3-glucan exhibits higher levels of hydration (**Fig. 3e**). Interestingly, the hydration levels showed a clear increase in response to increased temperature, with the yeast form displaying greater hydrophilicity (**Fig. 3e** and **3f**). This change likely represents an adaptation to the host environment, where higher hydration could aid in immune evasion and stress resistance.

GM has been widely utilized as a screening and adjunct diagnostic standard for detecting *T. marneffei* infections.(*99*) Our NMR data identified a significant presence of its building units, including Gal*f*, α-1,2-mannose, and α-1,6-mannose (**Fig. 2a** and **2d**). Given the widespread distribution of β-1,3-glucan in both rigid and mobile domains, it is plausible that some rigid chitin and partially mobile β-1,3-glucan are covalently linked with GM to form the GM-β-1,3-glucan-chitin complex. This hypothesis is strongly supported by earlier chemical analyses and ssNMR studies of *Aspergillus* species, where mobile GM, bimodal β-1,3-glucan, and chitin are linearly covalently bonded to form the GM-β-1,3-glucan-chitin complex.(*79*) In contrast to the low abundance of Gal*f* in *Aspergillus* species, Gal*f* constitutes nearly 30% of the entire mobile domain in *T. marneffei*, while α-1,2-mannose contributes less than half of that amount. If the GM backbone is constructed by mannose and Gal*f* and α-1,2-mannose are present in a 1:1 molar ratio, this suggests that not all mobile Gal*f* participates in GM formation, and a portion may be cross-linked with other components to fulfill structural roles.(*100*) Alternatively, if the molar ratio of Gal*f* to α-1,2-mannose within GM is not 1:1, the Gal*f* units could chemically link to the α-1,2-mannose backbone at various sites. Therefore, further investigation is necessary to clarify the unexpectedly high content of Gal*f* and its functional implications. An intriguing finding is the potential formation of GAG in the mobile domain of the hyphal cell wall, as indicated by the presence of its building units (GalN, GalNAc, and Gal) (**Fig. 2d**), while it is absent or undetected in the yeast cell wall. Although GAG is common in most fungal species and is typically a structural component of *Aspergillus* fungi, further experimental evidence is needed to confirm whether these units (GalN, GalNAc, and Gal are covalently linked to form GAG within the hyphal cell walls.(*101*) The presence of GalN and GalNAc in the hyphal (**Fig. 6b**) but not in the yeast form may represent a survival strategy of *T. marneffei* during interactions with host cells(**Fig. 6c**). Specifically, at early-stage infections of *T. marneffei*, the cationic GalN (GalNH_3_^+^) plays a crucial role in enhancing *T. marneffei*’s adhesion to anionic surfaces, such as human cells, while also promoting hyphal cohesion. This enables the fungal colony to endure and thrive under unfavorable environmental conditions.(*48, 102*)

The interactions between proteins and polysaccharides display phase-specific behavior (**Fig. 6b** and **6c**). Notably, lysine-containing proteins interact with chitin and chitosan in the hyphal form but are absent or undetectable in the yeast form (**Fig. 6d**). In contrast, the interactions between β-1,3-glucan and tryptophan-containing proteins are strong and consistently observed in both mold and yeast forms, regardless of cultivation temperature. The origin of lysine and tryptophan residues in these interactions remains unclear. Lysine, a positively charged amino acid, is crucial for protein structure and function. In the hyphal cell wall, lysine residues may either form covalent cross-links with other cell wall components, enhancing rigidity and resilience, or facilitate binding to negatively charged host molecules, like the mobile GalN.(*102*) However, our observations primarily focus on the inner layer, where chitin and chitosan, the most hydrophobic components, are surrounded by rigid β-1,3-glucans. Given that chitosan can also carry a positive charge and chitin is neutral under physiological conditions, spatial proximity between lysine and chitosan seems unlikely due to charge repulsion. This suggests that lysine may act as a covalent linker, with its side chain neutralizing its charge to allow proximity to rigid chitin and chitosan.

The source of lysine residues is uncertain, as the cell wall contains numerous proteins, like glycosylphosphatidylinositol (GPI)-anchored enzymes, heat-shock proteins and mannoproteins.(*82, 84*) Mp1p, a lysine-rich antigenic mannoprotein in *T. marneffei*, is widely found in yeast, mold, and conidia and is vital for modulating the host immune response during infection.(*103*) It is plausible that these interactions originate from Mp1p, which is located near the inner layer of the hyphal cell wall. In yeast form, however, Mp1p may be predominantly expressed in the mobile domain, potentially functioning as a fatty acid transporter from the host to the pathogen.(*104*) This may explain why the interactions between chitin/chitosan and lysine are absent or undetectable in the yeast form. The weakening or disruption of spatial interactions between lysine-containing proteins and rigid chitin/chitosan may be counterbalanced by the observed increase in long-range interactions among polysaccharides in the yeast cell wall. Additional studies are necessary to validate these speculations.

Another notable interaction involves β-1,3-glucan/chitosan and tryptophan, which remains stable across varying growth temperatures (**Fig. 6d**). This contrasts with the chitin/chitosan-lysine interaction, which is absent or negligible in hyphal cell walls. Given β-1,3-glucan’s dual role as a structural scaffold and hydrating agent within the inner layer of chitin and chitosan, the observed tryptophan residues may localize near this rigid core. However, the chitosan-tryptophan interaction remains enigmatic, as chitosan’s hydrophobicity and propensity to cross-link with chitin polymers complicate such associations. Notably, echinocandins antifungals targeting β-1,3-glucan synthesis, are ineffective against the yeast form of *T. marneffei*, implying a unique molecular architecture in this pathogen.(*105*) The persistence of β-1,3-glucan/chitosan-tryptophan interactions in both forms may reflect this distinct organization. Similar to lysine, the origin of the tryptophan residues is unresolved, yet their conserved presence highlights a potential therapeutic target for dual-phase antifungal agents against *T. marneffei*.

## CONCLUSION

In conclusion, this study provides a detailed molecular insight into the cell wall architecture of *T. marneffei*, focusing on its dimorphic transition between saprophytic and pathogenic forms. By utilizing high-resolution solid-state NMR, we identified distinct structural features exist in the hyphal and yeast forms, highlighting the critical roles of polysaccharides such as chitin, chitosan, β-1,3-glucan, and galactomannan in cell wall integrity and host interaction. The presence of various chitin synthases and their differential expression across the two phases supports the hypothesis that these structural components are dynamically regulated in response to environmental cues, like temperature shifts. Furthermore, phase-specific interactions between proteins and polysaccharides offer insights into the adaptive strategies that enable *T. marneffei* to evade immune detection during infection. These findings underscore the complexity of the fungal cell wall and its potential as a target for novel antifungal therapies, especially considering the unique molecular architecture of yeast form in *T. marneffei*. Further investigations are needed to validate the precise mechanisms underlying these molecular interactions and their implications for antifungal drug development.

## ACKNOWLEDGEMENT

The project was financially support by Guangxi Science and Technology Base and Talent Project (Grant No. 2022AC19006), Shenzhen Medical Research Fund (Grant No. A2403023), Science and Technology Major Project of Guangxi (Grant No. Guike AA24206048, Guike AA24206050) and National Natural Science Foundation of China (NSFC), (Grant No. 32560050, No. 32470021, and No. 32200024).

