## Supplemental File for "Distinct cell wall molecular architecture of dimorphic *Talaromyces marneffei* cells revealed by solid-state NMR spectroscopy"

### Table of contents

Supplementary figure 1. Flowchart of fungal cell cultivation and tests. 4

Supplementary figure 2. Images of *T. marneffei* cells. 5

Supplementary figure 3. 1D $^{13}$C NMR spectra and estimation of the relative polysaccharides to the protein/lipids 6

Supplementary figure 4. 1D $^{13}$C spectra of polysaccharides. 7

Supplementary figure 5. 2D DP J-based $^{13}$C INADEQUATE spectra for mobile molecules of *T. marneffei*. 8

Supplementary figure 6. Identification of GalN and GalNAc carbohydrates within mobile domain. 9

Supplementary figure 7. Rigid polysaccharides in the *T. mareffei*. 10

Supplementary figure 8. Intermolecular interactions observed within the yeasts 11

Supplementary figure 9. Water-edited experiments setup 12

Supplementary figure 10. Amino acids identified within *T. marneffei* 13

Supplementary table 1. The cell-wall thickness of both molds and yeasts. 14

Supplementary table 2. Solid-state NMR experimental parameters for fungal cell wall characterization. 15

Supplementary table 3. Estimation of molar composition of cellular components within *T. marneffei*. 16

Supplementary table 4. Molar composition of rigid polysaccharides in cell walls. 17

Supplementary table 5. Estimation of relative abundance of mobile cell wall polysaccharides. 18

Supplementary table 6. Relaxation dynamics of rigid polysaccharides by 1D $^{13}$C Torchia CP spectra. 19

Supplementary table 7. Water-hydration dynamics of polysaccharides. 20

Supplementary table 8. Site-specific water-hydration analysis of *T. marneffei*. 21

Supplementary table 9. chemical shifts of rigid polysaccharides in *T. marneffei* cell walls. 23

Supplementary table 10. chemical shifts of mobile polysaccharides in *T. marneffei* cell walls. 24

Supplementary table 11. $^{13}C$ Chemical shifts of identified rigid amino acids in *T. Marneffei* cells. 25

Supplementary table 12. $^{13}C$ Chemical shifts of identified mobile amino acids in *T. Marneffei* cells 27

Supplementary table 13. Intermolecular cross peaks in *T. marnefei.* 28

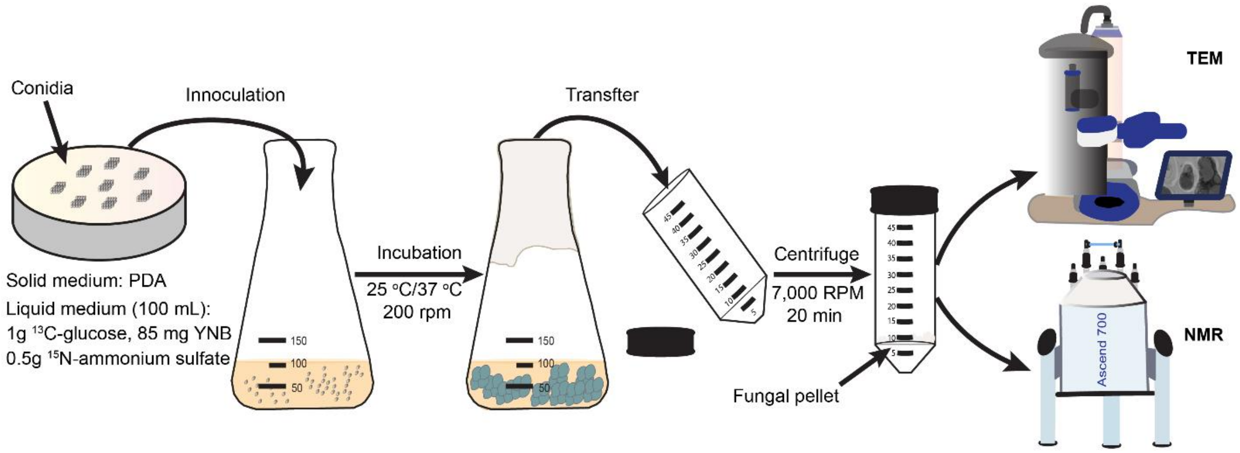

**Supplementary figure 1. Flowchart of fungal cell cultivation and tests.** The experimental workflow for fungal cultivation and sample preparation begins with the inoculation of fungal conidia, pre-cultured on YPD solid medium, into a 250-mL Erlenmeyer flask containing 100 mL of sterile growth medium ($^{13}$C-glucose, $^{15}$N-ammonium sulfate, and Yeast Nitrogen Base (YNB). The flask is then incubated in an orbital shaker at 180 rpm for 14 days at 25°C or 37°C. Following incubation, the culture is transferred to 50-mL centrifuge tubes and pelleted via centrifugation at 10,000 RPM for 30 minutes at 4°C. The resulting pellets are washed three times with phosphate-buffered saline (PBS) to eliminate residual $^{13}$C/$^{15}$N-labeled water-soluble metabolites or media components. Finally, the washed pellets are stored at 4°C until further analysis by transmission electron microscopy (TEM) or solid-state nuclear magnetic resonance (ssNMR).

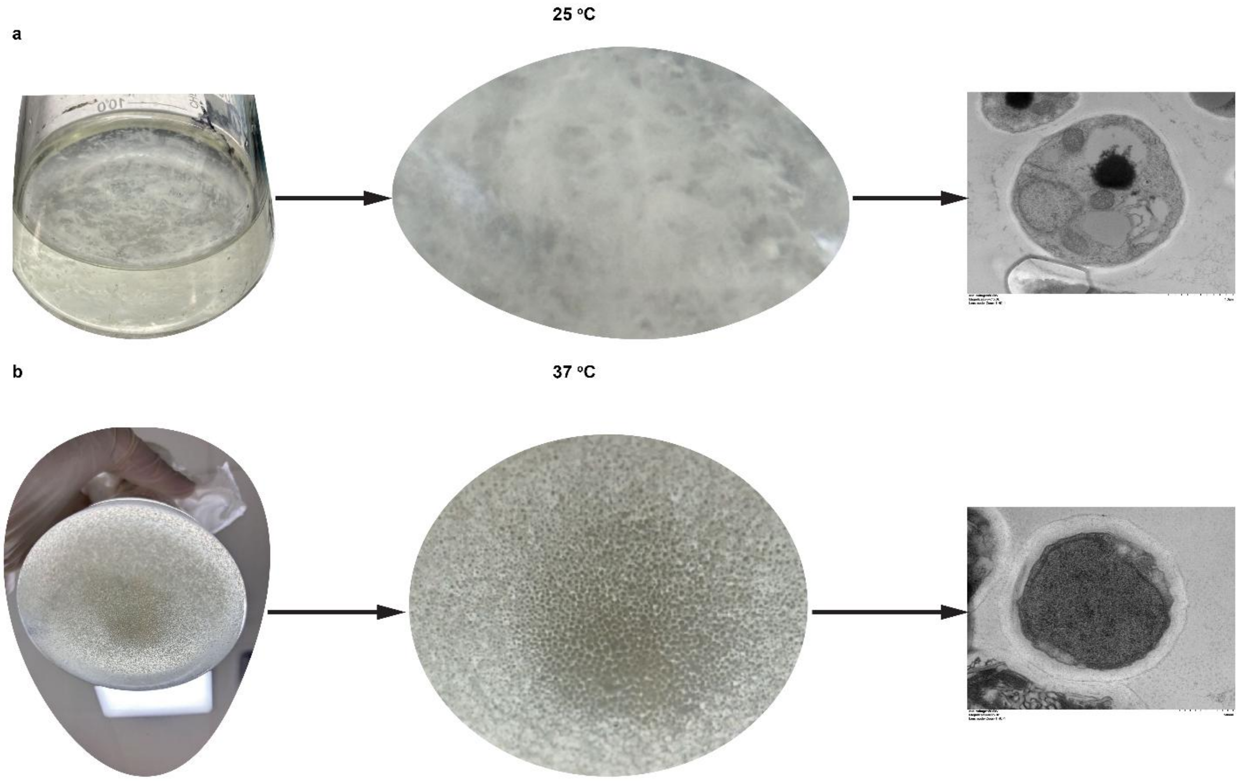

**Supplementary figure 2. Images of *T. marneffei* cells.** (**a**) Images of *T. marneffei* mycelial cells (25°C) were presented in an order from left to right: mycelia in growth medium, zoom-in part of mycelia in the growth medium and images by TEM. (**b**)Images of yeast cells (37°C) were presented in an order from left to right: mycelia in growth medium, zoom-in part of mycelia in the growth medium and images by TEM. The pictures of both mycelia and yeasts in the growth medium were taken using an iPhone 14.

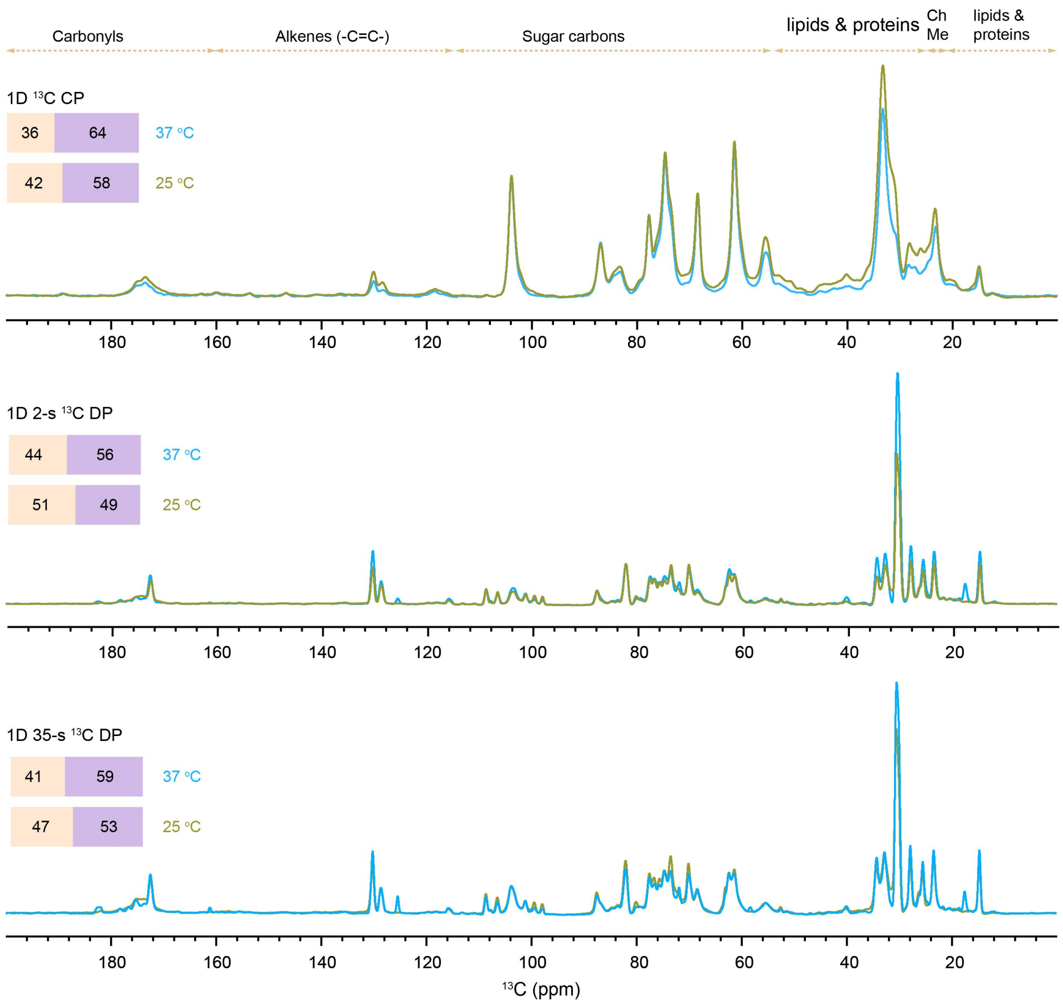

**Supplementary figure 3. 1D $^{13}$C NMR spectra and estimation of the relative polysaccharides to the protein/lipids. Top panel**, 1D $^{13}$C CP spectra for detecting the rigid molecules in both forms of molds and yeasts. **Middle panel**, 1D $^{13}$C DP spectra with 2-s recycle delay preferentially detecting the mobile molecules with fast $^{13}$C $T_1$ relaxation. **Bottom panel**, 1D $^{13}$C DP spectra with 35-s recycle delay specifically for quantitative quantitating all molecules within the intact fungal cells.

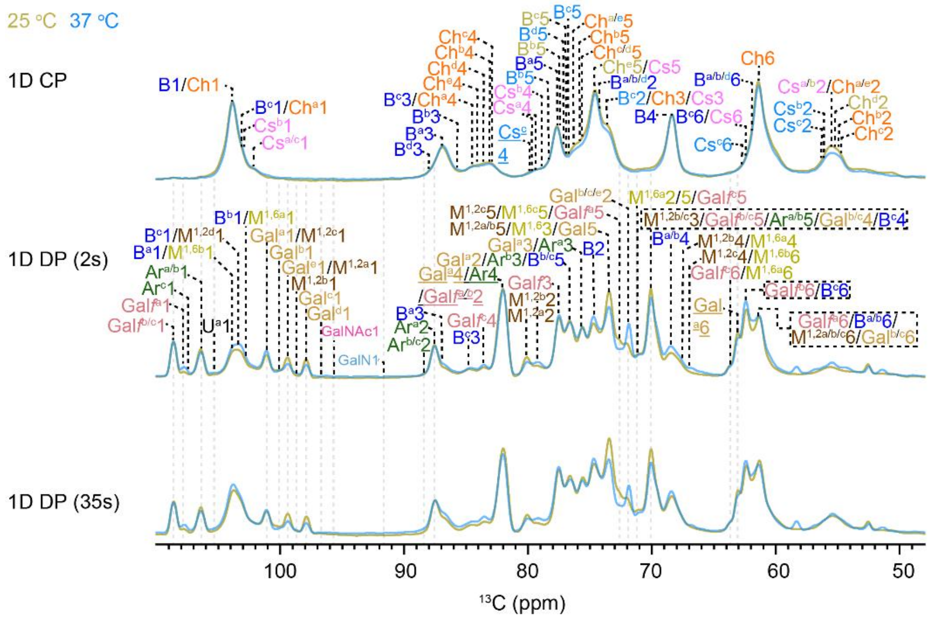

**Supplementary figure 4. 1D $^{13}$C spectra of polysaccharides.** Top-$^{13}$C CP spectrum detects the rigid molecules of the cell walls. Middle-$^{13}$C DP spectrum measured with 2-s recycle delays mobile molecules showing major signals from galfuranose, arabinose, β-1,3-glucose, a-1,2-mannose, a-1,6-mannose and galactose, and minor signals of type-a unknown, N-acetylgalactosamine and galactosamine. Bottom-Quantitative $^{13}$C DP measured with 35-s recycle delays.

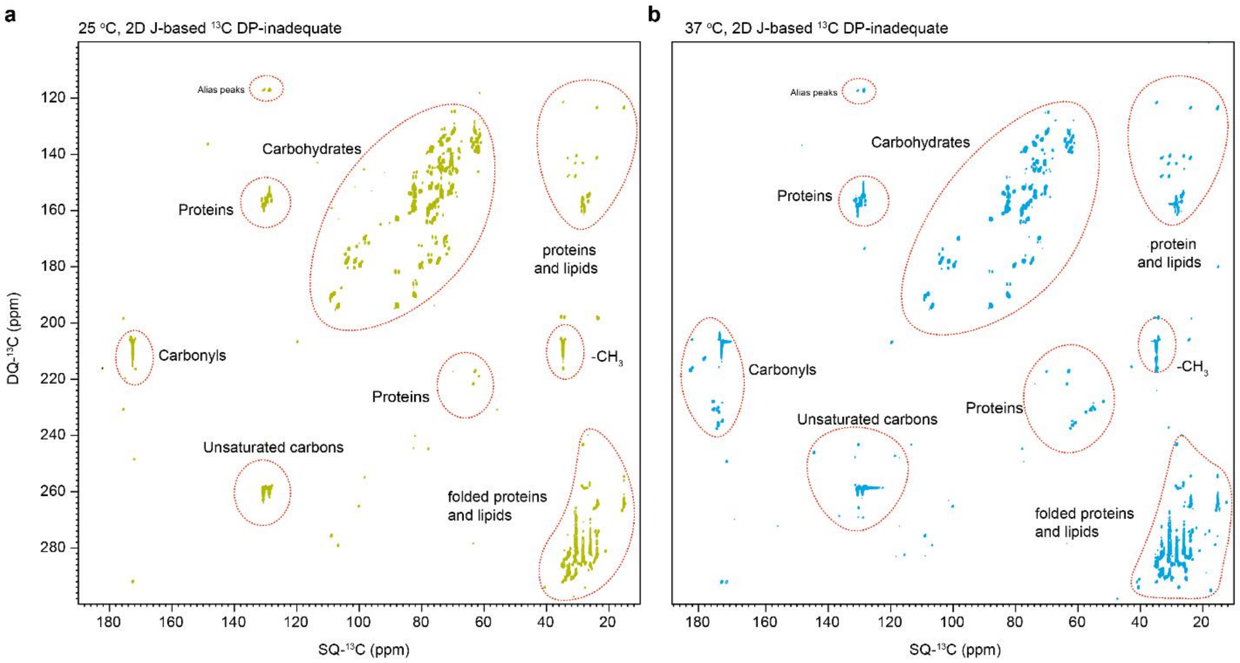

**Supplementary figure 5. 2D DP J-based $^{13}$C INADEQUATE spectra for mobile molecules of *T. marneffei*.** (**a**) and (**b**) Mobile molecules detected by DP-based 2D $^{13}$C refocused J-INADEQUATE spectrum, where the signals of proteins, lipids, and carbohydrates are labeled with red dash circles.

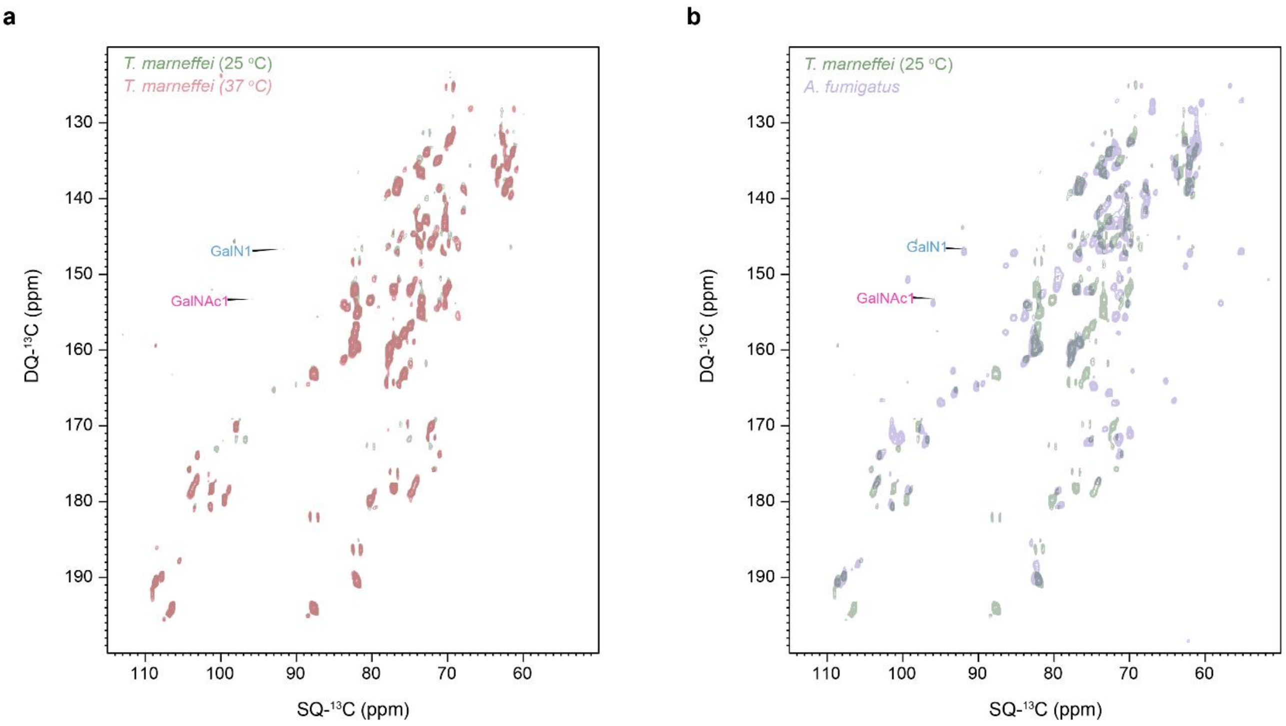

**Supplementary figure 6. Identification of GalN and GalNAc carbohydrates within mobile domain.** (**a**)Overlapping the 2D $^{13}C$ DP J-based INADEQUATE spectra of the molds (25 $^{o}C$) with the one of yeasts (37 $^{o}C$), specifically highlights the C1 signals of GalN and GalNAc within the mobile domain of the molds but are absent in the yeasts. (**b**)Overlapping the 2D $^{13}C$ DP J-based INADEQUATE spectra of the molds with the references one-A. fumigatus, clearly confirm the existence of both GalN and GalNAc carbohydrates. [1,2]

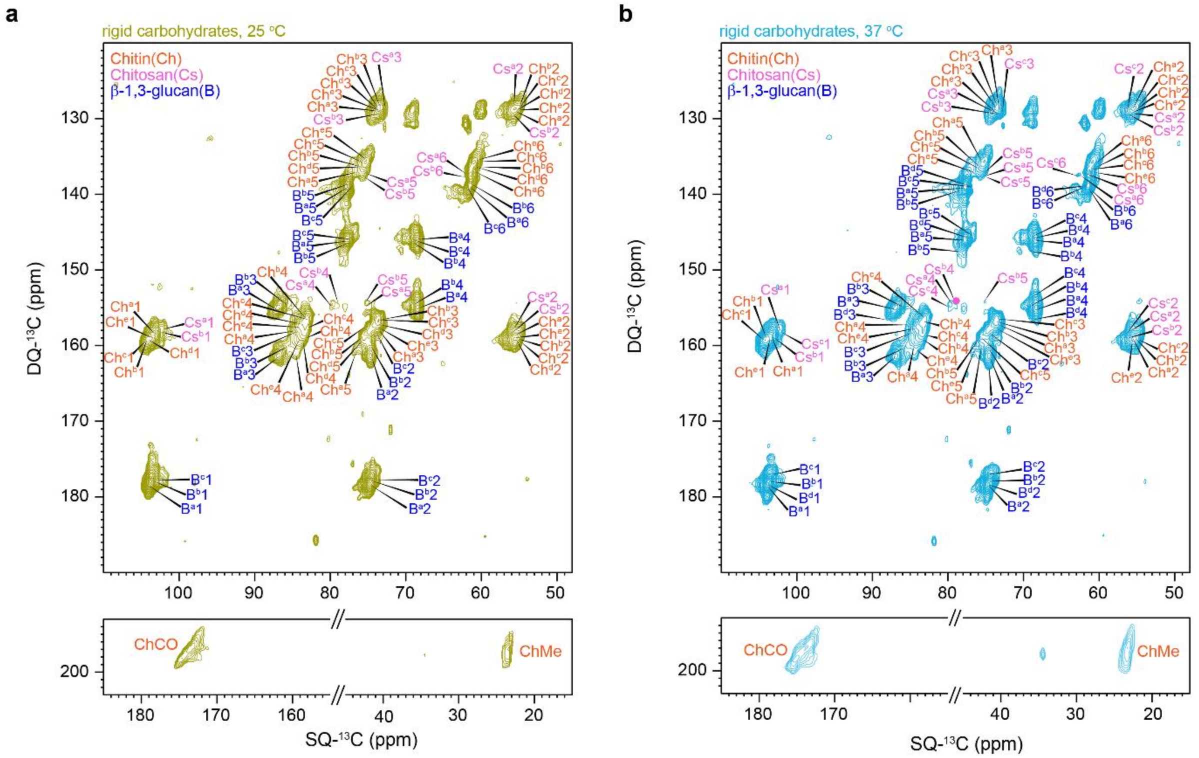

**Supplementary figure 7. Rigid polysaccharides in the *T. mareffei*. (a) & (b)** Through-bond carbon connectivity was determined using 2D $^{13}C$-$^{13}C$ CP refocused J-INADEQUATE spectra, which allowed for the identification of various carbohydrate types.

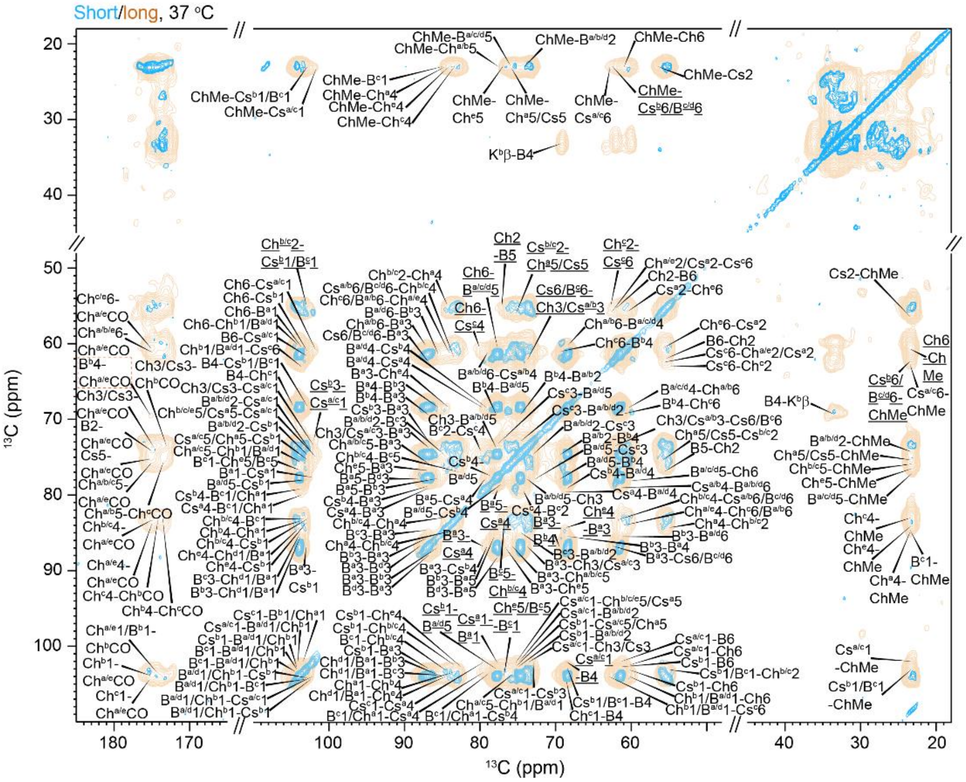

**Supplementary figure 8. Intermolecular interactions observed within the yeasts.** Overlapping of 2D $^{13}$C-$^{13}$C correlation spectra with 53-ms CORD mixing (blue) on 2D $^{13}$C-$^{13}$C PDSD with a very long mixing time of 1 s (orange).

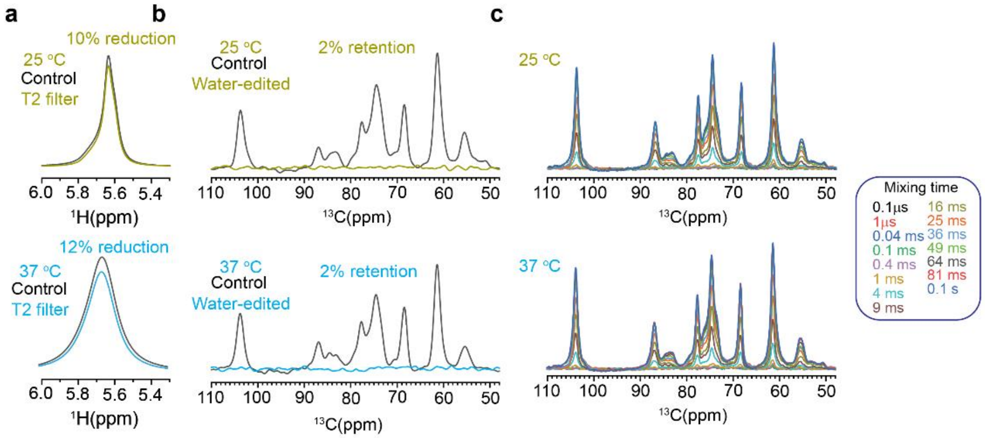

**Supplementary figure 9. Water-edited experiments setup.** (**a**)$^1$H-T$_2$ filtered (gold yellow-25 °C and blue-37 °C) $^1$H NMR spectra, with 10% (25 °C) or 12% (37 °C) of water signal reduction after using $^1$H T$_2$ filter. (**b**)$^1$H-T$_2$ filtered (gold yellow-25 °C and blue-37 °C) and control (black) $^{13}$C spectra are shown for both samples cultured under different temperatures. (**d**) 1D water-edited $^{13}$C spectra with $^1$H spin-diffusion times ranging from 0.1 μs to 0.1 s is presented for both samples (25 °C and 37 °C). All spectra were collected on 700 MHz spectrometer at 15 kHz MAS at 277 K.

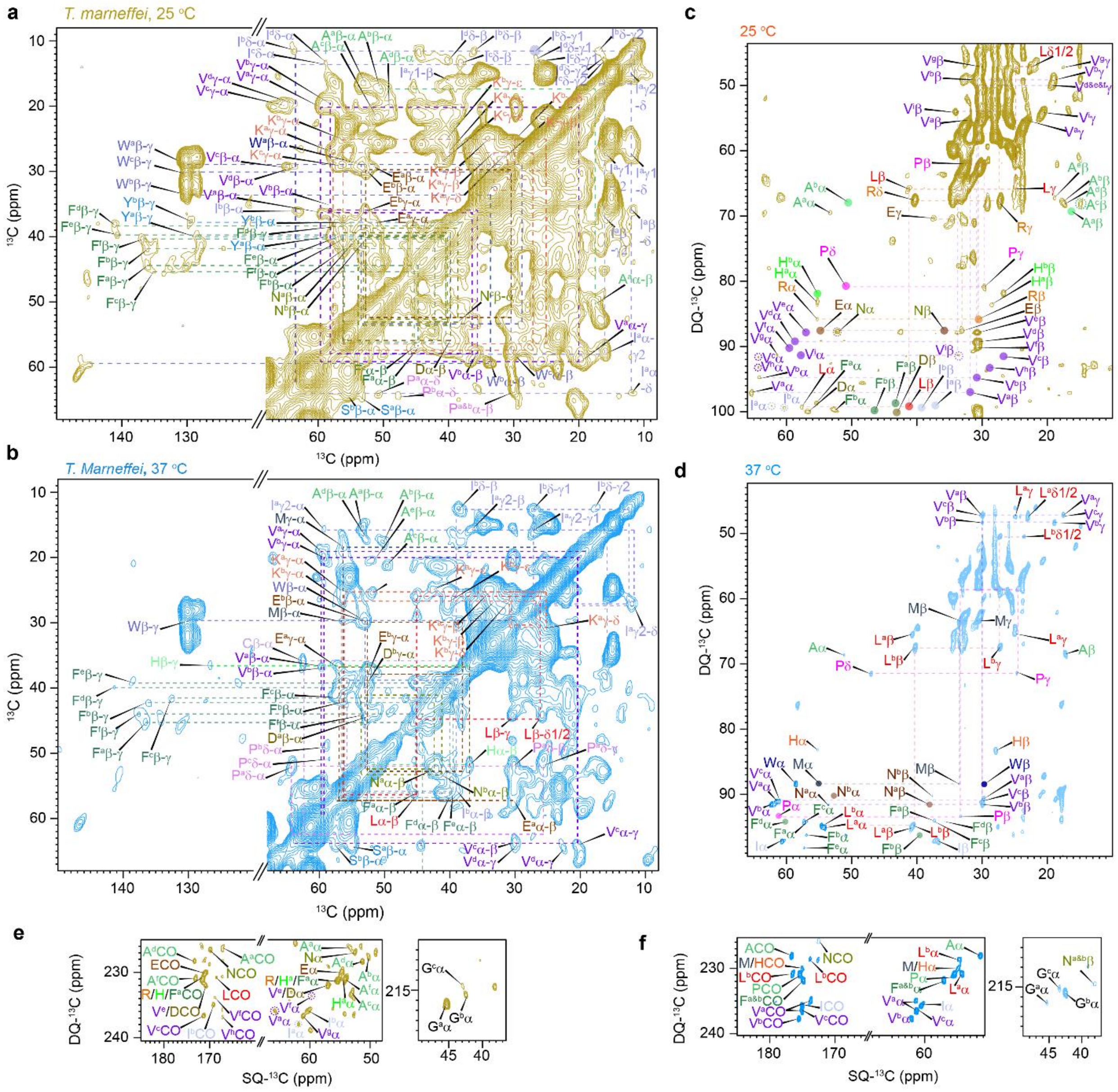

**Supplementary figure 10. Amino acids identified within *T. marneffei*.** (**a**),25 °C and (**b**)37 °C Schematic representations of assignments for the amino acids. (**c**) 25 °C and (**d**)37 °C aliphatic region of amino acids assigned within the mobile domain by 2D $^{13}$C DP J-INADEQUATE spectra. (**e**) 25 °C and (**f**) 37 °C show the correlational peaks (DQ-CO, SQ-Cα) of carbonyl with Cα of mobile amino acids from 2D $^{13}$C DP J-INADEQUATE spectra. Considering the processing parameters (LB -30 Hz and GB 0.03) we used for the protein region of CORD spectra and line broadening, the full width at half maximum (FWHM) was estimated to be 1 ppm for both CORD spectra. Following this logic, two red dashed lines with ±1 ppm inserted into panel c and panel d were used, the chemical shift difference δ of Amino acids are located within this range [-1, 1] ppm, as shown in **Figure 4g** and **4h**, these amino acids take the random coil conformation, while those with δ out of the above range are structured (smaller than -1, strand while larger than 1, α-helical).[2,3]

**Supplementary table 1. The cell-wall thickness of both molds and yeasts.** All measurements were analyzed using imageJ software and the unit is nm. Reading x is abbreviated as Rx, for example, R1 for Reading 1.

| Molds (25 $^oC$) | | | | | | | | | | | | | | | | |
|---|---|---|---|---|---|---|---|---|---|---|---|---|---|---|---|---|
| Rx | R1 | R2 | R3 | R4 | R5 | R6 | R7 | R8 | R9 | R10 | R11 | R12 | R13 | R14 | R15 | Average |
| Cell 1 | 58 | 64 | 62 | 66 | 62 | 62 | 64 | 63 | 62 | 63 | 65 | 67 | 64 | 64 | 64 | |
| Cell 2 | 59 | 74 | 69 | 74 | 56 | 78 | 74 | 68 | 63 | 75 | 85 | 85 | 88 | 82 | 78 | |
| Cell 3 | 49 | 55 | 70 | 52 | 57 | 50 | 57 | 66 | 49 | 53 | 64 | 50 | 48 | 55 | 66 | 67 ± 9 nm |
| Cell 4 | 75 | 68 | 82 | 75 | 54 | 69 | 81 | 68 | 73 | 83 | 75 | 72 | 77 | 73 | 75 | |
| Cell 5 | 71 | 69 | 58 | 66 | 66 | 72 | 66 | 70 | 71 | 68 | 68 | 73 | 58 | 58 | 65 | |
| Cell 6 | 65 | 62 | 73 | 73 | 72 | 67 | 67 | 58 | 61 | 56 | 63 | 60 | 73 | 75 | 69 | |
| Yeasts (37 $^oC$) | | | | | | | | | | | | | | | | |
| Cell 1 | 132 | 153 | 134 | 136 | 147 | 149 | 132 | 150 | 163 | 152 | 157 | 136 | 149 | 127 | 142 | Average |
| Cell 2 | 163 | 160 | 150 | 150 | 164 | 166 | 153 | 170 | 183 | 175 | 192 | 160 | 167 | 198 | 181 | |
| Cell 3 | 139 | 137 | 140 | 133 | 130 | 137 | 166 | 191 | 156 | 138 | 131 | 129 | 178 | 182 | 191 | |
| Cell 4 | 121 | 135 | 113 | 138 | 133 | 154 | 134 | 124 | 116 | 118 | 112 | 111 | 125 | 125 | 135 | 152 ± 21 nm |
| Cell 5 | 134 | 134 | 146 | 166 | 145 | 179 | 174 | 159 | 166 | 175 | 130 | 178 | 176 | 173 | 143 | |
| Cell 6 | 159 | 157 | 150 | 144 | 146 | 183 | 180 | 160 | 153 | 161 | 172 | 177 | 208 | 155 | 157 | |

**Supplementary table 2**. **Solid-state NMR experimental parameters for fungal cell wall characterization.** T = sample temperature; $B_0$ =17.4 T; MAS frequency=15 kHz; ns = number of scans; $d_1$ = recycle delay between scans; $t_{1,\ max}$ = maximum $t_1$ evolution time (for indirect dimension); $t_{1,\ inc}$ = increment for $t_1$ (for indirect dimension) evolution time; $\tau_{dw}$ = dwell time during direct FID acquisition; $\tau_{acq}$ = maximum acquisition time during direct FID detection; $\tau_{xy}$ = cross-polarization contact time during CP from channel X to channel Y; $\nu_{1H,\ dec}$ = dipolar decoupling field strength.

| Experiment | T(K) | NS | d1 (s) | $t_{1,\ max}$ (ms) | $t_{1,\ inc}$ (μs) | $\tau_{dw}$ (μs) | $\tau_{acq}$ (ms) | $\tau_{xy}$ (ms) | $\tau_{mix}$ (ms) | $\nu_{1H\ dec}$ (kHz) | Samples |
|---|---|---|---|---|---|---|---|---|---|---|---|
| 1D $^1H$ | 277 | 4 | 3 | | | 9 | 500 | | | 77.4 | |
| 1D $^{13}C$ CP | 277 | 128 | 2 | | | 7 | 14.33 | 0.5 | | 77.4 | |
| 1D $^{13}C$ DP | 277 | 128 | 2 | | | 11.3 | 20.0 | | | 63.1 | |
| 1D $^{13}C$ DP | 277 | 128 | 35 | | | 11.3 | 20.0 | | | 63.1 | |
| 2D CORD | 277 | 32 | 2 | 7.5 | 25 | 7 | 14.3 | 0.5 | 53 $\tau_{CORD}$ | 63.1 | *T. marneffei* (molds, 25 °C &yeasts, 37 °C) |
| 2D CP J-INADEQUATE | 277 | 32 | 2 | 7.5 | 25 | 7 | 14.3 | 0.5 | | 63.1 | |
| 2D DP J-INADEQUATE | 277 | 32 | 2 | 7.5 | 25 | 7 | 14.3 | | | 63.1 | |
| 2D water-edited | 277 | 32 | 2 | 7.5 | 25 | 7 | 14.3 | 0.5 | 50 $\tau_{PDSD}$ | 77.4 | |
| 1D Torchia $^{13}C$-$T_1$ | 277 | 128 | 2 | | | 7 | 14.3 | 0.5 | | 77.4 | |
| 2D PDSD | 277 | 32 | 2 | 7.5 | 25 | 7 | 14.3 | 0.5 | 1000 $\tau_{PDSD}$ | 77.4 | |

**Supplementary table 3. Estimation of molar composition of cellular components within *T. marneffei*.** The estimation of relative abundance of the carbohydrates and proteins and lipids were done by the integrals of the specific chemical shift ranges of the corresponding types of components, based on their fingerprint chemical shifts. To be specific, the chemical shift ranges used for the integration range from 110 to 52 ppm and 24-20 ppm for carbohydrates and the integrals from chemical shift ranges including 52-24 ppm and 20-0 ppm.

| | Molds (25 °C) | | Yeasts (37 °C) |
|---|---|---|---|
| Rigid | Carbohydrates | 36% | 42% |
| | Lipids and proteins | 64% | 58% |
| Mobile | Carbohydrates | 44% | 51% |
| | Lipids and proteins | 56% | 49% |
| All | Carbohydrates | 41% | 59% |
| | Lipids and proteins | 47% | 53% |

**Supplementary table 4. Molar composition of rigid polysaccharides in cell walls**. The molar percentages of rigid cell-wall polysaccharides are estimated using integrals (volume) of cross peaks in 2D $^{13}$C-$^{13}$C 53-ms CORD. Please note that the total sum of all components does not equal 100% due to rounding of certain values, which result in minor discrepancies.

| Molds (25 $^{o}$C) | | | | Yeasts (37 $^{o}$C) | | | |
|---|---|---|---|---|---|---|---|
| Carbohydrate | | Molar composition% | Total % | Carbohydrate | | Molar composition% | Total % |
| B | B$^{a}$ | 53 ± 5 | 57 ± 5 | B | B$^{a\&d}$ | 68 ± 8 | 72 ± 8 |
| | B$^{b}$ | 1 ± 0.4 | | | B$^{b}$ | 3 ± 0.4 | |
| | B$^{c}$ | 3 ± 0.5 | | | B$^{c}$ | 1 ±0.2 | |
| Ch | Ch$^{a\&e}$ | 10 ± 2 | 31 ± 2 | Ch$^{a}$ | Ch$^{a}$ | 2 ± 0.3 | 21 ± 4 |
| | Ch$^{b\&c\&d}$ | 21 ± 1 | | | Ch$^{b}$ | 15 ± 4 | |
| Cs | Cs$^{a}$ | 7 ± 2 | 11 ± 3 | | Ch$^{c}$ | 2 ± 1 | |
| | Cs$^{b}$ | 4 ± 2 | | | Ch$^{e}$ | 2 ± 0.4 | |
| Errors for the total for both molds and yeast were propagated using the following equations: $\delta f = \sqrt{\sum_{i=1}\left(\left(\frac{\delta f}{\delta x_i}\right)^2 \times (\delta x_i)^2\right)}$, where f is Total and $x_i$ is the specific subtype i of carbohydrates. | | | | Cs | Cs$^{a}$ | 3 ±1 | 8 ± 2 |
| | | | | | Cs$^{b}$ | 2 ± 1 | |
| | | | | | Cs$^{c}$ | 3 ± 1 | |

Peaks used for the estimation of rigid polysaccharides for molds-(1)β-1,3-glucan: B$^{a}$3-5, B$^{a}$3-2, B$^{a}$3-6, B$^{a}$4-3, B$^{b}$3-5, B$^{b}$4-3, B$^{b}$1-3, B$^{c}$4-3, B$^{c}$3-4 and B$^{c}$3-5; (2) chitin: Ch$^{a\&e}$1-CO, Ch$^{a\&e}$2-4, Ch$^{a\&e}$6-4, Ch$^{a\&e}$3-4, Ch$^{b/c/d}$6-4, Ch$^{b/c/d}$3-4, Ch$^{b/c/d}$5-4, Ch$^{b/c/d}$4-5&Ch$^{b/c/d}$4-3; (3)chitosan: Cs$^{a}$4-3, Cs$^{a}$3-4, Cs$^{a}$5-4, Cs$^{b}$5-4, Cs$^{b}$4-5, Cs$^{b}$5-2 and Cs$^{b}$6-4. For the yeasts-(1)b-1,3-glucan: B$^{a\&d}$3-5, B$^{a\&d}$3-2, B$^{a\&d}$4-3, B$^{b}$3-5, B$^{b}$3-4, B$^{b}$4-3, B$^{c}$3-4, B$^{c}$5-3, B$^{c}$4-3; (2) chitin: Ch$^{a}$1-4, Ch$^{a}$4-5, Ch$^{a}$2-4, Ch$^{b}$2-4, Ch$^{b}$4-2, Ch$^{b}$6-4, Ch$^{b}$4-6, Ch$^{c}$6-1, Ch$^{c}$6-2, Ch$^{c}$4-6, Ch$^{e}$4-5, Ch$^{e}$5-4, Ch$^{e}$1-4, Ch$^{e}$4-1; (3)chitosan: Cs$^{a}$2-1, Cs$^{a}$1-2, Cs$^{a}$6-4, Cs$^{b}$5-2, Cs$^{b}$2-1, Cs$^{b}$5-4, Cs$^{c}$4-3, Cs$^{c}$5-4, Cs$^{c}$3-4, Cs$^{c}$1-4.

**Supplementary table 5. Estimation of relative abundance of mobile cell wall polysaccharides.** The molar percentages of mobile cell-wall polysaccharides are estimated by using well-resolved peaks (C1 and C2) of 2D $^{13}C$ DP J-based INADEQUATE spectra.

| Mobile polysaccharides | | | | |
|---|---|---|---|---|
| | Molds, 25 °C | | Yeasts, 37 °C | |
| Polysacharide | % | Peak | % | Peak |
| N-acetylgalactosamine | 0.2 | C1, C2 | n/a | n/a |
| Galactosamine | 0.3 | C1, C2 | n/a | n/a |
| Galactose | 24.6 | C1, C2 | 21.8 | C1, C2 |
| Galfuranose | 26.1 | C1, C2 | 28.1 | C1 |
| α-1,6-mannose | 2.1 | C1, C2 | 2.2 | C1, C2 |
| α-1,2-mannose | 14.1 | C1, C2 | 13.1 | C1, C2 |
| β-1,3-glucan | 5.5 | C1, C2 | 9.1 | C1, C2 |
| Arabinose | 23.9 | C1, C2 | 21.0 | C1, C2 |
| F | 0.5 | C1 | 1.9 | C5, C6 |
| Unknown | 2.7 | C1, C2 | 2.7 | C1, C2 |

**Supplementary table 6. Relaxation dynamics of rigid polysaccharides by 1D $^{13}$C Torchia CP spectra.** The normalized data (I(t)) were fitted to a single-exponential equation, $I(t) = \exp(-t/T_1)$, to determine the $^{13}$C-$T_1$ constants, which reflect the dynamics of the corresponding carbohydrate polymers. The unit of $T_1$ is seconds (s). The errors for the individual $T_1$ values were derived from the fitting equation, while the errors for the average $T_1$ were calculated using the standard deviation function STDEV.P in Excel.

| | | | Molds (25 °C) | | | | Yeasts (37 °C) | | |
|---|---|---|---|---|---|---|---|---|---|
| Carbohydrate | | $T_1$ | R | Average $T_1$ | Carbohydrates | | $T_1$ | $R^2$ | Average $T_1$ |
| | Ch$^{b\&c}$1 | 7.46 ± 0.24 | 0.99 | | | Ch$^{c}$1 | 5.46 ± 0.44 | 0.94 | |
| | Ch$^{a\&c}$4 | 6.86 ± 0.53 | 0.94 | | | Ch$^{b\&c}$4 | 9.64 ± 0.60 | 0.95 | |
| Ch | Ch$^{b\&c}$4 | 10.26 ± 0.32 | 0.99 | 7.95 ± 2.32 | Ch | Ch$^{a\&c}$4 | 6.44 ± 0.66 | 0.88 | 6.97 ± 2.20 |
| | Ch$^{d}$4 | 10.27 ± 0.36 | 0.99 | | | Ch$^{b\&c}$2 | 8.83 ± 0.56 | 0.96 | |
| | Ch$^{a\&b\&d}$5 | 4.88 ± 0.39 | 0.95 | | | Ch$^{b\&c}$5 | 4.47 ± 0.45 | 0.91 | |
| | Cs$^{a}$1 | 2.63 ± 0.24 | 0.96 | | | Cs$^{a\&c}$1 | 2.58 ± 0.25 | 0.96 | |
| | Cs$^{b}$1 | 2.23 ± 0.19 | 0.96 | | | Cs$^{c}$3 | 4.18 ± 0.43 | 0.92 | |
| Cs | Cs$^{a}$4 | 2.18 ± 0.21 | 0.95 | 2.57 ± 0.38 | Cs | Csa4 | 1.93 ± 0.26 | 0.89 | |
| | Cs$^{b}$4 | 2.68 ± 0.22 | 0.96 | | | Cs$^{b}$4 | 1.98 ± 0.26 | 0.90 | 2.84 ± 1.02 |
| | Csa3 | 3.12 ± 0.28 | 0.95 | | | Cs$^{b}$4 | 2.32 ± 0.29 | 0.90 | |
| B | B$^{a}$3 | 4.05 ± 0.14 | 0.99 | | | Cs$^{a\&c}$5 | 4.07 ± 0.22 | 0.98 | |
| | B$^{b}$3 | 4.05 ± 0.20 | 0.99 | | | B$^{d}$3 | 4.13 ± 0.20 | 0.97 | |
| | B$^{a\&b}$5 | 3.74 ± 0.07 | 0.99 | 3.74 ± 0.45 | B | B$^{a}$3 | 4.16 ± 0.09 | 0.99 | |
| | Ba&b2 | 4.12 ± 0.14 | 0.99 | | | B$^{b}$3 | 4.23 ± 0.16 | 0.99 | |
| | B$^{a}$4 | 3.53 ± 0.06 | 0.99 | | | B$^{a\&d}$5 | 3.90 ± 0.07 | 0.99 | 4.01 ± 0.29 |
| | B$^{b\&c}$4 | 2.94 ± 0.10 | 0.99 | | | B6 | 3.47 ± 0.13 | 0.99 | |
| | | | | | | B$^{a.b\&d}$2 | 4.19 ± 0.13 | 0.99 | |

**Supplementary table 7. Water-hydration dynamics of polysaccharides.** The pre-normalized data was fitted to the single exponential equation, y=1-A*exp(-x/T), where x is the square root of mixing time and T is the buildup time constant.

| | $\sqrt{\text{Mixing time}}$ ($\sqrt{\text{ms}}$) | 25 °C (molds) | | | 37 °C (yeasts) | | |
|---|---|---|---|---|---|---|---|
| | | B[a]3 | Ch[b&c]4 | Cs[a]4 | | | |
| Data | 0.1E-7 | 1.72106E-4 | 0.00458 | 0.00499 | 0.0032 | 8.53137E-4 | 0.00141 |
| | 0.03162 | 0.00128 | 0.00959 | 0.00645 | 0.00358 | 0.00932 | 0.04155 |
| | 0.2 | 0.00176 | 0.01087 | 0.04353 | 0.00615 | 0.02543 | 0.02043 |
| | 0.31623 | 0.00243 | 0.01469 | 0.06312 | 0.01793 | 0.02929 | 0.02922 |
| | 0.63246 | 0.00871 | 0.02441 | 0.07096 | 0.02613 | 0.03349 | 0.07493 |
| | 1 | 0.0336 | 0.03423 | 0.083 | 0.07062 | 0.04285 | 0.08865 |
| | 2 | 0.17685 | 0.13094 | 0.154 | 0.31911 | 0.11534 | 0.26941 |
| | 3 | 0.39567 | 0.23079 | 0.26567 | 0.61805 | 0.29341 | 0.49594 |
| | 4 | 0.54549 | 0.27994 | 0.38014 | 0.79508 | 0.38833 | 0.57661 |
| | 5 | 0.70565 | 0.50713 | 0.47776 | 0.92333 | 0.56779 | 0.76722 |
| | 6 | 0.81406 | 0.61068 | 0.6735 | 0.97 | 0.76249 | 0.8698 |
| | 7 | 0.95709 | 0.72332 | 0.83497 | 0.98706 | 0.82181 | 1 |
| | 8 | 0.97876 | 0.89191 | 0.90184 | 1 | 0.93214 | 0.9821 |
| | 9 | 1 | 0.96967 | 1 | 0.99189 | 0.96602 | 0.99118 |
| | 10 | 0.98459 | 1 | 0.99636 | 0.95646 | 1 | 0.90231 |
| A | | 1.00426 ± 0.01131 | 0.99957 ± 0.0127 | 0.99782 ± 0.0117 | 1.00091 ± 0.01063 | 1.0029 ± 0.01178 | 1.00125 ± 0.00902 |
| $R^2$ | | 0.96629 | 0.94165 | 0.9547 | 0.97382 | 0.95319 | 0.97781 |
| T | | 4.296 ± 0.50912 | 6.10499 ± 0.78832 | 5.49133 ± 0.65573 | 3.09295 ± 0.37914 | 5.2366 ± 0.62891 | 3.8281 ± 0.3732 |
| Average T | | 5.30±0.44 | | | 4.05±0.23 | | |

**Supplementary table 8. Site-specific water-hydration analysis of *T. marneffei*.** Water-hydration experiments were conducted using 2D 13C-13C water-edited DARR, where 1D slices of both rows and columns were extracted from 2D experiments (Control and water-edited). The site-specific water-hydration levels were done by obtaining the peak intensity of the corresponding carbon sites from the above extracted 1D slices (S1 from the water-edited spectra and S0 from the control spectra) and S1/S0 ratios were calculated. No error bar was included, since we use the statistical analysis for all the data in a violin plot (**Fig. 5c**).

| Molds, 25 °C | | | | | | | | Yeasts, 37 °C | | | | | | | |
|---|---|---|---|---|---|---|---|---|---|---|---|---|---|---|---|
| Peaks | S0 | S1 | S1/S0 | Peaks | S0 | S1 | S1/ | Peaks | S0 | S1 | S1/S | Peaks | S0 | S1 | S1/S0 |
| **Ch$^{a}$6-6** | 36855033344 | 4905882944 | 0. 13 | Ch$^{d}$3-4 | 36362041344 | 6067860608 | 0.1 | **Ch$^{a}$4-4** | 5985226240 | 880556544 | 0.15 | Ch$^{e}$5-2 | 632612352 | 230830528 | 0.36 |
| Ch$^{a}$6-5 | 4625916928 | 559039488 | 0. 12 | Ch$^{d}$1-4 | 12460402688 | 3470270208 | 0.2 | Ch$^{a}$2-4 | 1896804864 | 323430720 | 0.17 | Ch$^{e}$4-2 | 514458624 | 290740160 | 0.57 |
| Ch$^{a}$6-3 | 3427055104 | 533022208 | 0. 16 | Ch$^{d}$5-4 | 9800513536 | 1575652096 | 0.1 | Ch$^{a}$1-4 | 1008990720 | 210395904 | 0.21 | **Cs$^{a/c}$1-1** | 6132244992 | 1949338816 | 0.32 |
| Ch$^{a}$6-1 | 2120597504 | 364916352 | 0. 17 | Ch$^{d}$2-4 | 4553512448 | 843069760 | 0.1 | Ch$^{a}$5-4 | 703969792 | 104628160 | 0.15 | Cs$^{a/c}$2-1 | 1543155456 | 469941120 | 0.30 |
| Ch$^{a}$6-2 | 2062242816 | 291564864 | 0. 14 | Ch$^{d}$6-4 | 4537210880 | 781300480 | 0.1 | Ch$^{a}$2-4 | 598902784 | 149368192 | 0.25 | Cs$^{a/c}$5-1 | 1351617536 | 517143296 | 0.38 |
| Ch$^{a}$6-4 | 953424384 | 102610112 | 0. 11 | **Ch$^{d}$4-4** | 4023895040 | 474488640 | 0.1 | Ch$^{a}$6-4 | 164583680 | 19363648 | 0.12 | Cs$^{a/c}$3-1 | 1332981760 | 571374464 | 0.43 |
| **Ch$^{a}$4-4** | 8209811968 | 1034263296 | 0. 13 | Ch$^{e}$4-3 | 7436725760 | 1058109376 | 0.1 | **Ch$^{a}$3-3** | 13104768768 | 2373621056 | 0.18 | Cs$^{a/c}$6-1 | 273279232 | 77588480 | 0.28 |
| Ch$^{a}$4-3 | 3955160576 | 534174016 | 0.14 | Ch$^{e}$4-5 | 4115637248 | 595587456 | 0.1 | Ch$^{a}$3-1 | 4584319488 | 973543040 | 0.21 | Cs$^{a/c}$4-1 | 259423744 | 97493952 | 0.38 |
| Ch$^{a}$4-1 | 1477533696 | 513846720 | 0.35 | Ch$^{e}$4-1 | 2622626816 | 447371648 | 0.1 | Ch$^{a}$3-5 | 4119152896 | 1263093760 | 0.31 | **Cs$^{a/c}$1-1** | 4754300928 | 740301952 | 0.16 |
| Ch$^{a}$4-5 | 1037663744 | 90250816 | 0.09 | Ch$^{e}$4-2 | 1573936640 | 381542464 | 0.2 | Ch$^{a}$3-2 | 2639940096 | 434499200 | 0.16 | Cs$^{a/c}$1-2 | 1098003200 | 365947584 | 0.33 |
| Ch$^{a}$4-6 | 1035852288 | 80959232 | 0.08 | **Ch$^{e}$4-4** | 1294853632 | 56275136 | 0.0 | Ch$^{a}$3-4 | 1706862592 | 283243968 | 0.17 | Cs$^{a/c}$1-5 | 910612736 | 570503360 | 0.63 |
| Ch$^{a}$4-2 | 843366400 | 145143360 | 0.17 | Ch$^{e}$4-6 | 1074193920 | 190693184 | 0.1 | Ch$^{a}$3-6 | 1108451584 | 319366336 | 0.29 | Cs$^{a/c}$1-3 | 1133740288 | 538444928 | 0.47 |
| **Ch$^{c}$1-1** | 10759939584 | 3028756672 | 0.28 | Ch$^{e}$3-4 | 35450126336 | 84232192 | 0.0 | Ch$^{b&c}$4- | 20771361536 | 3671484096 | 0.18 | Cs$^{a/c}$1-6 | 99573120 | 45339008 | 0.46 |
| Ch$^{c}$1-3 | 5375842816 | 1890289408 | 0.35 | Ch$^{e}$5-4 | 19327130112 | 54582720 | 0.0 | Ch$^{b&c}$4- | 6282043648 | 2032691456 | 0.32 | Cs$^{a/c}$1-4 | 153847424 | 65363776 | 0.42 |
| Ch$^{c}$1-2 | 2599040512 | 857124608 | 0.33 | Ch$^{e}$1-4 | 11684693504 | 53639488 | 0.0 | Ch$^{b&c}$4- | 3779265792 | 1277502784 | 0.34 | **Cs$^{b}$5-5** | 19096168448 | 4906774528 | 0.26 |
| Ch$^{c}$1-5 | 2423418368 | 821394368 | 0.34 | Ch$^{e}$2-4 | 4936707072 | 126761728 | 0.0 | Ch$^{b&c}$4- | 2865079552 | 488811840 | 0.17 | Cs$^{b}$5-3 | 9087217152 | 1342921152 | 0.15 |
| Ch$^{c}$1-4 | 1226811904 | 330968832 | 0.27 | **Ch$^{e}$4-4** | 4115637248 | 85182656 | 0.0 | **Ch$^{b&c}$4-** | 2122803712 | 533978944 | 0.25 | Cs$^{b}$5-1 | 4695653120 | 1619926336 | 0.34 |
| Ch$^{c}$1-6 | 698720768 | 152227200 | 0.22 | Ch$^{e}$6-4 | 3479185408 | 133117632 | 0.1 | Ch$^{b&c}$4- | 1593697792 | 492585664 | 0.31 | Cs$^{b}$5-4 | 2943149056 | 574724160 | 0.20 |
| **Ch$^{c}$4-4** | 5246353408 | 949796160 | 0.18 | **Cs$^{a}$3-3** | 21793788416 | 3688160576 | 0.1 | Ch$^{b&c}$3- | 2149010944 | 583744064 | 0.27 | Cs$^{b}$5-6 | 2522229760 | 785779136 | 0.31 |
| Ch$^{c}$3-4 | 3492264960 | 587471040 | 0.17 | Cs$^{a}$5-3 | 8260887552 | 1276345664 | 0.1 | Ch$^{b&c}$2- | 1121169920 | 285380992 | 0.25 | Cs$^{b}$5-2 | 1146587904 | 306300672 | 0.27 |
| Ch$^{c}$5-4 | 1971487744 | 594873536 | 0.30 | Cs$^{a}$2-3 | 4427164672 | 409528896 | 0.0 | Ch$^{b&c}$5- | 1027790080 | 351437184 | 0.34 | Cs$^{b}$4-4 | 5481831168 | 1849653312 | 0.34 |
| Ch$^{c}$2-4 | 1901864448 | 317148480 | 0.17 | Cs$^{a}$1-3 | 2374294016 | 176047040 | 0.0 | Ch$^{b&c}$1- | 743576320 | 420650624 | 0.57 | Cs$^{b}$4-5 | 2705022976 | 649215616 | 0.24 |
| Ch$^{c}$6-4 | 1312967168 | 466118720 | 0.36 | Cs$^{a}$4-3 | 1805615104 | 207117248 | 0.1 | Ch$^{b&c}$6- | 631927040 | 147056576 | 0.23 | Cs$^{b}$4-6 | 1270019840 | 421503808 | 0.33 |
| Ch$^{c}$1-4 | 1300810752 | 320923392 | 0.25 | Cs$^{a}$6-3 | 1508412416 | 371134848 | 0.2 | **Ch$^{e}$4-4** | 4997274624 | 751389952 | 0.15 | Cs$^{b}$4-3 | 791375360 | 289074560 | 0.37 |
| **Ch$^{d}$4-4** | 5924534784 | 1185545280 | 0.20 | Cs$^{a}$4-1 | 11421707264 | 1237061312 | 0.1 | Ch$^{e}$4-3 | 2351052544 | 469113664 | 0.20 | Cs$^{b}$4-1 | 434051840 | 151535168 | 0.35 |
| Ch$^{d}$4-3 | 3989028864 | 729495232 | 0.18 | Cs$^{a}$4-2 | 2669432832 | 306533632 | 0.1 | Ch$^{e}$4-5 | 943420416 | 227273856 | 0.24 | Cs$^{b}$4-2 | 233698304 | 67635136 | 0.29 |
| Ch$^{d}$4-5 | 1996848640 | 459403584 | 0.23 | Cs$^{a}$4-5 | 2306565632 | 288076672 | 0.1 | Ch$^{e}$4-2 | 577626880 | 292987008 | 0.51 | B$^{a}$3-4 | 4541873664 | 2257583232 | 0.50 |
| Ch$^{d}$4-2 | 1672254464 | 537030912 | 0.32 | Cs$^{a}$4-3 | 1888014336 | 118507136 | 0.0 | Ch$^{e}$4-1 | 576756736 | 218882240 | 0.38 | **B$^{a}$3-3** | 4535391232 | 2035843200 | 0.45 |
| Ch$^{d}$4-6 | 1596031488 | 257516672 | 0.16 | **Cs$^{a}$4-4** | 325914112 | 79649216 | 0.2 | Ch$^{e}$4-6 | 549830144 | 237685056 | 0.43 | B$^{a}$3-2 | 4411572736 | 2466743936 | 0.56 |
| Ch$^{d}$4-1 | 1581721600 | 381317824 | 0.24 | **Cs$^{b}$1-1** | 14632494592 | 1332601152 | 0.0 | **Ch$^{e}$2-2** | 6300017408 | 1316341248 | 0.21 | B$^{a}$3-1 | 4380860672 | 2152986432 | 0.49 |
| **Ch4-4** | 5278868992 | 981762816 | 0.19 | Cs$^{b}$1-3 | 5899266560 | 409468672 | 0.0 | Ch$^{e}$3-2 | 2675523584 | 1065311232 | 0.40 | B$^{a}$3-5 | 2627035904 | 1438298432 | 0.55 |
| Ch3-4 | 3443716608 | 602818880 | 0.18 | Cs$^{b}$1-2 | 2929144832 | 112271168 | 0.0 | Ch$^{e}$1-2 | 1911836416 | 483709952 | 0.25 | B$^{a}$3-6 | 1894801152 | 987743872 | 0.52 |

| | | | |
|---|---|---|---|
| Ch5-4 | 2045461504 | 279079936 | 0.14 |
| Ch2-4 | 1417739776 | 257002880 | 0.18 |
| Ch6-4 | 676225024 | 166671296 | 0.25 |
| **B$^{a}$3-3** | 6181319680 | 2267094976 | 0.37 |
| B$^{a}$3-2 | 5845373952 | 2376354944 | 0.41 |
| B$^{a}$3-1 | 5697375232 | 2141579264 | 0.38 |
| B$^{a}$3-4 | 5615305728 | 2199581440 | 0.39 |
| B$^{a}$3-5 | 3412302848 | 1438125184 | 0.42 |
| B$^{a}$3-6 | 2266461184 | 1101475712 | 0.49 |
| B$^{a}$4-3 | 8939674112 | 2175538112 | 0.24 |
| **B$^{a}$3-3** | 6517389824 | 2259386368 | 0.35 |
| B$^{a}$6-3 | 6300937216 | 1721008512 | 0.27 |
| B$^{a}$2-3 | 5348424192 | 2414386560 | 0.45 |
| B$^{a}$1-3 | 4845070336 | 2227968000 | 0.46 |
| B$^{a}$5-3 | 4804809216 | 1486899456 | 0.31 |
| B$^{a}$1-4 | 25260517376 | 6412052480 | 0.25 |
| B$^{a}$2-4 | 13675988480 | 4401129856 | 0.32 |
| **B$^{a}$4-4** | 8728842752 | 2034196544 | 0.23 |
| B$^{a}$6-4 | 7145206784 | 1458249920 | 0.20 |
| B$^{a}$3-4 | 6042735616 | 2151735936 | 0.36 |
| B$^{a}$5-4 | 5537737728 | 1881439680 | 0.34 |
| **B$^{b}$4-4** | 22059845120 | 3752731328 | 0.17 |
| B$^{b}$4-5 | 7841965056 | 2024048000 | 0.26 |
| B$^{b}$4-6 | 7751483904 | 1542473728 | 0.20 |
| B$^{b}$4-2 | 7749987328 | 1710960320 | 0.22 |
| B$^{b}$4-1 | 5062482944 | 775784576. | 0.15 |
| B$^{b}$4-3 | 2880207872 | 569690816. | 0.20 |
| **B$^{b}$4-4** | 3110911488 | 649893184. | 0.21 |
| B$^{b}$5-4 | 1497900544 | 630080512. | 0.42 |
| B$^{b}$6-4 | 1770734592 | 576858624. | 0.33 |
| B$^{b}$2-4 | 2122608640 | 716131584. | 0.34 |
| B$^{b}$1-4 | 1553521664 | 711005696. | 0.46 |
| B$^{b}$3-4 | 2860043264 | 462991744. | 0.16 |
| **B$^{c}$4-4** | 22141775872 | 9647097920 | 0.44 |
| B$^{c}$4-6 | 8328671744 | 2411037376 | 0.29 |
| B$^{c}$4-5 | 7390074368 | 4131289856 | 0.56 |
| B$^{c}$4-2 | 5640973824 | 2581880512 | 0.46 |
| B$^{c}$4-1 | 3923888640 | 1507139008 | 0.38 |
| B$^{c}$4-4 | 1758325760 | 936764800. | 0.53 |

| | | | |
|---|---|---|---|
| Cs$^{b}$1-5 | 2624409600 | 250633856 | 0.2 |
| Cs$^{b}$1-6 | 484864512 | 63823808 | 0.2 |
| Cs$^{b}$1-4 | 315267584 | 25551936 | 0.2 |
| **Cs$^{b}$4-4** | 7950607360 | 1228826816 | 0.1 |
| Cs$^{b}$4-5 | 3849069056 | 561369856 | 0.1 |
| Cs$^{b}$4-3 | 1898413568 | 462713984 | 0.2 |
| Cs$^{b}$4-6 | 911540224 | 166834112 | 0.1 |
| Cs$^{b}$4-1 | 480851968 | 69469120 | 0.1 |
| Cs$^{b}$4-2 | 309264896 | 36833600 | 0.1 |
| **Cs$^{b}$4-4** | 7950607360 | 1173481088 | 0.1 |
| Cs$^{b}$5-4 | 3873024512 | 380325696 | 0.1 |
| Cs$^{b}$3-4 | 2372521984 | 518888064 | 0.2 |
| Cs$^{b}$2-4 | 125226496 | 22365952 | 0.1 |
| **Cs$^{b}$3-3** | 26226012160 | 5249159360 | 0.2 |
| Cs$^{b}$3-5 | 25864086016 | 4417205312 | 0.1 |
| Cs$^{b}$3-4 | 3852422144 | 460169472 | 0.1 |
| Cs$^{b}$3-1 | 3788994560 | 682389632 | 0.1 |
| Cs$^{b}$3-6 | 2880807424 | 615794752 | 0.2 |
| Cs$^{b}$3-2 | 2568498176 | 700555392 | 0.2 |
| B$^{c}$3-4 | 3822880768 | 1021666880. | 0.2 |
| **B$^{c}$4-4** | 1712077824 | 800710016.0 | 0.4 |
| B$^{c}$1-4 | 1419469312 | 421436224.0 | 0.3 |
| B$^{c}$5-4 | 838302208 | 332081088.0 | 0.4 |
| B$^{c}$6-4 | 763390464 | 402799104.0 | 0.5 |

| | | | |
|---|---|---|---|
| Ch$^{c}$6-2 | 955119616 | 276342144 | 0.29 |
| B$^{a}$6-3 | 4667180032 | 1761440384 | 0.38 |
| **B$^{a}$3-3** | 4548684288 | 2093492032 | 0.46 |
| B$^{a}$2-3 | 3624274688 | 2123296768 | 0.59 |
| B$^{a}$5-3 | 3553212672 | 1369377856 | 0.39 |
| B$^{a}$1-3 | 3423176448 | 2132143744 | 0.62 |
| B$^{b}$4-3 | 1986480896 | 952506240 | 0.48 |
| **B$^{b}$3-3** | 1957012480 | 723234624 | 0.37 |
| B$^{b}$2-3 | 1622188800 | 847864448 | 0.52 |
| B$^{b}$6-3 | 1428670720 | 521171072 | 0.36 |
| B$^{b}$1-3 | 1100779264 | 456496448 | 0.41 |
| B$^{b}$5-3 | 826657792 | 605679808 | 0.73 |
| B$^{b}$3-2 | 2328434944 | 847864448 | 0.36 |
| **B$^{b}$3-3** | 1954776832 | 923234624 | 0.47 |
| B$^{b}$3-1 | 1792262912 | 556496448 | 0.31 |
| B$^{b}$3-4 | 1215607040 | 593000576 | 0.49 |
| B$^{b}$3-6 | 673558784 | 421171072 | 0.63 |
| B$^{b}$3-5 | 493139712 | 275199104 | 0.56 |
| B$^{c}$4-3 | 5616490240 | 1897618624 | 0.34 |
| B$^{c}$4-2 | 2182974464 | 786072384 | 0.36 |
| **B$^{c}$4-4** | 1380013056 | 249585920 | 0.18 |
| B$^{c}$4-1 | 674448640 | 342787200 | 0.51 |
| B$^{c}$4-5 | 566169088 | 124005312 | 0.22 |
| B$^{c}$4-6 | 301679104 | 181819584 | 0.60 |
| B$^{c}$3-4 | 5677826304 | 1170834496 | 0.21 |
| B$^{c}$2-4 | 2308989952 | 550888832 | 0.24 |
| **B$^{c}$4-4** | 1055041280 | 388455744 | 0.37 |
| B$^{c}$1-4 | 1015106560 | 367090304 | 0.36 |
| B$^{c}$5-4 | 653271808 | 207945920 | 0.32 |
| B$^{c}$6-4 | 159702016 | 30216576 | 0.19 |
| B$^{d}$4-3 | 1903741184 | 837204608 | 0.44 |
| **B$^{d}$3-3** | 1027212288 | 778838656 | 0.76 |
| B$^{d}$2-3 | 957295104 | 614772288 | 0.64 |
| B$^{d}$5-3 | 926213632 | 624657600 | 0.67 |
| B$^{d}$1-3 | 839305728 | 578839680 | 0.69 |
| B$^{d}$6-3 | 701141760 | 552518400 | 0.79 |
| **B$^{d}$5-5** | 8994761216 | 5907270272 | 0.66 |
| B$^{d}$5-6 | 8380498176 | 3939309760 | 0.47 |
| B$^{d}$5-4 | 6042315264 | 2599782848 | 0.43 |

| | | | |
|---|---|---|---|
| B$^{a}$4-3 | 6655787520 | 2526409664 | 0.38 |
| B$^{d}$5-2 | 43833797 | 2163313472 | 0.49 |
| B$^{d}$5-1 | 30743884 | 1361040128 | 0.44 |
| B$^{d}$5-3 | 91150848 | 581448384 | 0.64 |

**Supplementary table 9. chemical shifts of rigid polysaccharides in *T. marneffei* cell walls**. Superscripts are used to denote different allomorphs. Not applicable (/). Unidentified (-). Minor forms (m).

| Carbohydrate | | C1 | C2 | C3 | C4 | C5 | C6 | CO | $CH_3$ | Experiments | References |
|---|---|---|---|---|---|---|---|---|---|---|---|
| *T. marneffei*, 25 °C | | | | | | | | | | | |
| chitin | a | 103.1 | 55.5 | 73.5 | 84.5 | 76.5 | 60.6 | 174.9 | 22.9 | $^{13}C$-$^{13}C$ CORD, $^{13}C$ CP J-INADEQUATE | Kang et al., 2018[4] and Fernando et al., 2023[2] |
| | b | 104.2 | 55.0 | 73.4 | 83.1 | 75.9 | 60.5 | 174.3 | 23.1 | | |
| | c | 104.5 | 54.9 | 73.7 | 82.9 | 75.7 | 60.1 | 173.2 | 22.9 | | |
| | d | 104.0 | 55.3 | 73.6 | 83.6 | 75.8 | 60.6 | 174.7 | 22.9 | | |
| | e | 103.6 | 55.4 | 73.5 | 84.1 | 75.1 | 60.4 | 175.1 | 23.1 | | |
| chitosan | a | 102.1 | 55.7 | 72.7 | 79.7 | 75.2 | 62.1 | / | / | | |
| | b | 102.4 | 55.8 | 73.8 | 78.9 | 75.4 | 62.3 | / | / | | |
| β-1,3-glucan | a | 103.9 | 74.6 | 86.9 | 68.4 | 77.7 | 61.4 | / | / | | |
| | b | 103.4 | 74.4 | 85.7 | 68.9 | 77.5 | 61.2 | / | / | | |
| | c(m) | 103.1 | 73.8 | 84.5 | 69.0 | 77.0 | 61.9 | / | / | | |
| *T. marneffei*, 37 °C | | | | | | | | | | | |
| chitin | a | 103.3 | 55.6 | 73.2 | 84.5 | 75.5 | 60.3 | 175.3 | 23.1 | | |
| | b | 104.0 | 55.1 | 73.6 | 83.2 | 76.0 | 60.3 | 174.3 | 22.9 | | |
| | c | 104.4 | 54.8 | 73.5 | 82.9 | 75.8 | 60.9 | 173.2 | 23.1 | | |
| | e | 103.6 | 55.6 | 73.6 | 84.1 | 76.5 | 60.6 | 175.2 | 23.2 | | |
| chitosan | a | 102.2 | 55.8 | 73.5 | 79.4 | 75.2 | 62.4 | / | / | | |
| | b | 102.9 | 56.1 | 73.3 | 78.9 | 75.2 | 62.0 | / | / | | |
| | c(m) | 102.0 | 56.3 | 72.8 | 79.9 | 75.0 | 62.7 | / | / | | |
| β-1,3-glucan | a | 103.9 | 74.6 | 86.9 | 68.4 | 77.7 | 61.4 | / | / | | |
| | b | 103.4 | 74.6 | 85.7 | 69.2 | 78.4 | 61.1 | / | / | | |
| | c(m) | 103.1 | 73.9 | 84.8 | 68.7 | 76.8 | 62.2 | / | / | | |
| | d(m) | 103.8 | 74.6 | 88.0 | 68.6 | 77.2 | 61.7 | / | / | | |

**Supplementary table 10. chemical shifts of mobile polysaccharides in *T. marneffei* cell walls**. lower case letters are used to denote different allomorphs. Not applicable (/). Unidentified (-).

| Carbohydrate | | C1 | C2 | C3 | C4 | C5 | C6 | CO | $CH_3$ | Experiment | References |
|---|---|---|---|---|---|---|---|---|---|---|---|
| β-1,3-glucan | a | 103.9 | 74.7 | 86.7 | 68.4 | 77.9 | 61.6 | / | / | $^{13}$C DP J-refocused Inadequate | Chakraborty et al., 2021[1]; Fernando et al., 2023[2]; Kang et al., 2018[4] |
| | b | 103.3 | 74.1 | 84.5 | 68.8 | 76.6 | 61.6 | / | / | | |
| | c | 103.7 | 74.5 | - | 70.4 | 76.7 | 62.2 | / | / | | |
| α-1,2-Mannose | a | 99.4 | 80.2 | 72.9 | 69.9 | 73.5 | 61.6 | / | / | | |
| | b | 98.8 | 79.5 | 70.8 | 67.7 | 73.9 | 61.8 | / | / | | |
| | c | 101.3 | 79.3 | 70.7 | 67.2 | 72.6 | 61.2 | / | / | | |
| | d | 103.4 | 77.2 | - | - | - | - | / | / | | |
| α-1,6-Mannose | a | 104.2 | 71.5 | 74.0 | 67.6 | 71.2 | 63.7 | / | / | | |
| | b | 103.1 | 70.8 | 73.8 | 70.4 | 72.6 | 67.3 | / | / | | |
| Galactose | a | 101.1 | 77.1 | 74.7 | 82.1 | 73.5 | 62.9 | / | / | | |
| | b | 100.7 | 72.2 | 73.6 | 70.2 | 72.9 | 61.6 | / | / | | |
| | c | 97.9 | 72.2 | 73.6 | 70.2 | 72.9 | 61.6 | / | / | | |
| | d | 96.7 | 75.0 | - | - | - | - | / | / | | |
| | e | 99.6 | 72.3 | - | - | - | - | / | / | | |
| Arabinose | a | 106.2 | 87.7 | 75.6 | 82.1 | 69.8 | / | / | / | | |
| | b | 106.6 | 88.3 | 76.3 | 82.2 | 69.8 | / | / | / | | |
| | c | 107.4 | 88.1 | - | - | - | / | / | / | | |
| Galfuranose | a | 107.8 | 82.0 | 77.5 | 82.7 | 72.5 | 61.3 | / | / | | |
| | b | 108.5 | 81.5 | 77.4 | 83.3 | 70.1 | 62.2 | / | / | | |
| | c | 108.9 | 83.2 | 77.2 | 83.2 | 71.2 | 63.6 | / | / | | |
| Galactosamine | | 91.6 | 54.7 | - | - | - | - | / | / | | |
| N-acetyl-galactosamine | | 95.7 | 57.6 | - | - | - | - | / | / | | |
| Fucose | a(25°C) | 92.9 | 79.2 | 73.4 | 70 | 62.7 | 15.4 | / | / | | Furevi et al[5] |
| | b(25°C) | 92.0 | 80.6 | 69.7 | 75.2 | 60.7 | 16.1 | / | / | | |
| | 37°C | 92.4 | 77.8 | 73.4 | 72.0 | 58.5 | 17.8 | / | / | | |

**Supplementary table 11. $^{13}$C Chemical shifts of identified rigid amino acids in *T. Marneffei* cells.** Superscripts are used to represent the same type of amino acids (AAs). Not applicable (/). Unidentified (-). The rigid AAs are identified by 2D 53 ms $^{13}$C-$^{13}$C CORD (CORD) spectra.

| 25 °C | | | | | | | | |
|---|---|---|---|---|---|---|---|---|
| Amino acids | | Cα | Cβ | Cγ | Cδ | Cε | CO | CO' |
| Isoleucine | $I^a$ | 58.3 | 36.0 | 26.8(15.0) | 12.2 | / | 173.7 | / |
| | $I^b$ | 63.6 | 36.1 | 26.4(17.0) | 11.5 | / | 173.0 | / |
| | $I^c$ | 58.7 | 38.1 | 26.4(15.7) | 13.5 | / | - | / |
| | $I^d$ | 58.7 | 40.2 | 26.4(16.1) | 13.0 | / | - | / |
| Alanine | $A^a$ | 52.8 | 16.6 | / | / | / | 173.8 | / |
| | $A^b$ | 50.9 | 17.0 | / | / | / | 173.4 | / |
| | $A^c$ | 51.0 | 21.7 | / | / | / | 171.0 | / |
| | $A^d$ | 49.1 | 20.7 | / | / | / | 172.2 | / |
| Valine | $V^a$ | 59.0 | 36.4 | 19.7 | / | / | 173.1 | / |
| | $V^b$ | 58.4 | 36.7 | 19.5 | / | / | 172.0 | / |
| | $V^c$ | 64.8 | 29.1 | 19.6 | / | / | 172.0 | / |
| | $V^d$ | 63.8 | 29.2 | 19.5 | / | / | 173.8 | / |
| Lysine | $K^a$ | 56.2 | 34.0 | 25.2 | 30.0 | 40.5 | 174.4 | / |
| | $K^b$ | 57.8 | 33.2 | 27.3 | 30.9 | 40.1 | 174.3 | / |
| | $K^c$ | 56.4 | 32.0 | 29.0 | - | 41.4 | 175.6 | / |
| Tryptophan | $W^a$ | 53.3 | 29.2 | 130.1 | 145.1 | - | 173.6 | / |
| | $W^b$ | 57.6 | 30.6 | 130.1 | - | - | 173.8 | / |
| | $W^c$ | 59.6 | 33.6 | 130.2 | - | - | 173.9 | / |
| Phenylalanine | $F^a$ | 56.9 | 44.4 | 136.1 | - | - | 172.8 | / |
| | $F^b$ | 56.0 | 41.9 | 136.1 | / | / | 172.9 | / |
| | $F^c$ | 57.2 | 45.1 | 135.0 | - | - | 175.0 | / |
| | $F^d$ | 56.9 | 38.3 | 141.1 | - | - | 173.2 | / |
| | $F^e$ | 56.0 | 39.9 | 140.8 | - | - | 174.6 | / |
| | $F^f$ | 56.1 | 40.5 | 137.1 | - | - | 174.5 | / |
| Tyrosine | $Y^a$ | 58.1 | 39.9 | 128.5 | - | - | 174.5 | / |
| | $Y^b$ | 55.0 | 37.9 | 129.4 | - | - | 173.6 | / |
| Asparagine | $N^a$ | 53.7 | 39.9 | / | / | / | 174.4 | 177.5 |
| | $N^b$ | 52.6 | 40.1 | / | / | / | 174.5 | 177.4 |
| | $N^c$ | 53.4 | 38.4 | / | / | / | 172.9 | 178.2 |
| Aspartic | D | 55.8 | 39.7 | / | / | / | 172.7 | 175.1 |

| 37 °C | | | | | | | | |
|---|---|---|---|---|---|---|---|---|
| Amino acids | | Cα | Cβ | Cγ | Cδ | Cε | CO | CO' |
| Isoleucine | $I^a$ | 59.3 | 36.9 | 26.6(15.5) | 11.9 | / | 173.8 | / |
| | $I^b$ | 55.4 | 38.7 | 27.2(17.5) | 12.4 | / | 174.7 | / |
| Leucine | $L^b$ | 56.5 | 45.4 | 29.1 | 25.8 | / | 175.0 | / |
| Alanine | $A^a$ | 53.4 | 15.4 | / | / | / | 171.2 | / |
| | $A^b$ | 50.3 | 17.4 | / | / | / | 171.2 | / |
| | $A^c$ | 49.5 | 21.1 | / | / | / | 173.5 | / |
| | $A^d$ | 52.9 | 17.2 | / | / | / | 173.5 | / |
| | $A^e$ | 52.1 | 20.3 | / | / | / | 174.0 | / |
| Valine | $V^a$ | 60.0 | 36.7 | 18.6 | / | / | 173.5 | / |
| | $V^b$ | 59.2 | 36.7 | 19.9 | / | / | 174.5 | / |
| | $V^c$ | 64.0 | 30.1 | 20.5 | / | / | 176.2 | / |
| | $V^d$ | 64.4 | 29.9 | 22.5 | / | / | 175.8 | / |
| Histidine | H | 51.8 | 36.8 | 126.6 | - | - | 172.9 | / |
| Lysine | $K^a$ | 56.2 | 33.9 | 24.8 | 30.6 | 42.7 | 174.8 | / |
| | $K^b$ | 56.4 | 33.2 | 26.8 | 30.6 | 40.5 | 173.6 | / |
| Tryptophan | W | 53.1 | 29.7 | | | | 173.8 | / |
| Methionine | M | 56.2 | 29.8 | 18.4 | / | / | 170.9 | / |
| Phenylalanine | $F^a$ | 56.5 | 45.1 | 136.9 | - | - | 174.3 | / |
| | $F^b$ | 56.6 | 42.2 | 136.3 | - | - | 173.5 | / |
| | $F^c$ | 57.6 | 41.6 | 137.2 | - | - | 172.8 | / |
| | $F^d$ | 56.7 | 39.9 | 141.3 | - | - | 171.9 | / |
| | $F^e$ | 56.1 | 39.2 | 138.5 | - | - | 171.5 | / |
| | $F^f$ | 52.7 | 44.1 | 137.6 | - | - | 174.2 | / |
| Aspartic | $D^a$ | 51.7 | 43.6 | / | / | / | 170.8 | 174.0 |
| | $D^a$ | 52.8 | 39.4 | / | / | / | 171.7 | 175.5 |
| Asparagine | $N^a$ | 51.6 | 41.9 | / | / | / | 173.1 | 176.5 |
| | $N^b$ | 53.6 | 40.9 | / | / | / | 173.0 | 177.0 |
| Glutamic | $E^a$ | 57.0 | 30.0 | 36.6 | / | / | 173.2 | 178.2 |
| | $E^b$ | 52.6 | 29.6 | 37.6 | / | / | 172.9 | |
| Proline | $P^a$ | 64.3 | 30.2 | 25.7 | 52.3 | / | 172.9 | / |

| | | | | | | | | |
|---|---|---|---|---|---|---|---|---|
| Glutamic | E[a] | 51.4 | 29.4 | 38.3 | / | / | 171.2 | 173.9 |
| | E[b] | 52.8 | 29.9 | 35.4 | / | / | 173.6 | 176.4 |
| Serine | S[a] | 53.3 | 64.5 | / | / | / | 173.2 | / |
| | S[b] | 55.4 | 64.2 | / | / | / | 174.1 | / |
| Proline | P[a] | 64.1 | 29.9 | - | 50.5 | / | 174.0 | / |
| | P[b] | 64.1 | 29.2 | - | 47.6 | / | 170.8 | / |

| | | | | | | | | |
|---|---|---|---|---|---|---|---|---|
| | P[b] | 59.0 | - | - | 48.9 | / | 173.1 | / |
| | P[c] | 58.5 | - | - | 51.0 | / | - | / |
| Serine | S[a] | 55.0 | 64.7 | / | / | / | 173.7 | / |
| | S[b] | 57.1 | 64.1 | / | / | / | 173.1 | / |

Note: The identified asparagine and aspartic acids cannot be definitively confirmed due to their highly overlapping chemical shifts.

**Supplementary table 12. $^{13}$C Chemical shifts of identified mobile amino acids in *T. Marneffei* cells.** Superscripts are used to represent the same type of amino acids (AAs). Not applicable (/). Unidentified (-). The rigid AAs are identified by 2D 53 ms $^{13}$C-$^{13}$C CORD (CORD) spectra.

**25 ºC**

| Amino acids | | Cα | Cβ | Cγ | Cδ | Cε | CO | CO’ |
|---|---|---|---|---|---|---|---|---|
| Alanine | $A^a$ | 53.6 | 16.0 | / | / | / | 172.4 | / |
| | $A^b$ | 50.3 | 17.5 | / | / | / | 177.4 | / |
| | $A^c$ | 53.4 | 15.0 | / | / | / | 173.4 | / |
| | $A^d$ | 51.2 | 17.2 | / | / | / | 176.7 | / |
| | $A^e$ | 48.2 | 18.8 | / | / | / | 174.3 | / |
| | $A^f$ | 54.1 | - | / | / | / | 176.7 | / |
| Aspartic acid | D | 56.7 | 43.2 | / | / | / | 175.0 | 182.1 |
| Glutamic | E | 54.7 | 33.3 | 37.2 | / | / | 175.2 | 185.9 |
| Phenylalanine | $F^a$ | 55.5 | 43.2 | 136.7 | - | - | 175.3 | / |
| | $F^b$ | 52.9 | 47.0 | 131.4 | - | - | - | / |
| Glycine | $G^a$ | 45.2 | / | / | / | / | 171.5 | / |
| | $G^b$ | 42.4 | / | / | / | / | 173.1 | / |
| | $G^c$ | 42.9 | / | / | / | / | 172.0 | / |
| Histidine | $H^a$ | 55.5 | 28.2 | 129.5 | - | - | 175.3 | / |
| | $H^b$ | 54.9 | 26.9 | 127.8 | - | - | 175.6 | / |
| Leucine | L | 57.7 | 41.7 | 24.8 | 22.9 | | 173.7 | / |
| Asparagine | N | 52.4 | 35.6 | - | - | - | 174.0 | 177.5 |
| Arginine | R | 56.1 | 29.9 | 27.2 | 40.5 | / | 175.2 | / |
| Proline | P | - | 32.5 | 30.2 | 50.6 | - | - | - |
| Valine | $V^a$ | 65.4 | 31.6 | 23.5 | / | / | 170.0 | / |
| | $V^b$ | 64.1 | 30.8 | 18.6 | / | / | - | / |
| | $V^c$ | 64.1 | 27.6 | - | / | / | - | / |
| | $V^d$ | 57.0 | 30.8 | 19.4 | / | / | 173.6 | / |
| | $V^e$ | 58.8 | 30.8 | 19.4 | / | / | - | / |
| | $V^f$ | 59.4 | 30.8 | - | / | / | 172.3 | / |
| | $V^g$ | 61.4 | 29.9 | - | / | / | 175.4 | / |
| | $V^h$ | 64.5 | 28.8 | - | / | / | - | / |
| | $V^i$ | 57.9 | 33.7 | 19.9 | / | / | - | / |
| Isoleucine | $I^a$ | 62.4 | 36.9 | - | - | - | 176.1 | |

**37 ºC**

| Amino acids | | Cα | Cβ | Cγ | Cδ | Cε | CO | CO’ |
|---|---|---|---|---|---|---|---|---|
| Alanine | A | 51.6 | 17.1 | / | / | / | 176.6 | / |
| Phenylalanine | $F^a$ | 57.4 | 37.4 | 135.2 | - | - | 175.2 | / |
| | $F^b$ | 57.2 | 39.3 | - | - | - | 175.2 | / |
| Glycine | $G^a$ | 45.2 | / | / | / | / | 171.5 | / |
| | $G^b$ | 42.4 | / | / | / | / | 173.4 | / |
| | $G^c$ | 43.3 | / | / | / | / | 172.0 | / |
| Histidine | H | 55.4 | 27.6 | 128.3 | - | - | 175.7 | / |
| Leucine | $L^a$ | 54.5 | 40.6 | 25.1 | 23.4 | | 176.8 | / |
| | $L^b$ | 54.8 | 40.3 | 27.4 | 23.0 | | 173.9 | |
| Methionine | M | 54.9 | 33.3 | 30.5 | / | / | 175.7 | / |
| Asparagine | $N^a$ | 53.6 | 37.9 | / | / | / | 170.2 | 176.9 |
| | $N^b$ | 52.7 | 27.8 | / | / | / | - | - |
| Proline | P | 60.4 | 33.0 | 24.8 | 46.5 | / | 175.2 | / |
| Valine | $V^a$ | 61.5 | 29.7 | 17.6 | / | / | 173.6/175.2 | / |
| | $V^b$ | 62.3 | 29.7 | 18.5 | / | / | 175.7 | / |
| | $V^c$ | 61.1 | 30.1 | 16.8 | / | / | 175.1 | / |
| Tryptophan | W | 58.5 | 29.8 | 127.5 | - | - | - | / |
| Isoleucine | I | 60.7 | 36.8 | - | - | - | 175.4 | / |

| | | | | | | |
|---|---|---|---|---|---|---|
| I[b] | 60.3 | 39.1 | - | - | - | 175.2 |

**Supplementary table S13. Intermolecular cross peaks in *T. marnefei.*** The chemical shifts for the two dimensions of the spectra ($\omega_1$ and $\omega_2$). Asterisks indicate interactions where only one peak is observed along the diagonal line with the absence of the other. The bold font indicates that these interactions occur between this carbon site with the other ones, i.e., Cs$^{a}$4-Ch$^{d}$1/**B$^{a}$1**, represents the interaction Cs$^{a}$4-B$^{a}$1.

| **25 °C** | | | **37 °C** | | |
|---|---|---|---|---|---|
| Interaction | $\omega_1$ or $\omega_2$ (ppm) | $\omega_1$ or $\omega_2$ (ppm) | Interaction | $\omega_1$ or $\omega_2$ (ppm) | $\omega_1$ or $\omega_2$ (ppm) |
| Chitin-chitosan | | | Chitin-chitosan | | |
| *Ch$^{c}$1-Cs$^{b}$1 | 104.5 | 102.4 | *Cs$^{c}$1-B$^{b}$1/**Ch$^{a}$1** | 102.0 | 103.3 |
| Ch$^{d}$4-Cs$^{b}$1 | 83.6 | 102.4 | Cs$^{a/c}$1-B$^{a/d}$1/**Ch$^{b}$1** | 102.0/102.2 | 104.0 |
| Ch$^{c}$4-Cs$^{a}$4 | 84.1 | 79.7 | Cs$^{b}$1-B$^{a/d}$1/**Ch$^{b}$1** | 102.9 | 104.0 |
| Cs$^{a}$4-**Ch$^{d}$1**/B$^{a}$1 | 79.7 | 104.0 | Cs$^{a}$4-B$^{c}$1/**Ch$^{a}$1** | 79.4 | 103.3 |
| Cs$^{b}$4-**Ch$^{d}$1**/B$^{a}$1 | 78.9 | 104.0 | Cs$^{b}$4-B$^{c}$1/**Ch$^{a}$1** | 78.9 | 103.3 |
| Cs2-Ch$^{b/c}$1 | 55.8 | 104.2 | Cs$^{a/c}$5/**Ch$^{a}$5**-Cs$^{b}$1 | 75.5 | 102.9 |
| Ch$^{a/b/d}$6-Cs$^{b}$4 | 60.5 | 78.9 | Ch$^{e}$4-Cs$^{b}$1 | 84.1 | 102.9 |
| Cs6-Ch$^{d}$4 | 62.1 | 83.6 | Ch$^{b/c}$4-Cs$^{b}$1 | 83.2 | 102.9 |
| Cs$^{a}$3-Ch$^{b/c}$4 | 72.7 | 83.1 | **Ch$^{b/c/e}$5**/Cs$^{a}$5-Cs$^{a/c}$1 | 76.0 | 102.2 |
| Ch2-Cs6 | 55.0 | 62.1 | **Ch3**/Cs3-Cs$^{a/c}$1 | 73.5 | 102.2 |
| Cs$^{a}$1-ChMe | 102.1 | 22.9 | **Ch$^{b}$1**/B$^{a/d}$1-Cs$^{c}$6 | 104.0 | 62.7 |
| Cs$^{b}$1-ChMe | 102.4 | 22.9 | Ch6-Cs$^{b}$1 | 60.6 | 102.2 |
| Cs$^{a}$4-ChMe | 79.7 | 22.9 | Ch6-Cs$^{a/c}$1 | 60.6 | 102.2 |
| Ch$^{c/d}$5/**Cs$^{b}$5**-ChMe | 75.4 | 22.9 | Ch$^{b/c}$2-**Cs$^{b}$1**/B$^{c}$1 | 104.4 | 102.9 |
| Cs$^{a}$5-ChMe | 75.2 | 22.9 | **Cs$^{a/b}$6**/B$^{c/d}$6-Ch$^{b/c}$4 | 62.0 | 82.9 |
| Cs1-Ch$^{a/e}$CO | 102.1 | 175.1 | Cs$^{b/c}$2-**Ch$^{a}$5**/Cs5 | 56.3 | 75.5 |
| **Cs5**/Ch$^{e}$5-Ch$^{c}$CO | 75.2 | 173.2 | Ch$^{c}$2-Cs$^{c}$6 | 54.8 | 62.7 |
| Cs5-Ch$^{e}$CO | 75.2 | 175.1 | **Ch$^{a/e}$2**/Cs$^{a}$2-Cs$^{c}$6 | 103.3 | 62.7 |
| **Ch$^{a}$4**/B$^{c}$3-Cs2 | 84.5 | 55.8 | Cs$^{a}$2-Ch$^{e}$6 | 55.8 | 103.6 |
| Cs$^{a}$1-B$^{a}$1/**Ch$^{d}$1** | 102.1 | 104.0 | Ch6-Cs$^{c}$4 | 60.6 | 79.9 |
| Cs$^{b}$1-B$^{a}$1/**Ch$^{d}$1** | 102.4 | 104.0 | **Cs6**/B$^{c}$6-**Ch3**/Cs$^{a/b}$3 | 62.4 | 73.5 |
| Cs6-**Ch$^{d}$1**/B$^{a}$1 | 62.1 | 104.0 | ChMe-**Cs$^{b}$1**/B$^{c}$1 | 22.9 | 102.9 |
| Ch$^{b/c}$1-Cs6 | 104.2 | 62.1 | ChMe-Cs$^{a/c}$1 | 22.9 | 102.2 |
| Chitin-chitin | | | ChMe-Ch$^{a}$5/**Cs5** | 23.1 | 75.2 |
| **Ch$^{a}$1**/B$^{c}$1-Ch$^{c}$1 | 103.1 | 104.5 | ChMe-Cs$^{a/c}$6 | 23.1 | 62.4 |
| **Ch$^{d}$1**/B$^{a}$1-Ch$^{c}$1 | 104.0 | 104.5 | ChMe-**Cs$^{b}$6**/B$^{c/d}$6 | 23.1 | 62.0 |
| Ch$^{a}$5-**Ch$^{d}$1**/B$^{a}$1 | 76.5 | 104.0 | ChMe-Cs2 | 23.1 | 56.1 |
| **Ch$^{a}$4**/B$^{c}$3-**Ch$^{d}$1**/B$^{a}$1 | 84.5 | 104.0 | Cs5-Ch$^{a/e}$CO | 75.2 | 175.3 |
| Ch$^{b/c}$4-**Ch$^{a}$1**/B$^{c}$1 | 83.1 | 103.1 | Ch3/**Cs3**-Ch$^{a/e}$CO | 73.5 | 175.3 |
| Ch$^{b/c}$4-Ch$^{a}$4 | 75.9 | 76.5 | Ch3/**Cs3**-Ch$^{b}$CO | 73.5 | 174.3 |
| **Ch$^{a}$4**/B$^{c}$3-ChMe | 76.5 | 23.1 | Chitin-chitin | | |
| Ch$^{b}$5-ChMe | 75.7 | 22.9 | Ch$^{b}$4-Ch$^{a}$1 | 83.2 | 84.5 |
| **Ch$^{c/d}$5**/Cs$^{b}$5-ChMe | 75.7 | 22.9 | Ch$^{a/c}$5-**Ch$^{b}$1**/B$^{a/d}$1 | 75.5 | 103.3 |
| Ch$^{c}$1-Ch$^{a/e}$CO | 104.5 | 175.1 | Ch6-**Ch$^{b}$1**/B$^{a/d}$1 | 60.6 | 103.3 |
| **Ch$^{e}$1**/B$^{b}$1-Ch$^{b}$CO | 103.6 | 174.3 | Ch$^{b/c}$2-Ch$^{a}$4 | 54.8 | 84.5 |
| Ch$^{d}$4-Ch$^{c}$CO | 83.6 | 173.2 | **Ch$^{c}$6**/B$^{a/b}$6-Ch$^{a/e}$4 | 60.9 | 84.1 |
| Ch$^{e}$4-Ch$^{b}$CO | 82.9 | 174.3 | Ch$^{b/c}$4-Ch$^{a}$4 | 82.9 | 84.5 |
| Cs5/**Ch$^{e}$5**-Ch$^{c}$CO | 75.1 | 173.2 | ChMe-Ch$^{a}$4 | 23.1 | 84.5 |
| Ch$^{c}$2-**Ch$^{a}$4**/B$^{c}$3 | 54.9 | 84.5 | ChMe-Ch$^{e}$4 | 23.1 | 84.1 |
| Chitosan-chitosan | | | ChMe-Ch$^{c}$4 | 23.1 | 82.9 |
| Cs$^{a}$1-Cs$^{b}$1 | 102.1 | 102.4 | ChMe-Ch$^{a/b}$5 | 23.1 | 75.5 |
| Cs$^{b}$4-Cs$^{a}$1 | 78.9 | 102.1 | ChMe-Ch$^{e}$5 | 23.1 | 76.5 |
| Cs$^{b}$5-Cs$^{a}$1 | 75.4 | 102.1 | ChMe-**Ch$^{a}$5**/Cs5 | 23.1 | 75.5 |
| Cs$^{a}$4-**Cs$^{b}$3**/B2 | 79.7 | 73.8 | ChMe-Ch6 | 23.1 | 60.6 |
| β-1,3-glucan-chitin | | | Ch$^{c}$1-Ch$^{a/e}$CO | 104.4 | 175.3 |
| Ch$^{a}$1/**B$^{c}$1**-Ch$^{c}$1 | 103.1 | 104.5 | Ch$^{b}$1-Ch$^{a/e}$CO | 104.0 | 175.3 |
| **Ch$^{a}$1**/B$^{c}$1-B$^{a}$1 | 103.1 | 103.9 | **Ch$^{a/e}$1**/B$^{b}$1-Ch$^{a/e}$CO | 103.3/103.6 | 175.3 |
| Ch$^{e}$1/**B$^{a}$1**-Ch$^{c}$1 | 103.9 | 104.5 | Ch$^{b}$4-Ch$^{c}$CO | 83.2 | 173.2 |
| B$^{a}$3-**Ch$^{a}$1**/B$^{c}$1 | 86.9 | 103.1 | Ch$^{c}$4-Ch$^{b}$CO | 82.9 | 174.3 |
| B$^{a}$3-Ch$^{b/d}$1 | 86.9 | 104.2 | Ch$^{a/e}$4-Ch$^{a/e}$CO | 84.5/84.1 | 175.3 |
| B$^{a}$3-Ch$^{c}$1 | 86.9 | 104.5 | Ch$^{b/c}$4-Ch$^{a/e}$CO | 83.2 | 175.3 |
| Ch$^{a}$5-Ch$^{d}$1/**B$^{a}$1** | 76.5 | 103.9 | Ch$^{a/b}$5-Ch$^{c}$CO | 75.5/76.0 | 173.2 |
| **Ch$^{a}$4**/B$^{c}$3-Ch$^{d}$1/**B$^{a}$1** | 84.5 | 103.9 | Ch$^{a/b/c}$5-Ch$^{a/e}$CO | 75.8 | 175.3 |
| Ch$^{b/c}$4-Ch$^{a}$1/**B$^{c}$1** | 83.1 | 103.1 | **Ch3**/Cs3-Ch$^{a/e}$CO | 73.2/73.6 | 175.3 |
| B$^{a}$5-**Ch$^{a}$1**/B$^{c}$1 | 77.7 | 103.9 | **Ch3**/Cs3-Ch$^{b}$CO | 73.2/73.6 | 174.3 |
| Ch$^{b/c/d}$5-B$^{a}$1 | 75.8 | 103.9 | Ch$^{a/b/e}$6-Ch$^{a/e}$CO | 60.6 | 175.3 |
| B$^{a}$4-**Ch$^{a}$1**/B$^{c}$1 | 68.4 | 84.5 | Ch$^{c/e}$6-Ch$^{a/e}$CO | 60.9 | 175.3 |
| B6-Ch$^{c}$1 | 61.9 | 104.5 | Chitosan-chitosan | | |

| | | |
|---|---|---|
| B6-**Ch$^{a}$1**/B$^{c}$1 | 61.9 | 103.1 |
| B$^{a}$4-Ch$^{d}$4 | 68.4 | 83.6 |
| Ch$^{a/b/e}$3-B$^{a}$3 | 73.6 | 86.9 |
| **Ch$^{c}$3**/Cs$^{a/b}$3/B$^{c}$2-B$^{a}$3 | 73.7 | 86.9 |
| Ch$^{b/c/d}$5-B$^{a}$3 | 75.8 | 86.9 |
| B$^{c}$5-Ch$^{e}$4 | 77.0 | 82.9 |
| B$^{b}$3-Ch$^{d}$4 | 85.7 | 83.6 |
| Ch$^{a}$3-B$^{a}$3 | 73.5 | 86.9 |
| B5-**Ch3**/Cs3 | 77.5 | 73.5 |
| B4-Cs3/**Ch3** | 68.4 | 73.5 |
| Ch$^{b/d}$2-B$^{c}$5 | 55.3 | 77.0 |
| Ch2-B6 | 55.3 | 61.4 |
| Ch$^{a}$4/**B$^{c}$3**-ChMe | 84.5 | 22.9 |
| B5-ChMe | 77.5 | 22.9 |
| B$^{a/b}$2-ChMe | 74.4 | 22.9 |
| B6-ChMe | 61.4 | 22.9 |
| Ch$^{e}$1/**B$^{b}$1**-Ch$^{b}$CO | 103.4 | 174.3 |
| B$^{a/b}$2-Ch$^{a/d}$CO | 74.4 | 174.4 |
| Ch$^{a}$5-B$^{a}$3 | 76.5 | 86.9 |
| Ch$^{c}$2-Ch$^{a}$4/**B$^{c}$3** | 54.9 | 84.5 |
| Ch$^{e}$4-B$^{a/b}$6 | 84.1 | 61.2 |
| B$^{a}$3-Ch6 | 86.9 | 60.5 |
| β-1,3-glucan-chitosan | | |
| Cs$^{a}$1-**B$^{a}$1**/Ch$^{d}$1 | 102.1 | 103.9 |
| Cs$^{b}$1-**B$^{a}$1**/Ch$^{d}$1 | 102.4 | 103.9 |
| B$^{a}$3-Cs$^{b}$1 | 86.9 | 102.4 |
| B$^{a}$3-Cs$^{a}$1 | 86.9 | 102.1 |
| Cs$^{a}$4-Ch$^{d}$1/**B$^{a}$1** | 79.7 | 103.9 |
| Cs$^{b}$4-Ch$^{d}$1/**B$^{a}$1** | 78.9 | 103.9 |
| B$^{a}$5-Cs$^{b}$1 | 77.7 | 102.4 |
| B$^{a}$4-Cs1 | 68.4 | 102.1 |
| B6-Cs1 | 61.4 | 102.1 |
| B6-Cs$^{b}$4 | 61.4 | 78.9 |
| B$^{a}$4-Cs$^{b}$4 | 68.4 | 78.9 |
| B$^{a}$4-Cs$^{a}$4 | 68.4 | 79.7 |
| Ch$^{c}$3/**Cs$^{a/b}$3**/B$^{c}$2-B$^{a}$3 | 72.7/73.8 | 86.9 |
| Cs$^{b}$4-B$^{a}$3 | 78.9 | 86.9 |
| B5-Ch3/**Cs3** | 77.5 | 72.7 |
| Cs6-B$^{a/b}$2 | 62.1 | 74.6 |
| B4-**Cs3**/Ch3 | 68.4/68.9 | 72.7/73.8 |
| B6-Cs6 | 61.4 | 62.1 |
| B$^{a}$4-Cs6 | 68.4 | 62.1 |
| Ch$^{a}$4/**B$^{c}$3**-Cs2 | 84.5 | 55.7 |
| B5-Cs$^{b}$4 | 77.5 | 78.9 |
| B5-Cs$^{a}$4 | 77.5 | 79.7 |
| Cs6-Ch$^{d}$1/**B$^{a}$1** | 62.1 | 103.9 |
| Cs6-B$^{a}$3 | 62.1 | 86.9 |
| β-1,3-glucan- β-1,3-glucan | | |
| Ch$^{a}$1/**B$^{c}$1**-B$^{a}$1 | 84.5 | 86.9 |
| B$^{a}$3-Ch$^{a}$1/**B$^{c}$1** | 86.9 | 103.1 |
| Ch$^{a}$4/**B$^{c}$3**-Ch$^{d}$1/**B$^{a}$1** | 84.5 | 103.9 |
| B$^{a}$5-Ch$^{a}$1/**B$^{c}$1** | 77.7 | 103.1 |
| B$^{a}$4-Ch$^{a}$1/**B$^{c}$1** | 68.4 | 103.1 |
| B6-Ch$^{a}$1/**B$^{c}$1** | 61.4 | 103.1 |
| B$^{a/b}$6-B$^{c}$4 | 61.4 | 69.0 |
| Ch$^{c}$3/Cs$^{a/b}$3/**B$^{c}$2**-B$^{a}$3 | 74.4 | 86.9 |
| B$^{a}$5-B$^{b}$3 | 77.7 | 85.7 |
| B$^{b}$3-B$^{a}$3 | 85.7 | 86.9 |
| B$^{a/c}$6-B$^{b}$3 | 61.9 | 85.7 |
| B$^{a/c}$6-B$^{b}$4 | 61.9 | 68.9 |
| B$^{c}$4-B$^{a}$3 | 68.9 | 86.9 |
| Carbohydrate-protein | | |
| *K$^{a}$β-B$^{a}$3 | 34.0 | 86.9 |
| K$^{a}$β-Cs5/Ch$^{e}$5 | 34.0 | 75.2 |
| K$^{a}$β-Ch$^{a/b/d}$5 | 34.0 | 76.5/75.8 |
| Ch3/**Cs3**-Cs$^{a/c}$1 | 72.8/73.3 | 102.2 |
| Ch$^{b/c/e}$5/**Cs$^{a}$5**-Cs$^{a/c}$1 | 75.2 | 102.2 |
| Cs$^{a/c}$5/**Ch$^{a}$5**-Cs$^{b}$1 | 75.5 | 102.9 |
| *Cs$^{c}$1-Cs$^{a}$4 | 102.0 | 79.4 |
| Cs$^{b}$3-Cs$^{a/c}$1 | 73.3 | 102.2 |
| Cs$^{b/c}$2-Ch$^{a}$5/**Cs5** | 56.3 | 75.2 |
| **Cs6**/B$^{c}$6-Ch3/**Cs$^{a/b}$3** | 62.4 | 73.5 |
| β-1,3-glucan-chitin | | |
| B$^{c}$1-B$^{a/d}$1/**Ch$^{b}$1** | 103.1 | 104.0 |
| B$^{b/c}$1/Ch$^{a}$1-**Ch$^{c}$1** | 103.4 | 104.4 |
| B$^{b}$2-**Ch$^{e}$1**/B$^{a}$1 | 74.6 | 103.6 |
| B$^{c}$3-**Ch$^{e}$1**/B$^{a}$1 | 84.8 | 103.6 |
| Ch$^{b/c}$4-B$^{c}$1 | 82.9 | 103.1 |
| B$^{c}$1-**Ch$^{e}$5**/B$^{c}$5 | 103.1 | 76.5 |
| Ch$^{a/e}$5-Ch$^{b}$1/**B$^{a/d}$1** | 75.5 | 103.9 |
| B4-Ch$^{c}$1 | 68.7 | 104.4 |
| Ch6-Ch$^{b}$1/**B$^{a/d}$1** | 60.6 | 103.8 |
| *B$^{b}$4-Ch$^{a/e}$CO | 69.2 | 175.3 |
| Ch$^{b/c}$2-Cs$^{b}$1/**B$^{c}$1** | 54.8 | 103.1 |
| Cs$^{a/b}$6/**B$^{c/d}$6**-Ch$^{b/c}$4 | 62.2 | 82.9 |
| Ch$^{c}$6/**B$^{a/b}$6**-Ch$^{a/e}$4 | 61.1 | 84.5 |
| Ch$^{a/b}$6-B$^{a}$3 | 60.3 | 86.9 |
| B$^{a}$3-Ch$^{e}$4 | 86.9 | 84.1 |
| **Ch3**/Cs$^{a/c}$3-B$^{a}$3 | 73.2/73.5 | 86.9 |
| Ch$^{a/b/c}$5-B$^{a}$3 | 75.5/76.0 | 86.9 |
| Ch$^{b/c}$4-B$^{a}$3 | 83.2 | 86.9 |
| Ch$^{e}$5-B$^{a}$3 | 76.5 | 86.9 |
| Ch2-B5 | 55.1/55.6 | 77.7/78.4 |
| Ch2-B6 | 55.1/55.6 | 61.7 |
| Ch6-B$^{a/c/d}$5 | 60.6 | 77.2 |
| Cs6/**B$^{c}$6-Ch3**/Cs$^{a/b}$3 | 62.2 | 73.5 |
| Ch$^{a/b}$6-B$^{a/c/d}$4 | 60.3 | 68.6 |
| Ch$^{c}$6-B$^{b}$4 | 60.9 | 69.2 |
| Ch3-B$^{a/b/d}$5 | 73.2/73.6 | 77.2/77.7 |
| ChMe-Cs$^{b}$1/**B$^{c}$1** | 23.1 | 103.1 |
| ChMe-B$^{a/c/d}$5 | 23.1 | 77.2/77.7 |
| ChMe-B$^{a/b/d}$2 | 23.1 | 74.6 |
| Ch$^{a/e}$1/**B$^{b}$1**-Ch$^{a/e}$CO | 103.4 | 175.3 |
| B2-Ch$^{a/e}$CO | 74.6/73.9 | 175.3 |
| β-1,3-glucan-chitosan | | |
| *Cs$^{a/c}$1-Ch$^{a}$1/**B$^{b}$1** | 102.2 | 103.4 |
| Cs$^{a/c}$1-**B$^{a/d}$1**/Ch$^{b}$1 | 102.2 | 103.9 |
| Cs$^{b}$1-**B$^{a/d}$1**/Ch$^{b}$1 | 102.9 | 103.9 |
| B$^{a}$3-Cs$^{b}$1 | 86.9 | 102.9 |
| B$^{a/d}$5-Cs$^{b}$1 | 77.2/77.7 | 102.9 |
| B$^{a}$5-Cs$^{a/c}$1 | 77.7 | 102.2 |
| B$^{a/b/d}$2-Cs$^{b}$1 | 74.6 | 102.9 |
| B$^{a/b/d}$2-Cs$^{a/c}$1 | 74.6 | 102.2 |
| B4-**Cs$^{b}$1**/B$^{c}$1 | 68.7 | 102.9 |
| B4-Cs$^{a/c}$1 | 68.7 | 102.2 |
| Ch$^{b}$1/**B$^{a/d}$1**-Cs$^{c}$6 | 103.9 | 62.7 |
| B6-Cs$^{a/c}$1 | 61.7 | 102.2 |
| **Cs6**/B$^{c/d}$6-B$^{a}$3 | 62.4 | 86.9 |
| B$^{a/d}$4-Cs$^{b}$4 | 68.4 | 78.9 |
| B$^{a/d}$4-Cs$^{a}$4 | 68.4 | 79.4 |
| Cs$^{b}$3-B$^{a}$3 | 73.3 | 86.9 |
| Ch3/**Cs$^{a/c}$3**-B$^{a}$3 | 73.5 | 86.9 |
| Cs$^{a}$4-B$^{a}$3 | 79.4 | 86.9 |
| Cs$^{b}$4-B$^{a}$3 | 78.9 | 86.9 |
| Cs6/**B$^{c}$6**-Ch3/**Cs$^{a/b}$3** | 62.2 | 73.5 |
| B$^{a/b/d}$6-Cs$^{a/b}$4 | 61.4 | 79.9 |
| Cs$^{c}$3-B$^{a/d}$5 | 72.8 | 77.2/77.7 |
| Cs$^{c}$3-B$^{a/b/d}$2 | 72.8 | 74.6 |
| B$^{c}$2-Cs$^{c}$4 | 73.9 | 79.9 |
| Cs$^{b}$4-B$^{a/d}$5 | 78.9 | 77.2/77.7 |

| | | |
|---|---|---|
| W$^{c}$β-Cs$^{b}$3 | 33.6 | 73.8 |
| K$^{a/b/c}$β/W$^{c}$β-B4 | 32.0/34.0/33.6 | 68.4/68.9 |

| | | |
|---|---|---|
| B$^{a}$5-Cs$^{a}$4 | 77.7 | 79.4 |
| β-1,3-glucan- β-1,3-glucan | | |
| B$^{a/d}$6-B$^{b}$3 | 61.7 | 85.7 |
| Cs6/**B$^{c/d}$6**-B$^{a}$3 | 62.2 | 86.9 |
| B$^{a}$4-B$^{b}$3 | 68.4 | 85.7 |
| B$^{b}$4-B$^{a}$3 | 69.2 | 86.9 |
| B$^{a/b/d}$2-B$^{c}$3 | 74.6 | 84.8 |
| B$^{a}$5-B$^{d}$3 | 77.7 | 88.0 |
| B$^{a}$5-B$^{b}$3 | 77.7 | 85.7 |
| *B$^{c}$3-B$^{a}$3 | 84.8 | 86.9 |
| B$^{b}$3-B$^{a}$3 | 85.7 | 86.9 |
| B$^{d}$3-B$^{a}$3 | 88.0 | 86.9 |
| B$^{b}$4-B$^{a/d}$5 | 69.2 | 77.2/77.7 |
| B$^{b}$4-B$^{a/d}$2 | 69.2 | 74.6 |
| Carbohydrate-protein | | |
| K$^{a}$β/Wβ-B4 | 33.9/29.7 | 68.6 |
